# Structural and thermodynamic consequences of base pairs containing pseudouridine and N1-methylpseudouridine in RNA duplexes

**DOI:** 10.1101/2023.03.19.533340

**Authors:** Nivedita Dutta, Indrajit Deb, Joanna Sarzynska, Ansuman Lahiri

**Affiliations:** Department of Biophysics, Molecular Biology and Bioinformatics, University of Calcutta, 92, Acharya PrafullaChandra Road, Kolkata 700009, West Bengal, India; Institute of Bioorganic Chemistry, Polish Academy of Sciences, Noskowskiego 12/14, 61-704 Poznan, Poland

**Keywords:** Pseudouridine, N1-methylpseudouridine, RNA duplex, mRNA therapeutics, Structure and thermodynamics

## Abstract

Pseudouridine (Ψ) is one of the most common post-transcriptional modifications in RNA and has been known to play significant roles in several crucial biological processes. The N1-methyl derivative of pseudouridine i.e N1-methylpseudouridine has also been reported to be important for the stability and function of RNA. Several studies suggest the importance of pseudouridine and N1-methylpseudouridine in mRNA therapeutics. The critical contribution of pseudouridine, especially that of its N1-methyl derivative in the efficiency of the COVID-19 mRNA vaccines, suggests the requirement to better understand the role of these modifications in the structure, stability and function of RNA. In the present study, we have investigated the consequences of the presence of these modifications in the stability of RNA duplex structures by analyzing different structural properties, hydration characteristics and energetics of these duplexes. We have previously studied the structural and thermodynamic properties of RNA duplexes with an internal Ψ-A pair and reported the stabilizing effect of Ψ over U (Deb, I. et al. *Sci Rep* 9, 16278 (2019)). Here, we have extended our work to understand the properties of RNA duplexes with an internal m^1^Ψ-A pair and also theoretically demonstrate the effect of substituting internal U-G, U-U and U-C mismatches with the Ψ-G, Ψ-U and Ψ-C mismatches and also with the m^1^Ψ-G, m^1^Ψ-U and m^1^Ψ-C mismatches respectively, within dsRNA. Our results indicate the context-dependent stabilization of base stacking interactions by N1-methylpseudouridine compared to uridine and pseudouridine, presumably resulting from the increased molecular polarizability due to the presence of the methyl group.

## INTRODUCTION

Pseudouridine (Ψ), an isomer of uridine, is the most abundant RNA modification and has been reported to substantially contribute to the structural, dynamic and thermal stability of RNA ^1–7^. This modification is frequently observed in different types of RNA, e.g. tRNA, rRNA, tmRNA, snRNA, snoRNA and is also found in mRNA and lncRNA ^8–13^. The formation of Ψ from U is catalyzed by a class of highly conserved enzymes collectively known as the pseudouridine synthases ^14,15^. Pseudouridine contains a C-C (C1’-C5) glycosidic bond, instead of the C-N (C1’-N1) glycosidic bond in uridine and hence it contains an additional ring nitrogen atom (N1 imino atom) (Figure 1) that acts as an additional H-bond donor ^1,2,6^.

**Figure 1.**
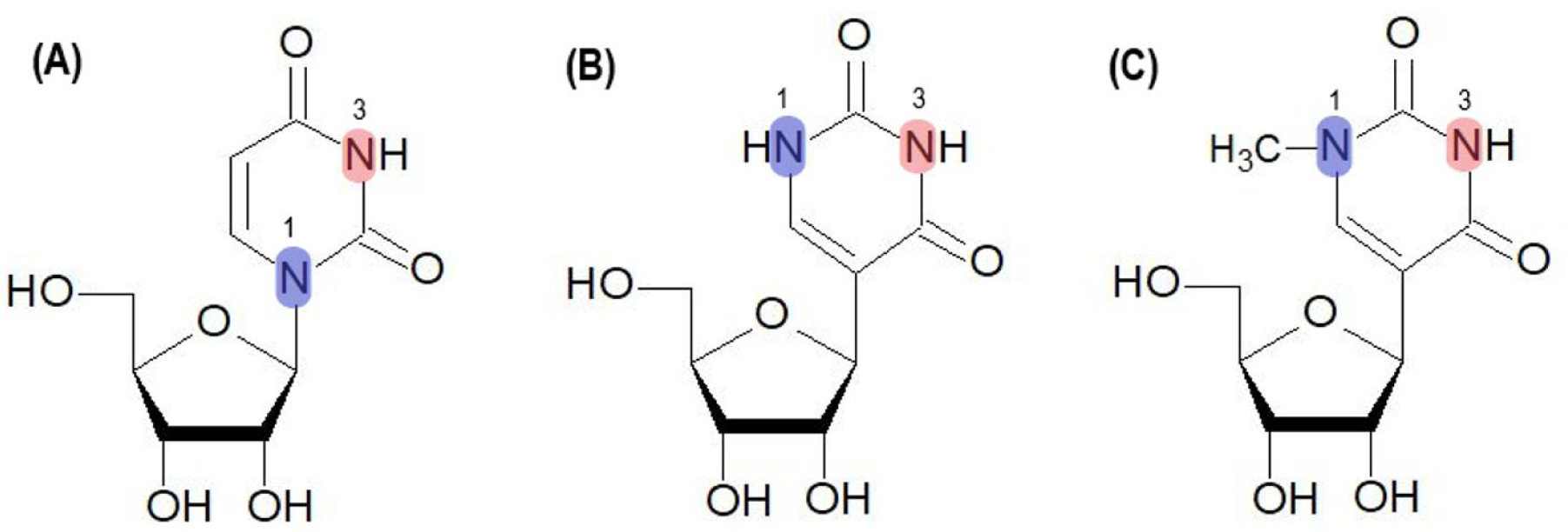
Structures of (A) uridine (U), (B) pseudouridine (Ψ) and N1-methylpseudouridine (m^1^Ψ).

The N1-methyl derivative of this modification, known as N1-methylpseudouridine or 1-methylpseudouridine (m^1^Ψ), initially identified as a metabolite in the *Streptomyces platensis* bacteria and later observed in the TΨC loop of archaeal tRNAs in place of ribothymidine (rT), has similar base pairing properties as that of uridine (U) or ribothymidine (rT) nucleobases ^16,17^. The ribosome assembly factor *Nep1* from *Saccharomyces cerevisiae* and *Methanocaldococcus jannaschii* has been reported to be a SAM-dependent pseudouridine-N1-specific methyltransferase ^18^. Chatterjee et al. (2012) reported that TrmY (tRNA (pseudouridine(54)-N(1))-methyltransferase) which is a SPOUT-type, SAM-dependent methyltransferase is responsible for the methylation of Ψ54 into m^1^Ψ54 in the tRNA of certain archaebacteria ^19^. Despite the absence of additional hydrogen bond donor due to the presence of methyl (-CH3) group attached with N1 (Figure 1), m^1^Ψ has also been reported to contribute to the stability of RNA structures ^6,20–22^.

Both pseudouridine and N1-methylpseudouridine have been found to play diverse and significant roles in several biological processes, e.g. stability and function of RNA, translation, RNA-protein interactions, innate immunity, etc ^23–28^. These modifications, especially m^1^Ψ have been reported to be extremely important in the efficiency of therapeutic mRNAs and mRNA vaccines as the incorporation of these modifications results in mRNA-transcript stability, enhanced translation efficiency and reduced immunogenicity ^27,29–34^. Interestingly, the FDA-approved COVID-19 mRNA vaccines developed by Pfizer-BioNtech and Moderna, substitute every U residue with m^1^Ψ ^32^.

Pseudouridine, also known as the universal base, can form more stable pairs with all the four canonical bases compared to those formed by U within a double helix ^3–4^. Experimental and theoretical studies have suggested that the substitution of U with Ψ results in the stabilization of the A-RNA conformation within ssRNA and dsRNA oligomers by the enhancement of base-stacking interactions and the presence of the additional hydration site also contributes to the stability ^3–6,35–37^. In their thermodynamic study, Hudson et al (2013) reported the nearest-neighbor (NN) thermodynamic parameters for the Ψ-A and suggested that the stabilization of the duplexes containing the Ψ-A pair largely depends on the sequence context ^3^. Another thermodynamic study by Kierzek et al (2014), revealed that substitution of U with Ψ results in stabilization of RNA duplexes depending on the position of Ψ within the dsRNA, base pair type, and the type and orientation of the neighboring WC pairs ^4^. A recent study which also reports the nearest neighbor parameters for Ψ-A and m^1^Ψ-A pair, suggested that the effect of the presence of Ψ/m^1^Ψ on the stability of RNA compared to U was observed to be consistent across the nearest neighbor (NN) base pairs ^40^.

The U-G, U-U, and U-C non-standard base pairs are commonly found in mRNA, rRNA, viroids, and retroviruses ^39^. U-G wobble pairs occur mostly at the end of helical stems and are non-isosteric with the G-U wobble pairs which mostly occur at the beginning of the helical stems ^40,41^. The structural properties of the U-G, U-U, and U-C pairs in the native as well as in the modified forms has been investigated by a number of studies till date ^39,40,42–48^. Libri et al (2002), investigated the role of Ψ-U mismatch in the recognition of 5’ splice site introns by U1 snRNP and suggested the stabilizing effect of this modification on the interaction of the U1 snRNA with its RNA substrate ^49^. However, they also reported the destabilization of short dsRNA oligonucleotide containing Ψ-U mismatch ^49^. Later, Kierzek et al (2014) in their thermodynamic study, reported the stabilization of dsRNA oligomer containing U-U mismatch upon the inclusion of the Ψ modification ^4^. Earlier studies suggested that the U-G and U-C base pairs, stabilized by interbase water-mediated hydrogen bonding interactions are incorporated within dsRNA structures with very small distortions of the A-RNA sugar-phosphate backbone ^39,42^. In the present work, we have studied the structural, dynamical, hydration and thermodynamic properties of 9-mer RNA duplexes of four different sequence contexts containing an internal m^1^Ψ-A pair, i.e. duplexes with the 5′-Gm^1^ΨC-3′/3′-CAG-5′, 5′-Cm^1^ΨG-3′/3′-GAC-5′, 5′-Um^1^ΨA-3′/3′-AAU-5′ and 5′-Am^1^ΨU-3′/3′-UAA-5′ motifs using molecular dynamics (MD) simulations (Table 1). Additionally, we have also analyzed the properties of 9-mer RNA duplexes (with the 5′-Gm^1^ΨC-3′/3′-CXG-5′ motif, X is G/U/C) containing internal m^1^Ψ-G, m^1^Ψ-U and m^1^Ψ-C pairs (or mismatches) respectively. For comparison we have studied the properties of 9-mer RNA duplexes (with the 5′-GUC-3′/3′-CXG-5′ motif) containing internal U-G, U-U and U-C pairs and RNA duplexes (with the 5′-GΨC-3′/3′-CXG-5′ motif, X is G/U/C) containing internal Ψ-G, Ψ-U and Ψ-C pairs respectively. Additionally, we have explored the hydrogen bonding patterns of the base pairs formed by U, Ψ and m^1^Ψ and the stabilization of the mismatches by interbase water-mediated interactions. Considering the growing importance of m^1^Ψ in mRNA therapeutics, our results might be helpful for better understanding of the role of m^1^Ψ in the structure, stability and function of RNA, especially synthetic therapeutic mRNAs.

**Table 1.**
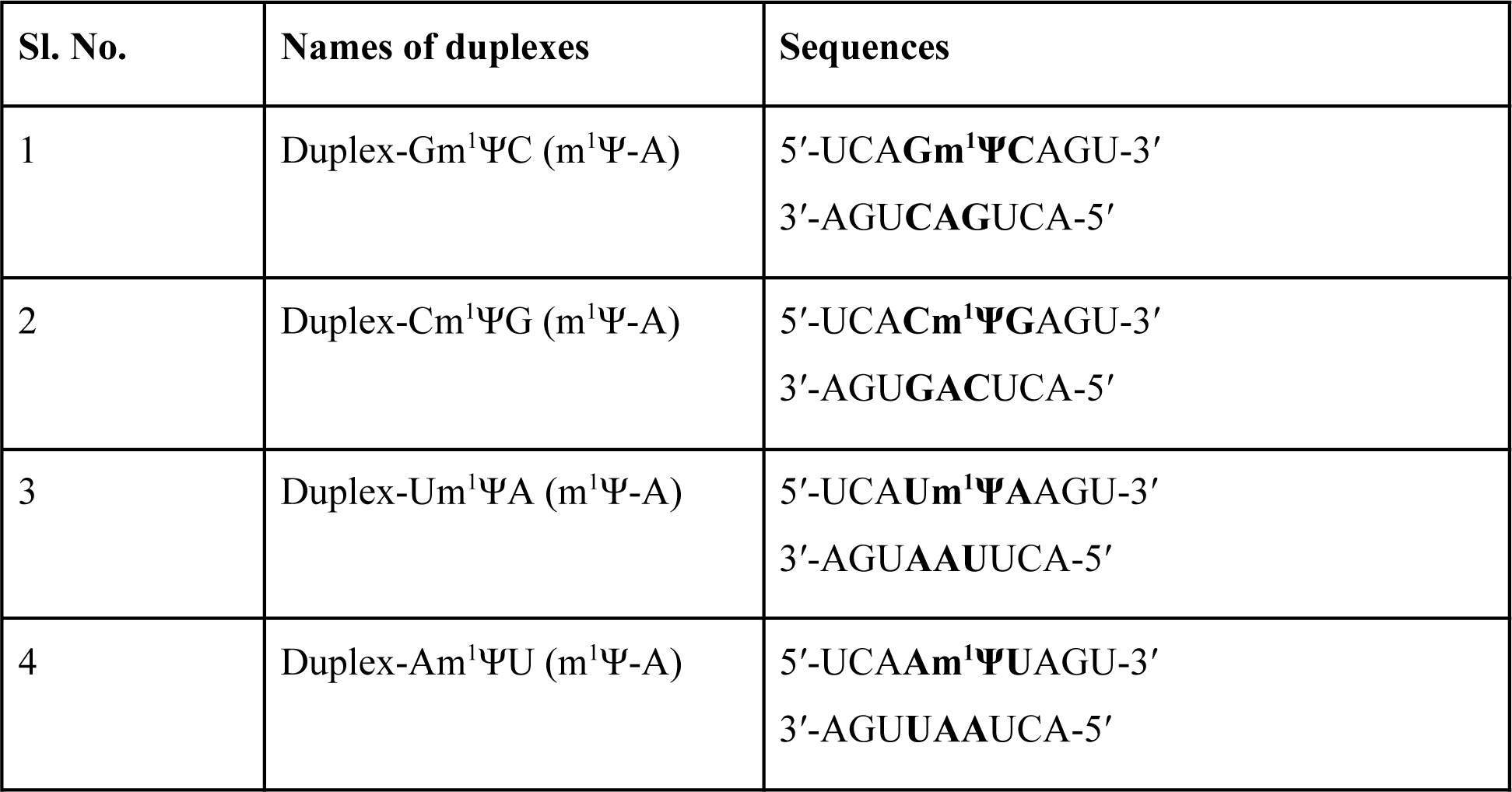
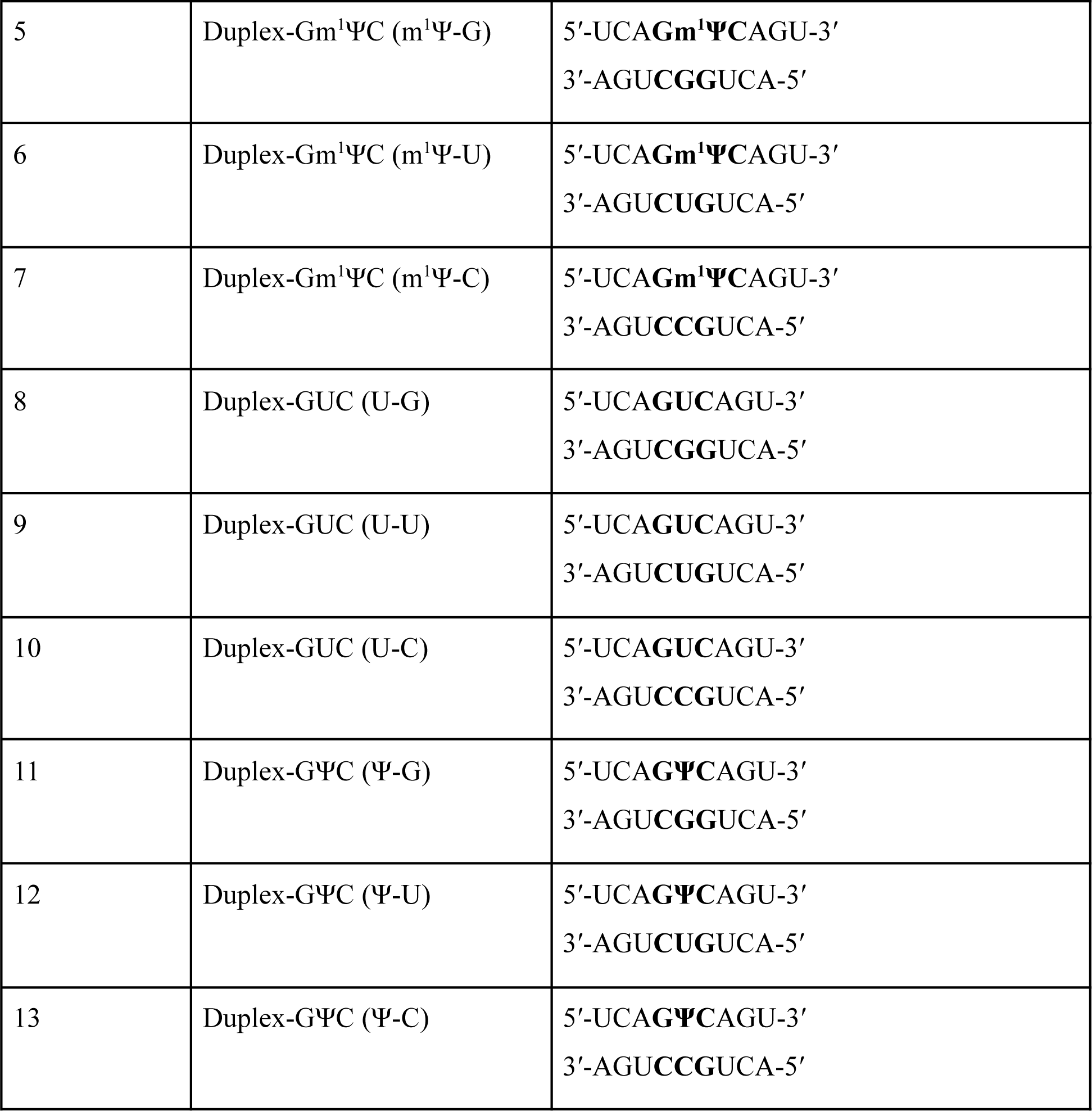
List of the 9-mer RNA duplexes studied in this work.

## METHODS

### Molecular dynamics simulations

The NMR structures of the duplexes corresponding to the GΨC and CΨG contexts (PDB IDs 611W and 611V, respectively)^5^ were obtained from the Protein Data Bank (https://www.rcsb.org/). For each duplex, the first model was modified using the tleap module of AmberTools18 for preparing the duplexes Duplex-Gm^1^ΨC (m^1^Ψ-A) and Duplex-Cm^1^ΨG (m^1^Ψ-A). The other 9-bp duplex structures (as described in Table 1) were prepared using either Rnacomposer (https://rnacomposer.cs.put.poznan.pl) or the nab utility of AmberTools18. The m^1^Ψ modification was incorporated at the 5th position by the rearrangement of atomic positions of U and addition of the methyl group at the N1-atom by using the tleap module. The RNA_OL3 force field parameters ^50^ were used for the regular RNA residues, ff99_bsc0_*χ*_IDRP_ force field parameters were used for the Ψ residue (to be consistent with the simulations Ψ-A pair by Deb et al. ^5^ and for the m^1^Ψ residue, the ff99_bsc0_*χ*_ND_ force field parameters recently reported by us^37^ were used.

For the duplexes containing m^1^Ψ (duplexes 1-7), the initial structures were solvated with TIP3P water molecules in truncated octahedral boxes with the minimum distance of 11 Å between any atom of the solute and the edge of the periodic box and the systems were then neutralized using Na+ ions. The energy minimization of each of the solvated and neutralized systems was carried out in three sets. Each of these sets consisted of 200 steps of steepest descent followed by 300 steps of conjugate gradient with a weak RMS force convergence criterion of 0.1 kcal/mol for the energy gradient. The solute molecules (duplexes) were held by a 25.0 kcal/mol-Å^2^ positional restraining force and by a 5.0 kcal/mol-Å^2^ positional restraining force in the first and the second set of minimization respectively. The duplexes were minimized in the absence of any restraints in the last set of minimization.

The energy minimized systems were then heated from 0 K to 300 K in 100 ps using constant volume Langevin dynamics with a collision frequency of 1 ps^-1^ holding the duplexes with a 15 kcal/mol-Å^2^ position restraint force. Next, the systems were equilibrated in four consecutive steps. In the first equilibration step, constant pressure Langevin dynamics using a collision frequency of 1 ps^-1^ was carried out at 300 K for 100 ps with a 2 fs time step to equilibrate the solvent molecules while the solute molecules (duplexes) were held by a positional restraint force of 10.0 kcal/mol-Å^2^. In the next three steps, 100 ps equilibration runs were performed at 300 K, using constant pressure Langevin dynamics with a collision frequency of 1 ps^-1^ holding the duplexes applying a 5 kcal/mol-Å^2^ restraint force, a 1 kcal/mol-Å^2^ restraint force and a 0.5 kcal/mol-Å^2^ restraint force, respectively. The conditions for the production run were similar to the equilibration steps except the application of the inter-strand Watson-Crick hydrogen bonding distance restraints to the terminal base pairs to avoid fraying of ends according to Saenger ^51^, allowing 0.1 Å movement from the equilibrium bond distance. The production runs were carried out for two independent sets of 500 ns yielding a total of 1 μs simulation for each duplex system and the trajectory files were written after every 10 ps for each set of simulations. Berendsen barostat was used to perform constant pressure simulations by turning on isotropic position scaling with a reference pressure of 1 atm and a pressure relaxation time of 1 ps. The nonbonded interactions within 8.0 Å long-range cutoff were considered during the minimization, equilibration and production steps. The particle mesh Ewald (PME) method was used for handling long-range electrostatic interactions during the simulations. The molecular dynamics simulations were carried out using the AMBER18 software package ^52^ using the GPU version of the pmemd module.

For the duplexes 8-13, The methods for the solvation, neutralization, ionization, minimization, equilibration and production runs for the control duplexes (containing U-G, U-U and U-C pairs) and the Ψ-modified duplexes (containing Ψ-G, Ψ-U and Ψ-C pairs) are provided in the supporting information.

### Analysis of trajectories

AmberTools18 ^52^ was used for the analysis of the simulated ensembles. Root mean square deviations, occurrence of hydrogen bonds and water bridges, distribution of sugar pucker and base orientation were calculated using suitable utilities available within *cpptraj*. All the calculations (except for the time evolution of RMSD (Figure S1) were performed using the last 400ns of the trajectory for each system. Hydrogen bonds were taken into account if the distance between the donor and acceptor atoms <3.0 Å and the donor-hydrogen-acceptor angle was ≥135°. For the representative images of additional water mediated interactions between the base pairs, hydrogen bonds with distance between the donor and acceptor atoms were <3.5 Å and donor-hydrogen-acceptor angle was ≥135° were considered. Cluster analysis was carried out using the average-linkage clustering algorithm with *cpptraj*, with the initial clustering for every 100 frames only. The number of clusters was set to 5 and the minimum distance between clusters greater than 4.0 Å and the coordinate RMSD of the heavy atoms of residues 5 and 14 was used as the distance metric. The water occupancy maps around the average structure obtained from MD were generated using the *grid* routine available in *cpptraj* and UCSF chimera tool (https://www.rbvi.ucsf.edu/chimera/) was utilized for visualization. The molecular mechanics force field-based base stacking and base pairing energies were calculated using the *lie* utility. A cutoff of 999.0 Å for both the electrostatic and van der Waals interactions was considered during these calculations. The energies were obtained as the sum of pairwise electrostatic (ELEC) and van der Waals (VDW) interaction energies between the base atoms. For the calculation of the structural parameters, we used the CURVES^+^/Canal programs ^53^. The free energy (ΔG) of duplex formation was calculated using MM-PBSA.py script availbale within AmberTools18. The MM-GBSA calculations were performed using a total of 4000 frames from the last 400 ns trajectories and an ionic strength of 150 mM using the single trajectory approach.

## RESULTS and DISCUSSION

### Effect of the presence of m^1^Ψ within RNA duplexes

In the present study, we have carried out MD simulations of the RNA duplexes containing central m^1^Ψ-A pairs (listed in Table 1) to understand the effect of N1-methylpseudouridylation on the structure, dynamics and hydration pattern of these duplexes. To examine the differences in the stabilities of the base pairs formed by m^1^Ψ with G, U and C with that of the m^1^Ψ-A pair, we have also simulated RNA duplexes containing central m^1^Ψ-G, m^1^Ψ-U and m^1^Ψ-C pairs (mismatches) respectively. We have compared the results from the simulations of the duplexes containing m^1^Ψ with the observations from the MD simulations of duplexes with the similar sequences containing U-A or Ψ-A pair reported earlier by Deb et al. ^5^ and the observations from the simulation of the U-G, U-U and U-C pairs and also Ψ-G, Ψ-U and Ψ-C pairs in the present work.

#### Structural properties of the duplexes

The calculations of the root mean square deviations (RMSD), radius of gyration (Rg) and inter-strand C1’-C1’ distances for the duplexes under this study, revealed that the duplexes containing the m^1^Ψ-A pair as well as those with the mismatches were able to retain the initial A-RNA conformation (Table S1-2; Figures S2-3).

#### Root mean square deviation (RMSD)

With reference to the minimized structure, Duplex-Gm^1^ΨC (m^1^Ψ-A) and Duplex-Cm^1^ΨG (m^1^Ψ-A) showed lower RMSDs than the other duplexes in this study. The RMSDs of Duplex-Um^1^ΨA (m^1^Ψ-A), Duplex-Am^1^ΨU (m^1^Ψ-A) were similar and ∼0.2 Å greater than Duplex-Gm^1^ΨC (m^1^Ψ-A) and Duplex-Cm^1^ΨG (m^1^Ψ-A) duplexes. On average, the RMSDs of Duplex-Gm^1^ΨC (m^1^Ψ-G), Duplex-GUC (U-G) and Duplex-GΨC (Ψ-G) were observed to be similar. For Duplex-Gm^1^ΨC (m^1^Ψ-U), the RMSD value was found to be greater than what were observed with Ψ and U. Interestingly, for Duplex-Gm^1^ΨC (m^1^Ψ-C), the RMSD was found to be similar to that of Duplex-GUC (U-C) while Duplex-GΨC (Ψ-C) showed ∼0.3 Å lower RMSD than what were observed with m^1^Ψ and U.

#### Radius of gyration

The radius of gyration (Rg) values for the duplexes 1-7 (considering only the heavy atoms) in this study were ∼11 Å. The Rg values for the duplexes containing m^1^Ψ-A pair were slightly greater than the values obtained for the similar duplexes containing U-A and Ψ-A pair ^5^. The Rg values for the duplexes containing m^1^Ψ-G, m^1^Ψ-U and m^1^Ψ-C pairs were also observed to be slightly greater than the values obtained for the similar duplexes containing U-G, U-U, U-C and Ψ-G, Ψ-U, Ψ-C pairs.

#### Intramolecular C1’-C1’ distances and the glycosidic bond angles (λ)

The C1’-C1’ distances for the m^1^Ψ-A pair was observed to be ∼10.8 Å for all contexts and were almost similar to the values for Ψ-A pair and slightly greater than the canonical U-A pair. The corresponding distances for the m^1^Ψ-A, m^1^Ψ-G, m^1^Ψ-U and m^1^Ψ-C pairs were observed to be different. For m^1^Ψ-G and m^1^Ψ-U, the values were respectively ∼0.2 Å and ∼1 Å lesser than that for m^1^Ψ-A, while the C1’-C1’ distance for m^1^Ψ-C was found to be ∼0.3 Å greater than that for m^1^Ψ-A. Similar trend was observed for the U and Ψ mismatches. The C1’-C1’ distances for U-C and Ψ-C pairs were greater than the distances for U-G and Ψ-G pairs followed by those for U-U and Ψ-U pairs, respectively. In general, The C1’-C1’ distances for U-U, Ψ-U and m^1^Ψ-U were found to be lesser than the other pairs. The C1’-C1’ distances of the two different geometries observed for the U-U, Ψ-U and m^1^Ψ-U pairs were similar and the values for the Ψ-U and m^1^Ψ-U pairs were ∼0.2 Å lesser than the U-U pair for the respective geometries. The average C1’-C1’ distances for the two geometries differed by ∼1.6, ∼0.6 and ∼1.2 Å respectively for U-C, Ψ-C and m^1^Ψ-C pairs.

On average, for all the duplexes containing m^1^Ψ-A pair in this study, the λ angles for m^1^Ψ(5) (C1’-C1’-C5) and A(14) (C1’-C1’-N9) nucleotides were found to be ∼52° and ∼57° respectively (Table S2 (C)). In the m^1^Ψ-G pair, the λ angles for m^1^Ψ was observed to be ∼69° and ∼46° respectively and for the Ψ-G and U-G pairs similar values were observed. For the m^1^Ψ-U and m^1^Ψ-C pairs as well as for the base pairs of U and Ψ with U and C respectively, we observed larger deviations of the values of the λ angles from the mean values during the course of the simulation due to the occurrence of two different geometries. More detailed discussion on the changes in the C1’-C1’ and λ angles with the base pair type and base pair geometries is provided later in this manuscript. The time evolution of the λ angles for the mismatches formed by U, Ψ and m^1^Ψ are shown in Figure S4.

#### Sugar pucker conformation and base orientation

The 5th (m^1^Ψ) and 14th (A/G/U/C) residues were observed to be in NORTH/ANTI conformation of sugar pucker and base orientation (Table S3 (A,B)). The glycosidic torsion angle (*χ)* for m^1^Ψ(5) was found to be ∼200° for duplex-Gm^1^ΨC (m^1^Ψ-A) and duplex-Cm^1^ΨG (m^1^Ψ-A), respectively and ∼204° and ∼202° for duplex-Um^1^ΨA (m^1^Ψ-A), duplex-Am^1^ΨU (m^1^Ψ-A) respectively (Figure S5). The glycosidic torsion angle (*χ)* for Ψ and U have been reported to be ∼195° and ∼205° for similar duplexes^5^. The values were ∼201° for the m^1^Ψ-G and m^1^Ψ-C mismatches and ∼198° for m^1^Ψ-U. For the control and Ψ-modified duplexes also, the 5th and 14th residues were in the NORTH/ANTI conformation of sugar pucker and base orientation (Table S3 (C)). As observed earlier for duplexes containing U-A and Ψ-A pairs^5^, for the duplexes containing mismatches, the glycosidic torsion angle (*χ)* for U5 was found to be ∼195° and that for Ψ5 was ∼205° suggesting that Ψ populated a low ANTI region of the glycosidic torsion space (Figure S6). The other residues within the RNA duplexes under this study were also observed to populate the NORTH/ANTI region of sugar pucker conformation and base orientation (data not shown).

#### Local base pair and base pair step parameters

To investigate the effect of the modifications on the local stability of the duplexes we analyzed the local base pair and base pair step parameters for each of the duplexes (Tables S4-9). The base pair parameters and base pair step parameters for the duplexes of the four different sequence contexts containing m^1^Ψ-A pair, were observed to be similar (Table S4, S6, Figures S7-8). However, there were significant differences in some of these parameters for the m^1^Ψ-A, m^1^Ψ-G, m^1^Ψ-U and m^1^Ψ-C pairs (Tables S4-7, Figures S9-10). The base-pair opening for the mismatches were found to be greater than the m^1^Ψ-A pair. For the m^1^Ψ-G pair, the value was observed to be somewhat greater than those for the U-G and Ψ-G pairs. The U-C pair showed the largest base pair opening out of all the base pairs studied in this work (Figures S11). The values of shear for the mismatches, particularly m^1^Ψ-G and m^1^Ψ-C, were significantly greater than what was observed for m^1^Ψ-A. For the control and Ψ-modified duplexes also, the values of shear were found to be greater for the mismatches compared to that for the U-A or Ψ-A pairs, respectively. The values of twist of the base pair steps containing m^1^Ψ showed much greater deviations from the mean values for the m^1^Ψ-U and m^1^Ψ-C pairs. Similarly, greater deviations from the mean value of twist for the U-U, U-C, Ψ-U and Ψ-C pairs were observed compared to the other pairs for the control and Ψ-modified duplexes. Larger values of shift were observed for the base pair steps containing m^1^Ψ-C pair compared to the other pairs with m^1^Ψ. Greater shifts were observed for base pair steps containing U-C, Ψ-C and m^1^Ψ-C pairs than the other mismatches with U, Ψ and m^1^Ψ, respectively (Figure S12).

#### Intramolecular hydrogen bond analysis

The analysis of the intramolecular hydrogen bond interactions revealed the formation of (m^1^Ψ)N3-HN3---(A)N1 and (A)N6-H61---(m^1^Ψ)O2 hydrogen bonds between m^1^Ψ(5) and A(14) for all the duplexes containing the m^1^Ψ-A pair (Table 2, S10, Figure 2). The frequency of the formation of these hydrogen bonds were similar for all four contexts.

**Figure 2.**
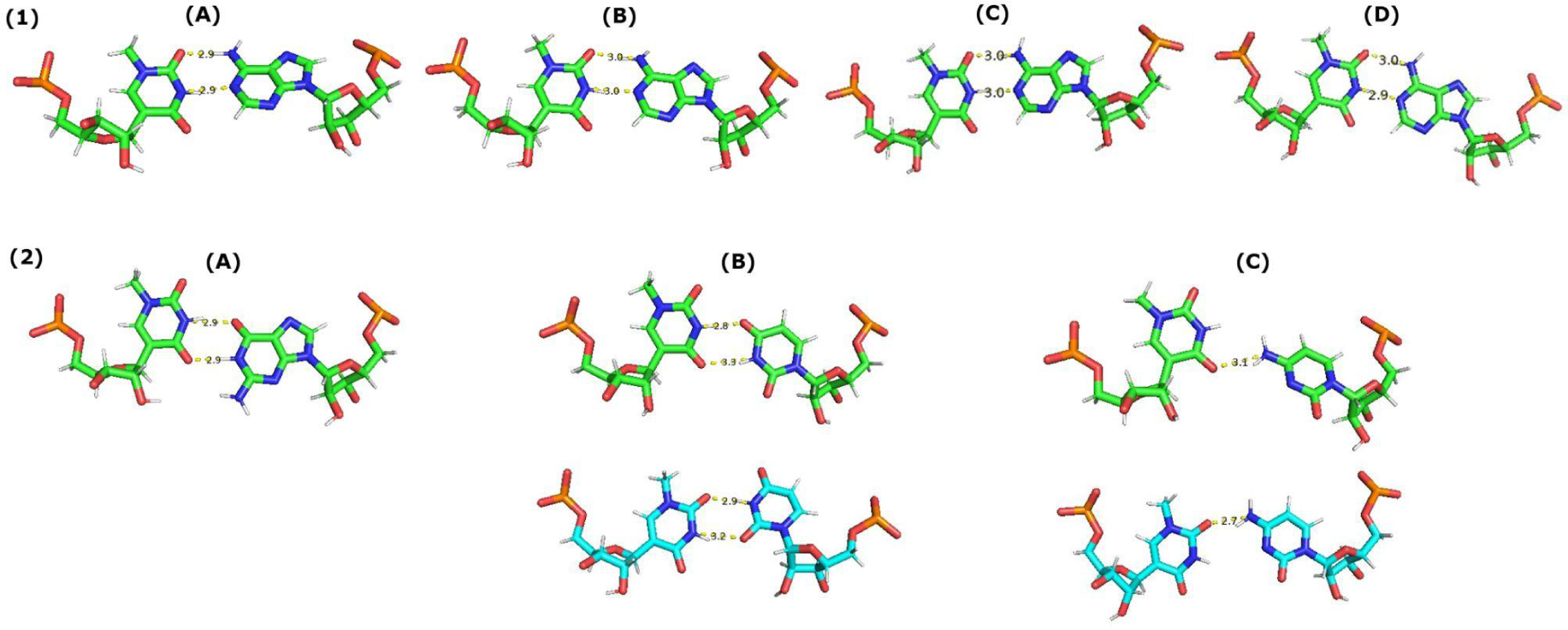
(1) Observed geometries of the m^1^Ψ-A pair within the duplexes (A) Duplex-Gm^1^ΨC, (B) Duplex-Cm^1^ΨG, (C) Duplex-Um^1^ΨA and (D) Duplex-Am^1^ΨU corresponding to the centroid structure obtained from the most populated cluster. (2) Observed geometries of the (A) m^1^Ψ-G, (B) m^1^Ψ-U and (C) m^1^Ψ-C mismatches respectively within Duplex-Gm^1^ΨC. For m^1^Ψ-G mismatch, the geometry corresponds to the centroid structure obtained from the most populated cluster. For the m^1^Ψ-U and m^1^Ψ-C mismatches, the geometries corresponding to the centroid structures obtained from the most populated cluster (top) and the second most populated cluster (bottom) are shown. Carbons are indicated in green for the most populated cluster and in cyan for the second most populated cluster, oxygens are shown in red, phosphorus in orange and nitrogens in blue respectively.

**Table 2.** Occurrence and the possible hydrogen bonding interactions of the different geometries for the mismatches

| Base pair | Geometry 1 |  | Geometry 2 |  |
| --- | --- | --- | --- | --- |
|  | Occurrence (%) | Possible H-bond/s | Occurrence (%) | Possible H-bond/s |
| U-G | 92 | (G14)N1-H1---(U5)O2 | NA |  |
|  |  | (U5)N3-HN3--- (G14)O6 |  |  |
| Ψ-G | 93 | (G14)N1-H1---(Ψ5)O4 | NA |  |
|  |  | (Ψ5)N3-HN3---(G14)O6 |  |  |
| m<sup>1</sup>Ψ-G | r1<sup>a</sup> 73 | (G14)N1-H1---(m<sup>1</sup>Ψ5)O4 | NA |  |
|  | r2 84 | (m<sup>1</sup>Ψ5)N3-HN3--- (G14)O6 |  |  |
| U-U | 49 | (U14)N3-H3---(U5)O4 | 41 | (U5)N3-H3- - - (U14)O4 |
|  |  | (U5)N3-H3---(U14)O2 |  | (U14)N3-H3- - - (U5)O2 |
| Ψ-U | 25 | (U14)N3-H3- - -(Ψ5)O2 | 68 | (Ψ5)N3-HN3- - - (U14)O4 |
|  |  | (Ψ5)N3-H3- - -(U14)O2 |  | (U14)N3-HN3- - - (Ψ5)O4 |
| m<sup>1</sup>Ψ-U | r1 40 | (U14)N3-H3- - -(m<sup>1</sup>Ψ5)O2 | r1 55 | (m<sup>1</sup>Ψ5)N3-HN3- - - (U14)O4 |
|  | r2 44 | (m<sup>1</sup>Ψ5)N3-H3- - -(U14)O2 | r2 48 | (U14)N3-H3---(m<sup>1</sup>Ψ5)O4 |
| U-C | 50 | (C14)N4-H41- - -(U5)O4 | 43 | (C14)N4-H41- - -(U5)O2 |
| Ψ-C | 35 | (C14)N4-H41- - -(Ψ5)O2 | 55 | (C14)N4-H41- - -(Ψ5)O4 |
| m<sup>1</sup>Ψ-C | r1 42 | (C14)N4-H41- - -(m<sup>1</sup>Ψ5)O2 | r1 45 | (C14)N4-H41- - -(m<sup>1</sup>Ψ5)O4 |
|  | r2 37 |  | r2 43 |  |

<sup>a</sup> Here r1 and r2 indicate the two independent sets (replicates) of simulation

We observed (G)N1-H1---(m^1^Ψ)O4 and (m^1^Ψ)N3-HN3---(G)O6 H-bonds for the m^1^Ψ-G pair, (G)N1-H1---(Ψ)O4 and (Ψ)N3-HN3---(G)O6 H-bonds for the Ψ-G pair and (G)N1-H1---(U)O2 and (U)N3-HN3---(G)O6 H-bonds for the U-G pair. The N1-H1---O4/O2 H-bond was observed to be more frequent compared to the N3-HN3---O6 H-bond (Table S11).

Hydrogen bonding patterns differed between the two geometries observed for each of the Y-U and Y-C pairs (where Y is U/Ψ/m^1^Ψ). The details of the H-bonds observed are provided in Table 2. In general the frequencies of the H-bonds observed for the Y-U and Y-C pairs were much lesser compared to those observed for the Y-A and Y-G pairs (Table S11).

#### Hydration properties of the duplexes

##### Formation of water bridges between phosphate backbone atoms

Due to the difference in the salt concentration in the simulated system for the duplexes containing m^1^Ψ(5) (duplexes 1-7) with that for the duplexes containing U/Ψ(5) (duplexes 8-13), their hydration patterns are not directly comparable. But we have discussed here about the observed changes in the hydration pattern for duplexes containing different base pairs for m^1^Ψ/Ψ/U.

The occupancies of water bridges formed between the phosphate oxygen atoms of m^1^Ψ(5) and those of its neighboring residues (4^th^ and 6^th^) for the duplexes containing m^1^Ψ-A pair were observed to vary depending on the sequence context (Table S12). The water occupancy maps for the duplexes (residues 4-6) in this study are shown in Figures S13-14. The occupancy for the OP2(5)-W-OP1(4) water bridge was found to be the highest for the Gm^1^ΨC context and that of the OP2(5)-W-OP2(4) water bridge interaction was found to be the highest for the Um^1^ΨA context. The occupancies of the OP2(6)-W-OP1(5) and OP2(6)-W-OP2(5) were observed to be significantly lesser for the Cm^1^ΨG and Um^1^ΨA contexts than what were observed for the Gm^1^ΨC and Am^1^ΨU contexts. Interestingly, for duplex-Gm^1^ΨC (m^1^Ψ-G), the occupancy of the OP2(5)-W-OP1(4) water bridge was found to be much greater than what were observed for the other duplexes (Table S13). The OP2(5)-W-OP1(4) water bridge interaction was more frequent for the m^1^Ψ-C pair than the duplexes (of context Gm^1^ΨC) containing m^1^Ψ-A or m^1^Ψ-U pairs. The OP2(5)-W-OP2(4) water bridge was found to be less frequent for the m^1^Ψ-U and m^1^Ψ-C pairs compared to the other duplexes. The occupancies of the OP2(6)-W-OP1(5) and OP2(6)-W-OP2(5) water bridge interactions were observed to be <5% for m^1^Ψ-G, while those for the m^1^Ψ-U pair followed by m^1^Ψ-C pair, were greater than what were observed for the other duplexes.

The OP2(5)-W-OP1(4) and OP2(5)-W-OP2(4) water bridges were found to be more frequent for duplex-GUC (U-G) and duplex-GΨC (Ψ-G) compared to what were observed for the similar duplexes containing U-A and Ψ-A pairs respectively ^5^ (Table S13) and the frequencies were slightly greater for duplex-GΨC (Ψ-G) than duplex-GUC (U-G). The frequencies of these two water bridges were observed to be similar for duplex-GUC (U-U) and duplex-GUC (U-C) and a little lesser than that for the duplex containing the canonical U-A pair. On the other hand, for duplex-GΨC (Ψ-U) and duplex-GΨC (Ψ-C), the frequencies of these water bridges were greater than those for duplex-GΨC (Ψ-A)^5^ as well as those for their canonical counterparts. The OP2(6)-W-OP1(5) and OP2(6)-W-OP2(5) water bridges were significantly less frequent for the duplex-GUC (U-G) and duplex-GΨC (Ψ-G) than for the other control and Ψ-modified duplexes. The frequencies of the OP2(6)-W-OP1(5) water bridge for the duplex-GUC (U-U) and duplex-GΨC (Ψ-U) pairs were observed to be higher than those for the other control and Ψ-modified duplexes, respectively. The frequencies of the OP2(6)-W-OP2(5) water bridge for duplex-GΨC (Ψ-U) and duplex-GΨC (Ψ-C), were found to be more than those for duplex-GΨC (Ψ-A) and duplex-GΨC (Ψ-G) and the observation was similar for the control duplexes.

For the Ψ-modified duplexes, the water bridging interaction between the OP2 and HN1 atoms of Ψ(5) were also observed while occurrence of such water bridge interaction can not be observed for the control duplexes due to the absence of the additional hydrogen bond donor, i.e. the N1H imino group. The occupancy of this water bridge was observed to be lower for duplex-GΨC (Ψ-U) than the other Ψ-modified duplexes. For the simulated Ψ-modified duplexes (duplexes 11-13) containing Ψ-G, Ψ-U and Ψ-C mismatches we observed the formation of water bridges between the OP2 atoms of Ψ(5) and its preceding ((i.e. its 5’ neighboring residue) G(4) residue mediated by the HN1 atom of Ψ(5) (Figure S15). In agreement with earlier experimental reports in/for different contexts of RNA ^54–56^, within the duplexes simulated in this study the presence of one structural water molecule coordinated between the OP2 atoms of Ψ(5) and G(4) and the HN1 atom of Ψ(5) was observed. Interestingly, a string of two bridging water molecules between these atoms was also observed during the simulation of these duplexes. The occurrence of two bridging water molecules were also observed for the duplexes of each of the four different contexts containing Ψ-A pair in our earlier study^5^. The presence of a chain of two water residues coordinated between the OP2 and HN1 atoms of Ψ and the OP2 atom of the 5’ G nucleotide has been also reported in the crystal structure of a U2 snRNA-intron branch site duplex (PDB ID: 3CGP) ^56^. As mentioned earlier, such water-mediated interactions involving Ψ, have been suggested to improve the stability of base stacking interactions by enhancing the phosphodiester backbone rigidity towards the 5′ direction^36^.

##### Radial distribution function

For all the m^1^Ψ-modified duplexes in the present study, the analysis of RDF of water oxygen atoms around the methyl hydrogen atom (H20) of m^1^Ψ(5) showed the presence of water molecules at a distance of ∼2.9 Å (Figures S16-17) Å and that around the methyl carbon (C10) of m^1^Ψ(5) was observed to be 3.3 Å. Calculation of the RDF of water oxygen atoms around the H5 atom of U(5) and that about the HN1 atom of Ψ(5) for the control and Ψ-modified duplexes showed a well-defined first hydration shell which formed between 1.5 Å and 2.5 Å with a peak at 2.05 Å. The water molecules were observed to be farther from the H5 atom of U5 for the control duplexes with the first solvation peak located at ∼2.9 Å (Figure S18). The first solvation peak with respect to the H5 atom of U(5) was found to be of significantly lower intensity compared to that of the HN1 atom of Ψ(5), especially for the duplex-GΨC (Ψ-C). Similarly, the occupancy of water molecules was observed to be much higher around the geometric center between the OP2 and HN1 atoms of Ψ(5) compared to that around the geometric center between the OP2 and H5 atoms of U(5) (Figure S19) for the duplexes under this study. These results are similar to what was observed for the duplexes containing U-A and Ψ-A pairs for the four sequence contexts in our earlier report ^5^.

#### Overall comparison of the structural properties of the base pairs formed by U, Ψ and m^1^Ψ with the canonical nucleotides

In the present study, we observed the formation of two hydrogen bonds similar to those in U-A and Ψ-A pairs, i.e. (m^1^Ψ)N3-HN3---(A)N1 and (A)N6-H61---(m^1^Ψ)O2 for the m^1^Ψ-A pair with both the residues in NORTH/ANTI conformation and an average C1’-C1’ distance of ∼10.8 Å, slightly greater than that for the canonical U-A pair, i.e. ∼10.6 Å as was also observed for Ψ-A pair within similar duplexes ^5^. The λ angles for the m^1^Ψ(5) and A(14) residues were observed to be nearly identical (∼55°) as has been reported for the WC U-A pairs ^39,41,59–62^. For the duplex under this study, U and G formed wobble base pair with two H-bonds (G)N1-H1---(U)O2 and (U)N3-HN3---(G)O6, with both the residues in NORTH/ANTI conformation and a C1’-C1’ distance ∼10.6 Å similar to the distance observed in experimental structures ^43,62–66^. Ψ and m^1^Ψ also formed wobble base pair with G two hydrogen bonds (G)N1-H1---(Ψ/m^1^Ψ)O4 and (Ψ/m^1^Ψ)N3-HN3---(G)O6 with C1’-C1’ distance ∼10.6 Å. The λ angles for U-G and Ψ-G pairs were observed to be similar i.e. ∼68° (U/Ψ) and ∼44° for G respectively, similar to the previously reported values for the U-G pair ^40,58^. The average shift of −12.5° and +15° of the λ angles from the values observed for WC base pairs (i.e the opposite drifts of the λ angles), respectively for the purine and pyrimidine bases forming a wobble base pair has been reported in earlier studies ^58,63–65^ Other than the two direct hydrogen bonding interactions, the U-G wobble base pair was also observed to be additionally supported either by two key water molecules (one in each groove) i.e. by the O4(5)-W-O6(14) and O2(5)-W-N2(14) (or O2’-O2(5)-W-N2(14) i.e the same water molecule bridging O2’ and O2 atoms of U and N2 atom of G) or by three key water molecules i.e O4(5)-W-O6(14) and O2’-O2(5)-W-N2(14) interactions and an water molecule linking the O4(5)-W-O6(14) with N7(14)) (Table S14; Figure S20) and similar water mediated interactions have been reported in earlier studies ^39,41,42,58,62^. Such stabilizing water mediated interactions between the bases were also observed for the Ψ-G and m^1^Ψ-G pairs i.e. by two key water molecules (O2(5)-W-O6(14) and O4(5)-W-N2(14) water bridges (or O2’-O4(5)-W-N2(14) i.e the same water molecule bridging O2’ and O2 atoms of U and N2 atom of G) or by three key water molecules i.e O2(5)-W-O6(14) and O2’-O4(5)-W-N2(14) interactions and a water molecule linking the O2(5)-W-O6(14) with N7(14)) (Table S14; Figures 3,S21). The frequencies of the water bridging interactions providing additional support to the mismatches in this study are tabulated in Table 1.

**Figure 3.**
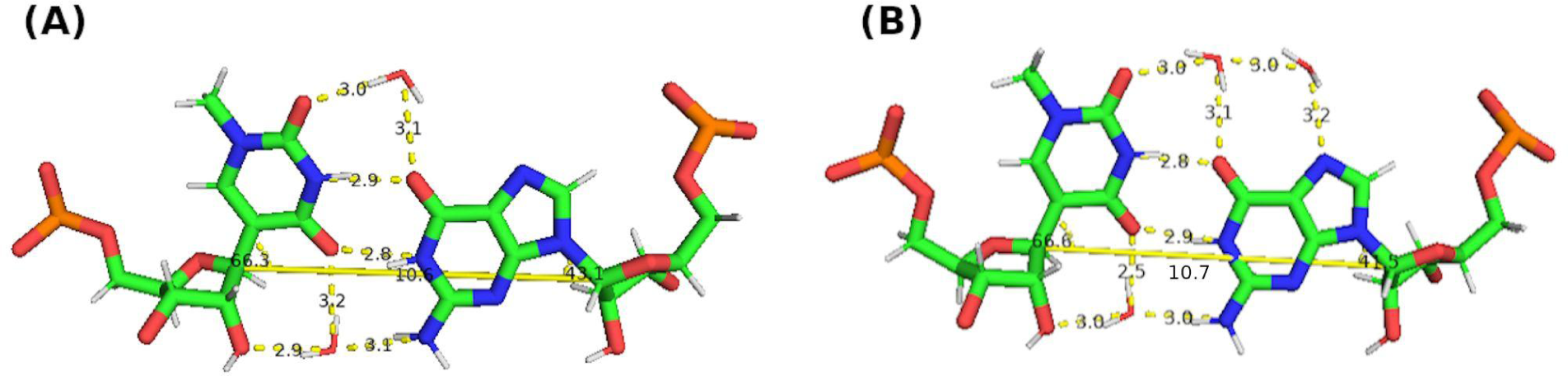
Snapshots of m^1^Ψ-G wobble base pair (corresponding to Duplex-Gm^1^ΨC (m^1^Ψ-G)) with water mediated interactions between the bases (A) by two key water molecules and (B) by three key water molecules, other than the two direct hydrogen bonding interactions. Distances between hydrogen bond acceptor and donor atoms are shown with yellow dotted lines. The C1’-C1’ distances and λ ( C1’-C1’-C5 for m^1^Ψ and C1’-C1’-N9 for G) angles are also shown.

For the duplex in the present study, U(5) formed wobble base pair with U(14) with either (U14)N3-H3---(U5)O4 and (U5)N3-H3---(U14)O2 hydrogen bonds (geometry 1) or (U5)N3-H3---(U14)O4 and (U14)N3-H3---(U5)O2 (geometry 2) hydrogen bonds with both uridine residues in the NORTH/ANTI conformation and an average C1’-C1’ distance of ∼9.9 Å. Sheng et al, (2014) in their study, observed such geometries for the U-U pair in 7-mer dsRNA crystal structures ^48^. Wobble base pairs of Ψ/m^1^Ψ(5) with U(14) were also observed with two direct hydrogen bonds i.e. either (14)N3-H3---(5)O2 and (5)N3-H3---(14)O2 hydrogen bonds (geometry 1) or (5)N3-H3---(14)O4 and (14)N3-H3---(5)O4 hydrogen bonds (geometry 2) with both uridine residues in the NORTH/ANTI conformation and C1’-C1’ distance of ∼9.7 Å, corresponding to the two most frequently observed geometries. The opposite drifts of the λ angles for the two pyrimidine (U/Ψ/m^1^Ψ(5) and U(14)) residues were observed. For each of the U-U, Ψ-U and m^1^Ψ-U pairs, corresponding to each of the two most frequently observed geometries, two different conformations i.e. a symmetric hydrogen-bonded conformation (C1’-C1’ distance of ∼10.5 Å) and a ‘stretched’ wobble conformation (C1’-C1’ distance of ∼9 Å), stabilized by additional water mediated interactions between the bases were observed as reported for the U-U base pair in earlier studies ^46,62^ (Figure 4; S22-23). Two key water molecules were observed (one in each groove) mediating the O4(5)-W-O4(14) and O2(14)-W-O2(5) water bridging interactions for the symmetric H-bonded U-U pair and O4(5)-W-O2(14) and O4(14)-W-O2(5)) interactions for the Ψ-U and m^1^Ψ-U pairs. Unlike the symmetric hydrogen-bonded wobble base pair, the asymmetric ‘stretched’ wobble pair contained only one direct hydrogen bond (i.e. (14)N3-H3---(5)O4 or (5)N3-H3---(14)O4 for U-U and (14)N3-H3---(5)O2 or (5)N3-H3---(14)O4 for Ψ-U and m^1^Ψ-U), a greater C1’-C1’ distance, lower values of λ angles and one residue (5th or 14th depending on the geometry) inclined towards the minor groove. Additional stabilization of the ‘stretched’ wobble base pairs was observed to be provided by two key water molecules mediating the O4(5)-W-O4(14) and O2(14)-W-N3(5) (geometry 1) or O4(5)-W-O4(14) and O2(5)-W-N3(14) (geometry 2) water bridging interactions for the U-U pair and (O4(14)-W-O2(5) and O2(14)-W-N3(5) (geometry 1) or O4(14)-W-O2(5) and O4(5)-W-N3(14) (geometry 2) interactions for the Ψ-U and m^1^Ψ-U pairs (Table S14; Figures 4,S22-23). A single hydrogen bonded ((U)N3-H3---(Ψ)O2) structure along with a O2(^2m^U)-W-N3(Ψ) water bridge has been observed for the Ψ-^2m^U mismatch within the tRNA^Gln^ anticodon loop ^66^.

**Figure 4.**
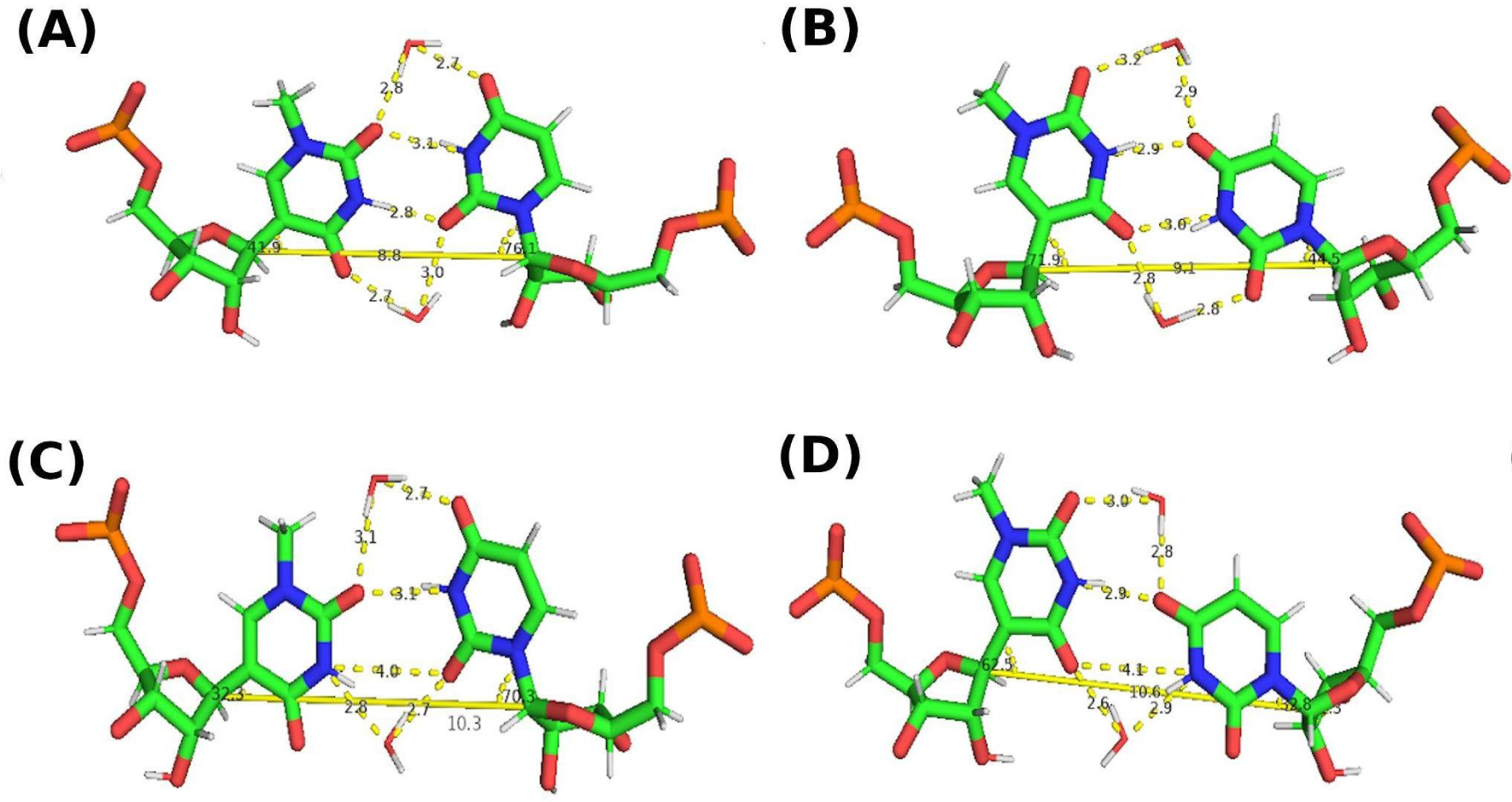
Snapshots of (A,B) Symmetric hydrogen bonded m^1^Ψ-U wobble base pair with water mediated interactions (O4(5)-W-O2(14) and O4(14)-W-O2(5)) by two key water molecules, other than the two direct hydrogen bonding interactions, respectively for the two geometries observed in this study and (C,D) ‘stretched m^1^Ψ-U’ wobble base pair with water mediated interactions (O2(14)-W-O4(5) and O2(14)-W-N3(5) (geometry 1) or O4(14)-W-O2(5) and O4(5)-W-N3(14) (geometry 2)) by two key water molecules other than one direct hydrogen bond, respectively for the two geometries observed in this study (corresponding to Duplex-Gm^1^ΨC (m^1^Ψ-U)). Distances between hydrogen bond acceptor and donor atoms are shown with yellow dotted lines. The C1’-C1’ distances and λ (C1’-C1’-C5 for m^1^Ψ and C1’-C1’-N1 for U) angles are also shown.

For the U-C pair, single hydrogen bonded geometries (with either (C14)N4-H41---(U5)O4 (geometry 1) or (C14)N4-H41---(U5)O2 (geometry 2) hydrogen bond) with both the residues in NORTH/ANTI and an average C1’-C1’ distance of ∼11.7 Å was observed. Similarly the Ψ-C and m^1^Ψ-C pairs also formed single hydrogen bonded geometries (with either (14)N4-H41---(5)O2 (geometry 1) or (14)N4-H41---(5)O4 (geometry 2) hydrogen bonds) with both the residues in the NORTH/ANTI conformation and average C1’-C1’ distances of ∼10.9 Å and ∼11 Å respectively. For the conformers with (14)N4-H41---(5)O4 H-bond for U-C and those with (14)N4-H41---(5)O2 H-bond for Ψ-C and m^1^Ψ-C pairs, the C1’-C1’ distances were found to be greater than that for the conformers with (14)N4-H41---(5)O2 H-bond for U-C and those with (14)N4-H41---(5)O4 H-bond for Ψ-C and m^1^Ψ-C pairs (Figures 5,S24-25). We also observed a geometry with two hydrogen bonds for the U-C ((14)N4-H41-O4(5) and (5)N3-HN3---N3(14) hydrogen bonds) as reported by ^47^ and also for the Ψ-C and m^1^Ψ-C pairs ((14)N4-H41-O4(5) and (5)N3-HN3---N3(14) H-bonds) but the population of the conformers containing this hydrogen bonding pattern was found to be very low. Additional support to the U-C pair was observed to be provided by either one key water molecule (N3(5)-W-N3(14)) or by five key water molecules (i.e. N3(5)-W-N3(14) and four water molecules connecting O4(5), N4(14) and phosphate oxygen of C14) other than the one direct hydrogen bonding interaction in geometry 1 and by one key water molecule (O2(5)-W-N3(14)) or by two key water molecules (O2(5)-W-N3(14) and (O4(5)-W-N4(14)) in geometry 2 (Figure S24). The additional stabilization of the U-C pair (containing a single hydrogen bond) in RNA duplex by water mediated interactions has been reported by Holbrook et al, (1991) and Cruse et al, (1994) ^39,42^. For the Ψ-C and m^1^Ψ-C pairs, the additional stabilization was observed to be provided by either one key water molecule (N3(5)-W-N3(14)) or by four key water molecules (N3(5)-W-N3(14) and three water molecules connecting O2(5), N4(14) and phosphate oxygen of C14) and else by five key water molecules (N3(5)-W-N3(14) and four water molecules connecting O2(5), N4(14) and phosphate oxygen of C14) for geometry 1 and by either one key water molecule (O4(5)-W-N3(14)) or by two key water molecules (O4(5)-W-N3(14) and (O2(5)-W-N4(14)) for geometry 2 (Figures 5,S25). The water occupancy maps showing the additional water-mediated stabilization of the mismatches are shown in Figures 26-28.

**Figure 5.**
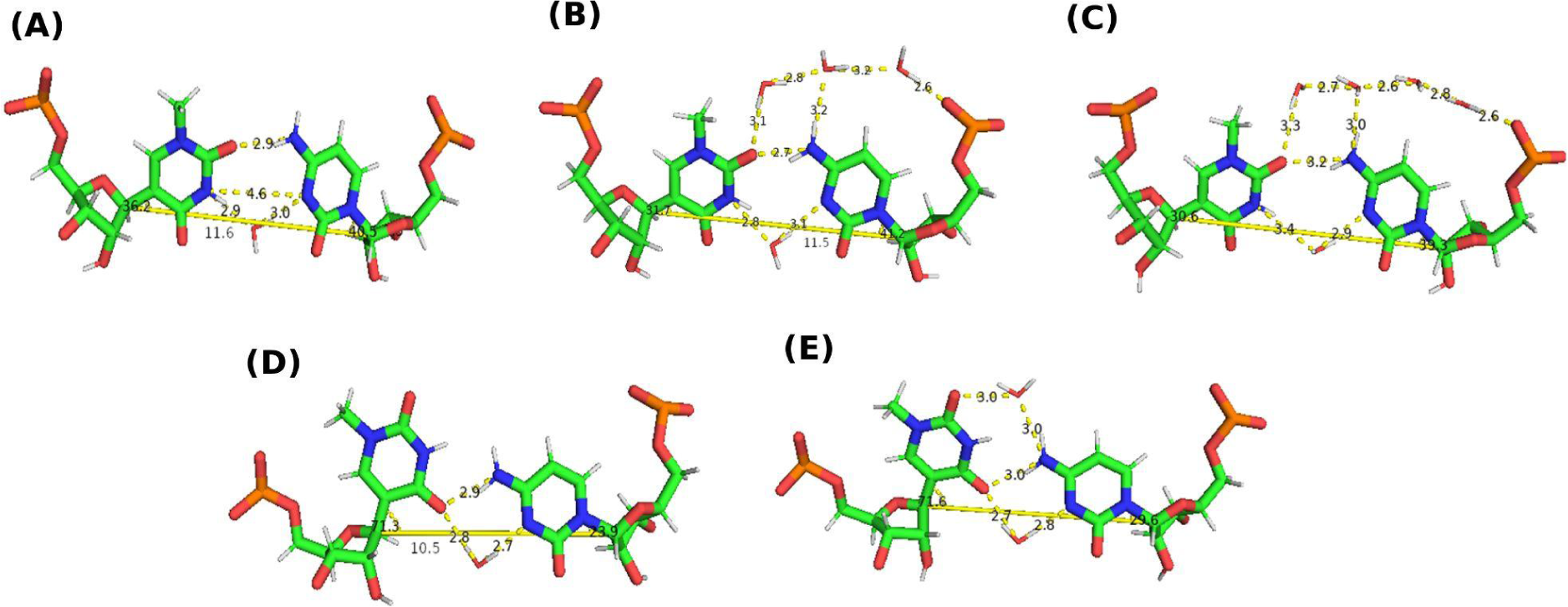
Snapshots of m^1^Ψ-C pair (corresponding to Duplex-Gm^1^ΨC (m^1^Ψ-C)) with water mediated interactions by (A) one key water molecule (N3(5)-W-N3(14)) (B) by 4 key water molecules (N3(5)-W-N3(14) and 3 water molecules connecting O2(5), N4(14) and phosphate oxygen of C14) and (C) by 5 key water molecules (N3(5)-W-N3(14) and 4 water molecules connecting O2(5), N4(14) and phosphate oxygen of C14), other than the one direct hydrogen bonding interaction for geometry 1, (D,E) by one key water molecule (O4(5)-W-N3(14)) or by two key water molecules (O4(5)-W-N3(14) and (O2(5)-W-N4(14)) other than one direct hydrogen bond for geometry 2. Distances between hydrogen bond acceptor and donor atoms are shown with yellow dotted lines. The C1’-C1’ distances and λ ( C1’-C1’-C5 for m^1^Ψ and C1’-C1’-N1 for C) angles are also shown.

##### Base pairing and stacking interactions

In general, for all the sequence contexts, we observed that the base pairing energies for the m^1^Ψ-A pair and the stacking energies for the base pair steps containing m^1^Ψ-A were significantly lesser (i.e. more negative) than what were observed for the U-A and Ψ-A pairs ^5^ indicating that the presence of m^1^Ψ in the duplexes resulted in more stable base pairing and stacking interactions (Table S15, Figure 6–7). The stacking energies were observed to vary depending on the sequence context as was earlier observed for the U-A and Ψ-A pairs ^5^. The following trend was observed for the stacking energies: 5’G-3’C/3’m^1^Ψ-5’A < 5’A-3’U/3’m^1^Ψ-5’A < 5’m^1^Ψ-3’A/3’A-5’U < 5’m^1^Ψ-3’A/3’G-5’C < 5’C-3’G/3’m^1^Ψ-5’A < < 5’U-3’A/3’m^1^Ψ-5’A < 5’m^1^Ψ-3’A/3’C-5’G < 5’m^1^Ψ-3’A/3’U-5’A corresponding to the duplexes containing the m^1^Ψ-A pair (Figure 7).

**Figure 6.**
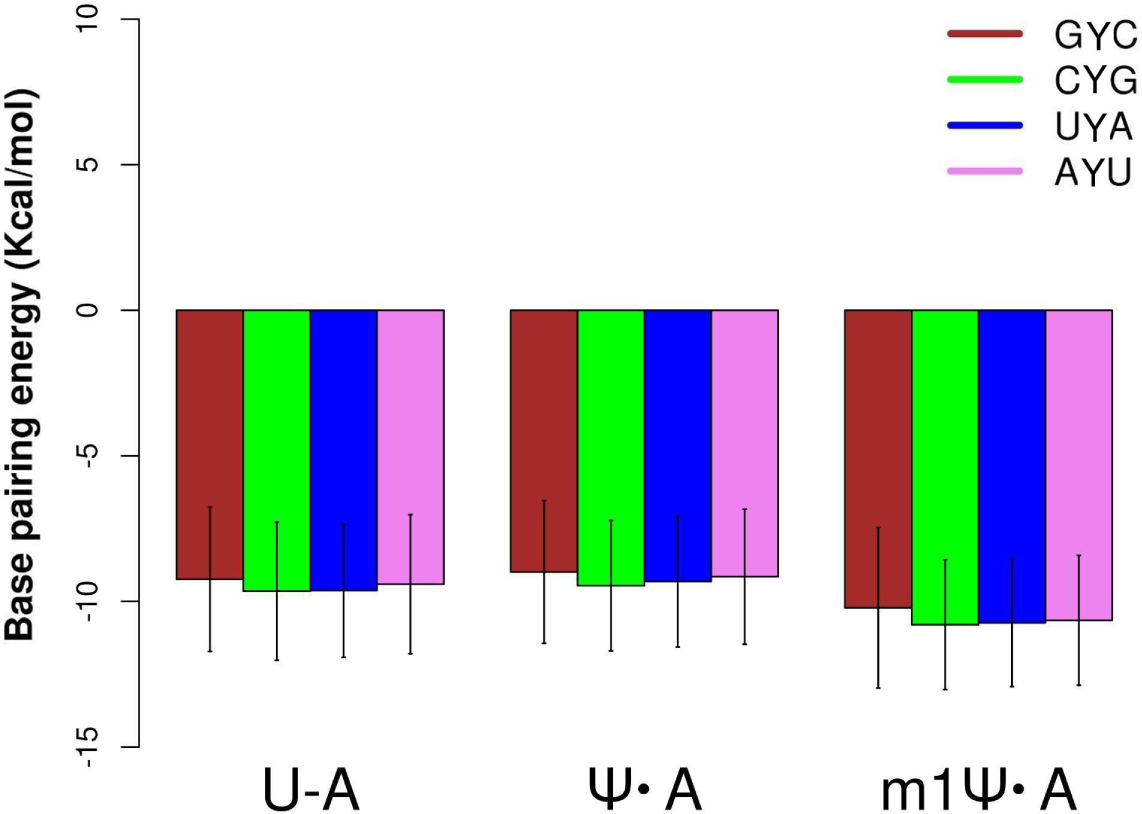
Comparison of the base pairing energies of the U-A, Ψ-A and m^1^Ψ-A pairs corresponding to the duplexes of four different contexts (Y in the figure legends indicates U, Ψ and m^1^Ψ). The values corresponding to the first set of simulations for m^1^Ψ-modified duplexes are plotted and the values for the U-A, Ψ-A pairs were obtained from Deb et al, ^5^.

**Figure 7.**
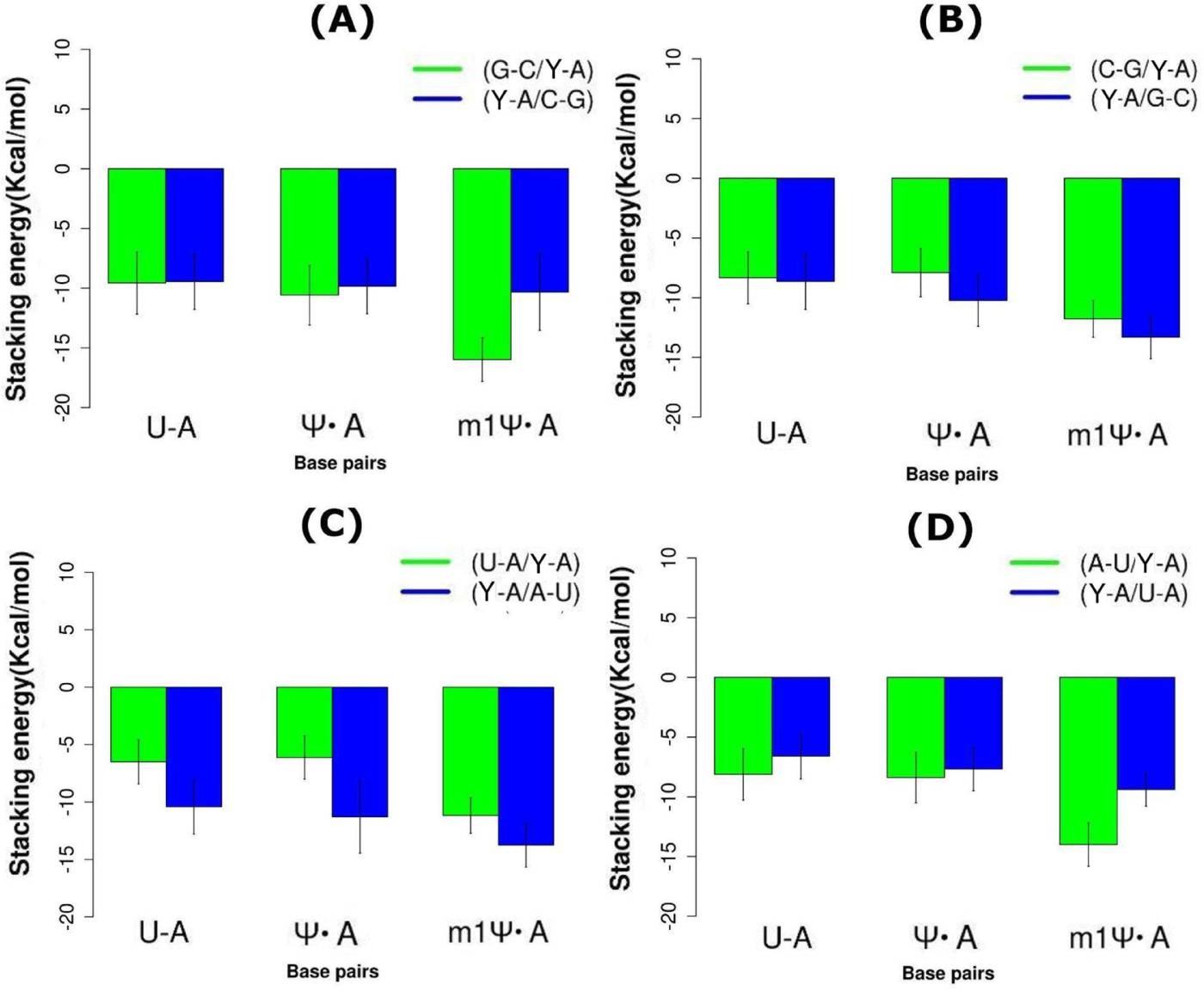
Comparison of the base pair stacking energies of the base pair steps containing U-A, Ψ-A and m^1^Ψ-A pairs respectively, corresponding to the duplexes of four different contexts (Y in the figure legends indicates U, Ψ and m^1^Ψ). The values corresponding to the first set of simulations for m^1^Ψ-modified duplexes are plotted and the values for the U-A, Ψ-A pairs were obtained from Deb et al, ^5^.

Comparison of the base pairing energies for the Y-A, Y-G, Y-U and Y-C pairs within duplex-GYC, revealed the following trend for the base pairs Y-G < Y-A< Y-U < Y-C (where Y indicated U/Ψ/m^1^Ψ) (Tables S16-22, Figures 8,S29). The stacking energies were observed to show the following trend 5’G-3’C/3’m^1^Ψ-5’G < 5’G-3’C/3’m^1^Ψ-5’A < 5’G-3’C/3’m^1^Ψ-5’C < 5’G-3’C/3’m^1^Ψ-5’U < 5’m^1^Ψ-3’A/3’C-5’G < 5’m^1^Ψ-3’C/3’C-5’G < 5’m^1^Ψ-3’G/3’C-5’G ∼ 5’m^1^Ψ-3’U/3’C-5’G while for the control duplexes (duplex-GUC) we observed 5’G-3’C/3’U-5’G < 5’G-3’C/3’U-5’A < 5’U-3’A/3’C-5’G < 5’G-3’C/3’U-5’C < 5’U-3’C/3’C-5’G < 5’U-3’G/3’C-5’G < 5’G-3’C/3’U-5’U < 5’U-3’U/3’C-5’G (Tables S16-22, Figure S30). For the Ψ modified duplexes (duplex-GΨC), the stacking energies showed the following trend: 5’G-3’C/3’Ψ-5’A ∼ 5’G-3’C/3’Ψ-5’G < 5’Ψ-3’A/3’C-5’G < 5’G-3’C/3’Ψ-5’C < 5’Ψ-3’C/3’C-5’G < 5’Ψ-3’G/3’C-5’G <

**Figure 8.**
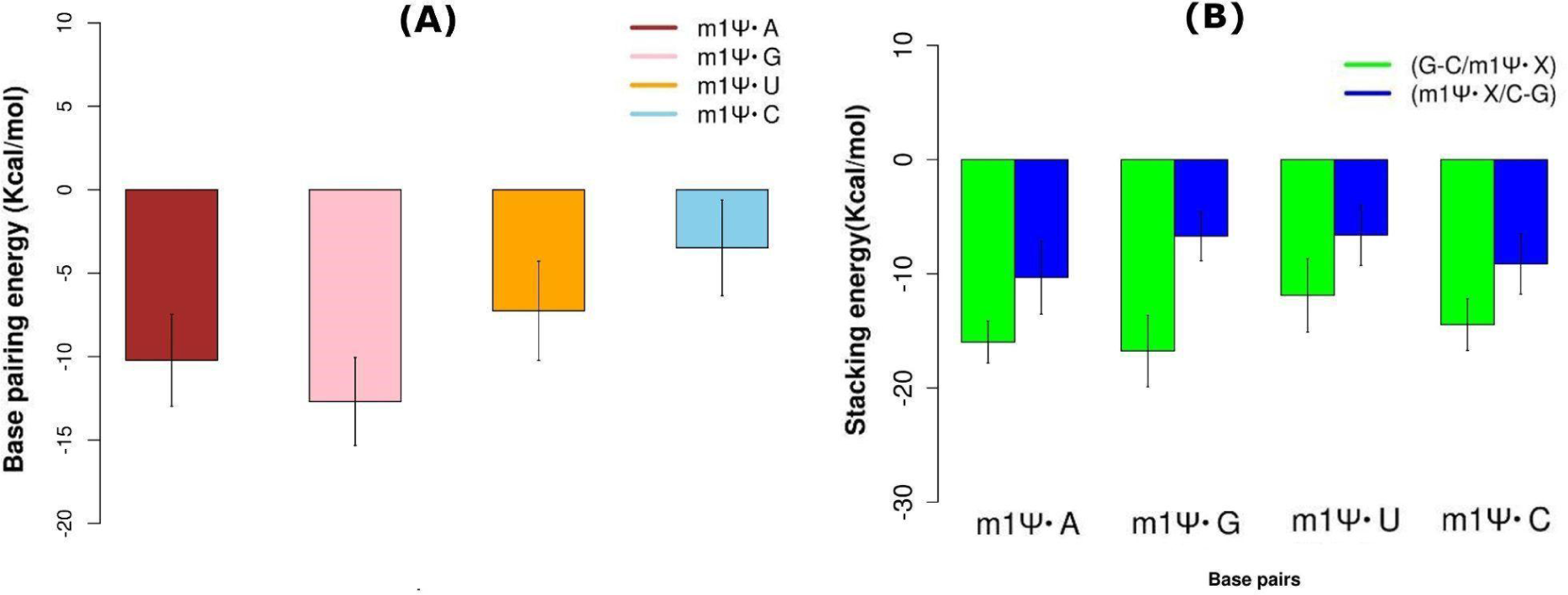
Comparison of the (A) base pairing energies of the m^1^Ψ-A pair and m^1^Ψ-G, m^1^Ψ-U and m^1^Ψ-C mismatches and (B) base pair stacking energies of the base pair steps containing the m^1^Ψ-A pair and m^1^Ψ-G, m^1^Ψ-U and m^1^Ψ-C mismatches within Duplex-Gm^1^ΨC for the first set of simulations (X in the figure legends indicates A/U/G/C).

5’Ψ-3’U/3’C-5’G < 5’G-3’C/3’Ψ-5’U (Tables S16-22, Figure S30). The base pairing energies and the base stacking energies were observed to be predominantly influenced by the electrostatic and the van der Waals interactions respectively.

##### Free energies of duplex formation

The calculation of the free energies of duplex formation (using MM-GBSA) revealed that the inclusion of the m^1^Ψ modification resulted in more stable duplexes for m^1^Ψ-A, m^1^Ψ-U and m^1^Ψ-C pairs than the control and Ψ-modified duplexes for each set of simulations (Tables S23-26). But, the duplex containing m^1^Ψ-G pair was found to be less stable compared to the control and Ψ-modified duplexes in the first set of simulations. But in the second set of simulations, Duplex-Gm^1^ΨC(m^1^Ψ-G) was more stable compared to Duplex-GΨC(Ψ-G) and Duplex-GUC(U-G). The changes in the free energies of duplex formation upon inclusion of the Ψ modification were observed to show a similar trend as was observed in the experiment ^4^ for the Ψ-A, Ψ-G and Ψ-U pairs (Table S24,27). But interestingly, the duplex containing Ψ-C pair was observed to be much more stable compared to the corresponding control duplex containing U-C pair. In general, the presence of the Ψ and m^1^Ψ modification was observed to stabilize the duplexes compared to the respective control duplexes.

There has been a lot of work connected with the mechanism of stabilization/destabilization of RNA structures as a result of pseudouridylation ^3–6,35–37^. The postulated mechanisms include stabilization through bridging water molecules that are not observed in the presence of uridines. Additionally, stacking is considered to be an important factor in determining the change in stabilities as well as the preference for C3’-endo conformation ^5,35–36^. In an earlier work from our group, NMR studies along with extensive quantum chemical modeling and molecular dynamics simulations showed that the additional hydration site present in pseudouridine contributed to the stabilization of the Ψ-modified duplexes over unmodified duplexes. In the context of stacking interactions, our results revealed the context-dependent stabilization of the base stacking energies for Ψ-modified duplexes compared to their unmodified counterparts and the stacking interactions were observed to be the key factor in the sequence context dependence of the increased stability pseudouridylated RNA ^5^.

In the present work, we have extended our investigation into the therapeutically important pseudouridine derivative m^1^Ψ. As has been previously observed in different experimental studies, N1-methylpseudouridine generally also has a stabilizing effect on RNA structure apart from its immensely intriguing biological effect of low immunogenicity. The question that needs to be answered is how the hydration, stacking and conformational factors differ for the derivative in comparison with those of pseudouridine. From our simulations, we observed that m^1^Ψ can form pairs with nucleobases other than adenine. Although, m^1^Ψ does not contain an additional hydration site unlike Ψ, we observed context-dependent stabilization of the duplexes with m^1^Ψ compared to Ψ and U, particularly in terms of stacking interactions. Our observation of the stabilization of the base stacking interactions by the N1-methylation of pseudouridine is consistent with the earlier reports suggesting the context-dependent destabilization of DNA (duplex), particularly the stacking interactions upon substitution of T (thymidine) with dU (deoxyuridine)^67^ and the enhancement of base stacking upon C5-methylation of cytosine^68^. The improved stacking observed with m^1^Ψ compared to Ψ might be due to the increased molecular polarizability resulting in the enhancement of the van der Waals interactions, as a consequence of the presence of the methyl group as is also reported for C5-methylation of pyrimidines^69^. Our quantum mechanical calculations (at B3LYP/6-311G(d) level of theory, using water environment by the PCM (Polarizable Continuum Model)) revealed that addition of methyl group at N1 position of pseudouridine nucleoside results in an increase in polarizability (data not shown). However, our results also suggest that the base pairs formed by m^1^Ψ with the canonical residues were additionally stabilized by inter-base water mediated interactions as was also observed for Ψ and U residues within similar duplexes.

Pseudouridine and its N1-methyl derivative are two key modifications used in mRNA therapeutics and are widely used in designing synthetic therapeutic mRNAs. Particularly, N1-methylpseudouridine has been used in the development of mRNA vaccines against COVID-19 disease ^32^ and there is an immensely growing interest for this modification in the field of mRNA therapeutics in the current years. Such mRNA vaccines containing m^1^Ψ have been reported to show high stability and efficacy over those containing the canonical counterpart ^32,70^. A recent study by Monroe et al ^71^ revealed the alteration of mRNA decoding as a result of Ψ and m^1^Ψ modifications during translation process and their study also suggested the context-dependent modulation of codon:anticodon base pairing interactions by Ψ and m^1^Ψ. Our study suggests the ability of m^1^Ψ to form more stable base pairs with the canonical residues compared to U or Ψ. Hence, the presence of this modification might result in enhanced stability of synthetic mRNA transcripts. It can also modulate the mRNA-tRNA interactions and hence the coding and translation efficiency.

## CONCLUSION

In the present study, we aimed to theoretically investigate the consequences of the presence of Pseudouridine (Ψ) and N1-methylpseudouridine in RNA duplexes, two commonly found post-transcriptional modifications which have been reported to play pivotal roles in the stability, dynamics, folding and functions of RNA. Both Ψ and m^1^Ψ have been previously reported to stabilize RNA duplexes over their canonical counterpart uridine ^3–5,38^. In this study, we have meticulously analyzed and re-examined the effect of these modifications in the structure, hydration pattern, dynamics and energetics of RNA duplexes to obtain a more comprehensive knowledge on the roles of these modifications.

Stabilization of C3’-endo sugar pucker conformation and water mediated interactions coordinated by the additional H-bond donor in Ψ stabilizing the interactions between the backbone atoms resulting in the enhancement of base stacking interactions have been proposed to be the the key mechanisms for the stabilization of RNA by Ψ ^1–7^. The stabilizing effect of it’s N1-methyl derivative (m^1^Ψ) in RNA has also been reported ^38^, but the mechanism of stabilization by this modification requires thorough investigation.

In the current work, our observations suggest that the presence of Ψ as well as m^1^Ψ resulted in more stable base stacking and base pairing interactions within the RNA duplexes under this study. The analysis of the hydration pattern revealed the formation of intra-residue water bridge between the OP2 and HN1 atoms of Ψ and inter-residue water bridge between the OP2 and HN1 atoms of Ψ and the OP2 atom of its 5’ nucleotide coordinating a single water molecule or by a chain of two water molecules within the Ψ-modified duplexes. This additional hydration site in Ψ contributes to the stabilization of the duplexes by the improvement of the interactions between the phosphodiester backbone atoms and hence also resulting in enhanced base stacking interactions over its canonical counterpart. Although m^1^Ψ lacks this additional hydration site, significant water bridging interactions between the backbone phosphate atoms were observed within the m^1^Ψ-modified duplexes. Energetically, m^1^Ψ was observed to form more stable base pair with A compared to Ψ and U in all sequence contexts but the base pairing energy values varied depending on the sequence context. m^1^Ψ was also observed to form more stable base pairs with G, U and C within the Gm^1^ΨC context. In general, m^1^Ψ was found to improve the base stacking interactions over U and Ψ for the duplexes in this study.

Overall, the results from our study indicate the context-dependent stabilization of RNA duplexes in presence of m^1^Ψ. Our results comprising the details on the structural and thermodynamic consequences of this modification on RNA duplexes and the trends in the context-dependent stabilities might be useful in understanding the stability and folding properties of RNA structures containing this modification and might also aid in the accurate design of m^1^Ψ-modified synthetic therapeutic mRNAs. In our future work, we would extend our investigation to evaluate the effect of this modification in other structural contexts e.g. stem-loop, apical and internal-loop motifs using integrated experimental and computational approaches to better explore the role of this modification in the stability of RNA.

## ASSOCIATED CONTENT

### Supporting Information

#### METHODS

MD simulations of unmodified and Ψ-modified duplexes. Table S1. Average and standard deviations of Root mean squared deviation and radius of gyration for the duplexes in this study. Table S2. Average and standard deviations of inter-strand C1’-C1’ distances (Å) and λ angles (for the 5th and 14th residues) of the duplexes. Table S3. Fraction (in %) of NORTH sugar pucker and ANTI base orientation of the 5th and 14th residues of the duplexes. Tables S4-5. Average values and standard deviations of local base-pair parameters for m^1^Ψ-modified duplexes. Table S6-7. Average values and standard deviations of local base-pair step parameters for m^1^Ψ-modified duplexes. Table S8. Average values and standard deviations of local base-pair parameters for unmodified and Ψ-modified duplexes. Table S9. Average values and standard deviations of local base-pair step parameters for unmodified and Ψ-modified duplexes. Tables S10-11. Frequencies (in %) of hydrogen bonds between the 5th and 14th residues for the duplexes in this study. Table S12-13. Water bridges (occupancy in %) formed by the backbone phosphate oxygen atoms of m^1^Ψ(5), U(5) and Ψ(5) with the phosphate oxygen atoms of its immediate neighboring residues for the duplexes. Table S14. Water bridging interaction between the bases of 5th and 14th residues in the duplexes (containing mismatches) in this study. Tables S15-22. Interaction energies of the base pairs and base-pair steps for the studied duplexes. Tables S23-27 Average and standard deviations of MM-GBSA binding free energies for the studied duplexes. Figure S1. Observed geometries of the U-G, U-U and U-C, Ψ-G, Ψ-U and Ψ-C mismatches in the studied duplexes. Figure S2-3. Time evolution plots of the root-mean-square deviations (heavy atoms) of the studied duplexes. Figure S4. Time evolution of λ angles for the 5th and 14th residues for the duplexes (containing mismatches) in this study. Figure S5. Population distribution of glycosidic torsion angle (χ) for m^1^Ψ(5) in the studied duplexes. Figures S6. Population distribution of glycosidic torsion angle (χ) for U(5) and Ψ(5) in the simulated duplexes. Figures S7-12. Plots of opening of the central 7 base pairs and shift of the central 6 base pair steps for the studied suplexes. Figures S13-14. Water occupancy maps for the duplexes (residues 4-6) for the studied duplexes. Figure S15. Snapshots of water bridges formed between the HN1 and OP2 atoms of Ψ(5) and the OP2 atom of G(4) for the Ψ-modified duplexes in this study. Figures S16-17. RDF of water oxygen atoms around the H20 (methyl hydrogen) atom and the C10 (methyl carbon) atom of m^1^Ψ(5) for the studied duplexes. Figure S18. RDF of water oxygen atoms around the H5 atom of U(5) and that about the HN1 atom of Ψ(5) for the studied duplexes. Figure S19. RDF of water oxygen atoms around geometric center of the H5 and OP2 atoms of U(5) and that about the geometric center of the HN1 and OP2 atoms of Ψ(5) for the studied duplexes. Figure S20-25. Snapshots of the base pairs (formed by the 5th and 14th residues) for the unmodified and Ψ-modified duplexes (containing mismatches) depicting water mediated interactions between the bases. Figure S28. Water occupancy maps for the mismatches under this study.

## DATA AND SOFTWARE AVAILABILITY

All data and software are available. Additional data which support the findings in this study are available upon request from the corresponding author. The software and web servers used in this study are as follows: AMBER18: http://ambermd.org/. Pymol: https://pymol.org/2/. ACD/ChemSketch (Freeware version): https://www.acdlabs.com/. Curves+/Canal http://gbio-pbil.ibcp.fr/tools/curves_plus/. UCSF chimera https://www.cgl.ucsf.edu/chimera/. R 3.5.0: https://cran.r-project.org/.

## AUTHOR INFORMATION

### Author Contributions

N.D., and A.L. jointly conceived and designed the study. N.D. and I.D. generated the data. N.D. analyzed the results. All authors reviewed the manuscript.

### Notes

The authors declare no competing financial interest.

## ACKNOWLEDGEMENTS

N.D. acknowledges support from the DST INSPIRE Senior Research Fellowship (DST/ INSPIRE Fellowship/2018/IF180895). The simulations and calculations were performed at the Poznan Supercomputing and Networking Center.

## Notes

### Competing Interest Statement

The authors have declared no competing interest.

