## Supplemental data for "Structural and thermodynamic consequences of base pairs containing pseudouridine and N1-methylpseudouridine in RNA duplexes"

**Author Affiliations**

**[*]Correspondence to**

ORCID of the authors: Nivedita Dutta (0000-0002-8371-9007), Indrajit Deb (0000-0002-2722-582X) Joanna Sarzynska (0000-0002-0500-2238), Ansuman Lahiri (0000-0002-7398-4114)

METHODS

***Molecular Dynamics (MD) simulations***

*Solvation, neutralization and ionization for the duplexes 8-13*

For the control duplexes (containing U-G, U-U and U-C pairs) and for the the Ψ-modified duplexes (containing Ψ-G, Ψ-U and Ψ-C pairs), the initial structures were truncated octahedral boxes with the minimum distance of 10 Å between any atom of the solute and the edge of the periodic box and the systems were then neutralized using Na+ ions. Additional $Na^+$ and $Cl^-$ ions were added to the solvated and neutralized systems (The Joung and Cheatham ion parameters [1] were used for addition of ions to achieve the salt concentration at 1.0 M to replicate the environment of UV melting experiments as reported in Kierzek et al, [2]). The additional Na+ and Cl- ions were swapped with random water residues which were not closer than 6.0 Å from any nucleotide residue and were not closer than 4.0 Å from any of the other $Na^+$/$Cl^-$ ions.

*Minimization, equilibration and production for the duplexes 8-13*

The solvated, neutralized and ionized systems were energy minimized by 200 steps of steepest descent followed by 300 steps conjugate gradient using a weak RMS force convergence criteria of 0.1 kcal/mol for the energy gradient. In this step, the duplexes were held by a positional restraint force of 25.0 kcal/mol/Å$^2$. Next, the solvent molecules were equilibrated holding the duplexes by a positional restraint force of 25.0 kcal/mol/Å$^2$ and by performing a constant pressure Langevin dynamics using a collision frequency of 1 ps$^{-1}$ for 50 ps at 300 K temperature. After that, two steps of energy minimizations each consisting of 500 steps of steepest descent followed by 2000 steps conjugate gradient minimizations, with the default RMS force convergence criteria of 0.00010 kcal/mol and a positional restraint force of 25.0 kcal/mol/Å$^2$ was applied to the duplexes in the first step. Next, three steps of equilibration were carried out before the production run. In the first step, the systems were slowly heated from 0 K to 300 K using constant volume Langevin dynamics in 80 ps with a collision frequency of 5 ps$^{-1}$ and the duplexes were held fixed with a restraint force of 15 kcal/mol/Å$^2$. For the next 20 ps, the temperature was fixed at a constant temperature of 300 K. After which, a 100 ps equilibration run was performed at 300 K by constant pressure Langevin dynamics with a collision frequency of 1 ps$^{-1}$, holding the duplexes with a 10 kcal/mol/Å$^2$ restraint force. In the final equilibration step, a 1 ns equilibration run was carried out at 300 K, keeping the conditions same as the of previous equilibration steps but removing the positional restraint on the duplexes. In this step,the inter-strand Watson-Crick hydrogen bonding distance restraints to the terminal base pairs to avoid fraying ends according to Saenger [3] allowing 0.1 Å movement from the equilibrium bond distance.

For the production run stage, the conditions were similar to the final equilibration step and the. trajectory files were written after every 10 ps for each set of simulations. The production MD run generated a total simulation of 500 ns for each system. Berendsen barostat was used to perform constant pressure simulations by turning on isotropic position scaling with a reference pressure of 1 atm and a pressure relaxation time of 1ps. The nonbonded interactions within 8.0 Å long-range cutoff were considered during the minimization, equilibration and production run steps. The particle mesh Ewald (PME) method was used for handling long-range interactions during the simulations. The molecular dynamics simulations were carried out using the AMBER14 software package[4] using the GPU version of the pmemd module.

**Table S1.** Average values and standard deviations (in parentheses) of the mass-weighted root-mean-square deviations (RMSDs) and radius of gyrations ($R_g$s) for the heavy atoms within the duplexes in this study.

| **Duplexes** | **RMSD (from minimized structure) (Å)[a]** | **$R_g$ (Å)[c]** |
|---|---|---|
| **Duplex-Gm$^1$ΨC (m$^1$Ψ-A)** | r1 1.26 (0.34)<br>r2 1.26 (0.33) | r1 11.26 (0.29)<br>r2 11.24 (0.28) |
| **Duplex-Cm$^1$ΨG (m$^1$Ψ-A)** | r1 1.21 (0.32)<br>r2 1.20 (0.32) | r1 11.16 (0.29)<br>r2 11.15 (0.29) |
| **Duplex-Um$^1$ΨA (m$^1$Ψ-A)** | r1 1.49 (0.32)<br>r2 1.49 (0.32) | r1 11.14 (0.28)<br>r2 11.13 (0.27) |
| **Duplex-Am$^1$ΨU (m$^1$Ψ-A)** | r1 1.42 (0.33)<br>r2 1.39 (0.29) | r1 11.17 (0.27)<br>r1 11.13 (0.25) |
| **Duplex-Gm$^1$ΨC (m$^1$Ψ-G)** | r1 1.45 (0.37)<br>r2 1.25 (0.33) | r1 11.17 (0.31)<br>r2 11.14 (0.29) |
| **Duplex-Gm$^1$ΨC (m$^1$Ψ-U)** | r1 1.78 (0.32)<br>r2 1.79 (0.32) | r1 11.13 (0.31)<br>r2 11.12 (0.29) |
| **Duplex-Gm$^1$ΨC (m$^1$Ψ-C)** | r1 1.74 (0.32)<br>r2 1.78 (0.38) | r1 11.11 (0.28)<br>r2 11.06 (0.29) |
| **Duplex-GΨC (Ψ-G)** | 1.35 (0.25) | 10.93 (0.27) |
| **Duplex-GΨC (Ψ-U)** | 1.52 (0.23) | 10.89 (0.26) |

| | | |
|---|---|---|
| **Duplex-GΨC (Ψ-C)** | 1.44 (0.28) | 10.86 (0.27) |
| **Duplex-GUC (U-G)** | 1.36 (0.28) | 10.92 (0.28) |
| **Duplex-GUC (U-U)** | 1.46 (0.27) | 10.85 (0.27) |
| **Duplex-GUC (U-C)** | 1.76 (0.56) | 10.65 (0.39) |

[a]RMSDs corresponding to the heavy atoms, calculated over the last 400 ns of the trajectories in reference to the energy minimized structure.
[b]RMSDs corresponding to the heavy atoms, calculated over the 400 ns of the trajectories in reference to the centroid structure of the most populated cluster.
[c]$R_g$ (radius of gyration) corresponding to the heavy atoms, calculated over the last 400 ns of the trajectories.
r1 and r2 indicate values obtainedfrom replicate 1 and replicate 2 respectively.

**Table S2.** Average values and standard deviations (in parentheses) (A-C) of inter-strand C1'-C1' distances (Å) and (D) that of λ angles (for the 5th and 14th residues).

(A) For the duplexes containing $m^1\Psi$

| Base pair | Duplex-Gm1ΨC ($m^1\Psi$-A) | Duplex-Cm1ΨG ($m^1\Psi$-A) | Duplex-Um1ΨA ($m^1\Psi$-A) | Duplex-Am1ΨU ($m^1\Psi$-A) | Duplex-Gm1ΨC ($m^1\Psi$-G) | Duplex-Gm1ΨC ($m^1\Psi$-U) | Duplex-Gm1ΨC ($m^1\Psi$-C) |
|---|---|---|---|---|---|---|---|
| U1-A18 | r1 10.55 (0.27)<br>r2 10.56 (0.30) | r1 10.57 (0.26)<br>r2 10.57 (0.25) | r1 10.57 (0.26)<br>r2 10.56 (0.27) | r1 10.55 (0.26)<br>r2 10.55 (0.27) | r1 10.57 (0.26)<br>r2 10.56 (0.26) | r1 10.52 (0.29)<br>r1 10.53 (0.29) | r1 10.55 (0.27)<br>r2 10.54 (0.29) |
| C2-G17 | r1 10.71 (0.18)<br>r2 10.71 (0.22) | r1 10.69 (0.18)<br>r2 10.69 (0.18) | r1 10.69 (0.18)<br>r2 10.69 (0.19) | r1 10.71 (0.18)<br>r2 10.71 (0.18) | r1 10.63 (0.20)<br>r2 10.71 (0.17) | r1 10.71 (0.17)<br>r2 10.70 (0.18) | r1 10.71 (0.17)<br>r2 10.67 (0.19) |
| G3-C16/ C3-G16/ A3-U16/ U3-A16 | r1 10.67 (0.32)<br>r2 10.68 (0.33) | r1 10.65 (0.32)<br>r2 10.63 (0.30) | r1 10.61 (0.31)<br>r2 10.61 (0.31) | r1 10.69 (0.29)<br>r2 10.68 (0.29) | r1 10.90 (0.33)<br>r2 10.63 (0.30) | r1 10.70 (0.32)<br>r2 10.76 (0.33) | r1 10.65 (0.31)<br>r2 10.76 (0.34) |
| G4-C15 | r1 10.70 (0.16) | r1 10.72 (0.16) | r1 10.63 (0.29) | r1 10.66 (0.32) | r1 10.72 (0.17) | r1 10.64 (0.19) | r1 10.70 (0.18) |

| | | | | | | | |
|---|---|---|---|---|---|---|---|
| | r2 10.70 (0.20) | r2 10.71 (0.16) | r2 10.62 (0.29) | r2 10.64 (0.32) | r2 10.73 (0.17) | r2 10.64 (0.18) | r2 10.69 (0.18) |
| m[1]Ψ5-X14 X=>(A/G/U/C) | r1 **10.77 (0.29)**<br><br>r2 **10.77 (0.29)** | r1 **10.84 (0.28)**<br><br>r2 **10.84 (0.28)** | r1 **10.81 (0.27)**<br><br>r2 **10.83 (0.27)** | r1 **10.77 (0.29)**<br><br>r2 **10.77 (0.29)** | r1 **10.58 (0.34)**<br><br>r2 **10.69 (0.25)** | r1 **9.82 (0.76)**<br><br>r2 **9.89 (0.74)**<br><br>**C1**<br>r1 **9.69 (0.67)**<br><br>r2 **9.81 (0.71)**<br><br>**C2**<br>r1 **9.71 (0.71)**<br><br>r2 **9.81 (0.69)** | r1 **11.09 (0.84)**<br><br>r2 **11.03 (0.96)**<br><br>**C1**<br>r1 **10.69 (0.47)**<br><br>r2 **10.62 (0.46)**<br><br>**C2**<br>r1 **11.86 (0.37)**<br><br>r2 **11.83 (0.34)** |
| C6-G13/ G6-C13/ U6-A13/ A6-U13 | r1 10.74 (0.17)<br><br>r2 10.74 (0.17) | r1 10.76 (0.17)<br><br>r2 10.76 (0.17) | r1 10.69 (0.28)<br><br>r2 10.68 (0.28) | r1 10.64 (0.28)<br><br>r2 10.65 (0.27) | r1 10.70 (0.19)<br><br>r2 10.74 (0.16) | r1 10.62 (0.21)<br><br>r2 10.62 (0.22) | r1 10.69 (0.23)<br><br>r2 10.66 (0.30) |
| A7-U12 | r1 10.66 (0.31)<br><br>r2 10.66 (0.31) | r1 10.55 (0.28)<br><br>r2 10.56 (0.28) | r1 10.62 (0.29)<br><br>r2 10.63 (0.29) | r1 10.66 (0.32)<br><br>r2 10.63 (0.28) | r1 10.69 (0.33)<br><br>r2 10.71 (0.30) | r1 10.70 (0.31)<br><br>r2 10.69 (0.32) | r1 10.69 (0.32)<br><br>r2 10.67 (0.32) |
| G8-C11 | r1 10.65 (0.19)<br><br>r2 10.68 (0.19) | r1 10.64 (0.19)<br><br>r2 10.65 (0.19) | r1 10.65 (0.18)<br><br>r2 10.64 (0.18) | r1 10.65 (0.18)<br><br>r2 10.66 (0.18) | r1 10.61 (0.23)<br><br>r2 10.65 (0.18) | r1 10.66 (0.18)<br><br>r2 10.66 (0.18) | r1 10.65 (0.19)<br><br>r2 10.65 (0.19) |
| U9-A10 | r1 10.63 (0.27)<br><br>r2 10.58 (0.28) | r1 10.65 (0.27)<br><br>r2 10.65 (0.27) | r1 10.63 (0.27)<br><br>r2 10.64 (0.27) | r1 10.64 (0.27)<br><br>r2 10.64 (0.27) | r1 10.65 (0.27)<br><br>r2 10.62 (0.28) | r1 10.62 (0.28)<br><br>r2 10.59 (0.28) | r1 10.62 (0.27)<br><br>r2 10.62 (0.27) |
| C1 and C2 (in blue) indicate the values for conformers within the most populated cluster and for those | | | | | | | |

| |
|---|
| within the second most populated cluster. r1 and r2 indicate values obtained from replicate 1 and replicate 2 respectively. |

(B) For the duplexes containing U and Ψ

| Base pair | **Duplex-GUC (U-G)** | **Duplex-GUC (U-U)** | **Duplex-GUC (U-C)** | **Duplex-GΨC (Ψ-G)** | **Duplex-GΨC (Ψ-U)** | **Duplex-GΨC (Ψ-C)** |
|---|---|---|---|---|---|---|
| U1-A18 | 10.58 (0.25) | 10.55 (0.27) | 10.55 (0.28) | 10.57 (0.26) | 10.55 (0.26) | 10.57 (0.26) |
| C2-G17 | 10.69 (0.19) | 10.72 (0.18) | 10.69 (0.18) | 10.72 (0.17) | 10.69 (0.19) | 10.71 (0.17) |
| G3-C16/ C3-G16/ A3-U16/ U3-A16 | 10.74 (0.33) | 10.72 (0.30) | 10.67 (0.29) | 10.64 (0.29) | 10.81 (0.34) | 10.65 (0.29) |
| G4-C15 | 10.72 (0.16) | 10.64 (0.19) | 10.70 (0.19) | 10.74 (0.16) | 10.64 (0.19) | 10.69 (0.19) |
| Y5-X14 Y=> (U/Ψ) X=>(G/U/C) | **10.58 (0.21)** | **9.98 (0.69)** <br> **C1 9.87 (0.63) C2 9.91 (0.66)** | **11.74 (1.45)** <br> **C1 12.23 (0.72) C2 10.65 (0.56)** | **10.68 (0.23)** | **9.77 (0.67)** <br> **C1 9.72 (0.60) C2 9.61 (0.55)** | **10.89 (0.78)** <br> **C1 10.65 (0.47) C2 11.27 (0.93)** |
| C6-G13/ G6-C13/ U6-A13/ A6-U13 | 10.74 (0.16) | 10.65 (0.18) | 10.73 (0.18) | 10.74 (0.16) | 10.63 (0.19) | 10.69 (0.18) |
| A7-U12 | 10.74 (0.31) | 10.74 (0.31) | 10.68 (0.32) | 10.73 (0.30) | 10.74 (0.31) | 10.73 (0.32) |
| G8-C11 | 10.65 (0.19) | 10.66 (0.18) | 10.64 (0.19) | 10.66 (0.18) | 10.67 (0.18) | 10.66 (0.18) |
| U9-A10 | 10.63 (0.27) | 10.62 (0.27) | 10.63 (0.27) | 10.63 (0.27) | 10.62 (0.27) | 10.63 (0.27) |
| C1 and C2 (in blue) indicate the values for conformers within the most populated cluster and for those | | | | | | |

| |
|---|
| within the second most populated cluster. |

(C) Average values and standard deviations (in parentheses) of λ angles for Y(5) (U/Ψ/m$^{1}$Ψ) and X14(A/U/G/C) for the duplexes under this study

| | | |
|---|---|---|
| | **m$^{1}$Ψ(5)**<br>**(C1'(A)-C1'(m$^{1}$Ψ)-C5(m$^{1}$Ψ))** | **A(14)**<br>**(C1'(m$^{1}$Ψ)-C1'(A)-N9(A))** |
| **Duplex-Gm$^{1}$ΨC (m$^{1}$Ψ-A)** | r1 54.06 (6.82)<br>r2 52.89 (4.37) | r1 56.36 (6.28)<br>r2 57.26 (4.27) |
| **Duplex-Cm$^{1}$ΨG (m$^{1}$Ψ-A)** | r1 52.17 (4.20)<br>r2 52.11 (4.22) | r1 56.39 (3.91)<br>r2 56.32 (3.89) |
| **Duplex-Um$^{1}$ΨA (m$^{1}$Ψ-A)** | r1 52.33 (3.91)<br>r2 52.08 (3.99) | r1 56.92 (3.74)<br>r2 56.76 (3.76) |
| **Duplex-Am$^{1}$ΨU (m$^{1}$Ψ-A)** | r1 52.56 (4.39)<br>r2 52.51 (4.49) | r1 57.15 (3.89)<br>r2 57.15 (3.87) |
| **Duplex-Gm$^{1}$ΨC (m$^{1}$Ψ-G)** | **m$^{1}$Ψ(5)**<br>**(C1'(G)-C1'(m$^{1}$Ψ)-C5(m$^{1}$Ψ))** | **G(14)**<br>**(C1'(m$^{1}$Ψ)-C1'(G)-N9(G))** |
| | r1 71.14 (10.07)<br>r2 67.74 (7.14) | r1 46.92 (7.39)<br>r2 44.68 (5.94) |
| **Duplex-Gm$^{1}$ΨC (m$^{1}$Ψ-U)** | **m$^{1}$Ψ(5)**<br>**(C1'(U)-C1'(m$^{1}$Ψ)-C5(m$^{1}$Ψ))** | **U(14)**<br>**(C1'(m$^{1}$Ψ)-C1'(U)-N1(U))** |
| | r1 54.09 (15.59)<br>r2 51.78 (18.28) | r1 52.44 (16.19)<br>r2 54.22 (16.78) |
| Most populated cluster | r1 66.62 (6.90)<br>r2 66.22 (7.19) | r1 39.32 (6.60)<br>r2 38.60 (6.48) |
| Second Most populated cluster | r1 36.36 (6.20)<br>r2 35.85 (6.87) | r1 71.28 (6.19)<br>r2 70.00 (6.37) |
| **Duplex-Gm$^{1}$ΨC (m$^{1}$Ψ-C)** | **m$^{1}$Ψ(5)**<br>**(C1'(C)-C1'(m$^{1}$Ψ)-C5(m$^{1}$Ψ))** | **C(14)**<br>**(C1'(m$^{1}$Ψ)-C1'(C)-N1(C))** |

| | | |
|---|---|---|
| | r1 54.80 (20.82)<br>r2 54.07 (21.80) | r1 36.96 (13.12)<br>r2 37.33 (12.63) |
| Most populated cluster | r1 75.36 (6.81)<br>r2 76.21 (6.75) | r1 26.46 (5.44)<br>r2 27.32 (5.56) |
| Second Most populated cluster | r1 33.22 (6.64)<br>r2 33.84 (5.54) | r1 40.94 (5.57)<br>r2 41.35 (4.82) |
| **Duplex-GUC (U-G)** | **U(5)**<br>**(C1'(G)-C1'(U)-N1(U))** | **G(14)**<br>**(C1'(U)-C1'(G)-N9(G))** |
| | 68.70 (4.51) | 43.73 (4.43) |
| **Duplex-GUC (U-U)** | **U(5)**<br>**(C1'(U14)-C1'(U5)-N1(U5))** | **U(14)**<br>**(C1'(U5)-C1'(U14)-N1(U14))** |
| | 51.42 (16.79) | 53.21 (17.26) |
| Most populated cluster | 36.76 (6.13) | 68.04 (5.65) |
| Second Most populated cluster | 67.31 (5.93) | 36.14 (6.57) |
| **Duplex-GUC (U-C)** | **U(5)**<br>**(C1'(C)-C1'(U)-N1(U))** | **C(14)**<br>**(C1'(U)-C1'(C)-N1(C))** |
| | 47.85 (25.33) | 34.89 (16.46) |
| Most populated cluster | 26.13 (11.38) | 33.28 (11.66) |
| Second Most populated cluster | 72.64 (9.03) | 31.61 (15.10) |
| **Duplex-GΨC (Ψ-G)** | **Ψ(5)**<br>**(C1'(G)-C1'(Ψ)-C5(Ψ))** | **G(14)**<br>**(C1'(Ψ)-C1'(G)-N9(G))** |
| | 68.10 (5.68) | 43.94 (5.13) |
| **Duplex-GΨC (Ψ-U)** | **Ψ(5)**<br>**(C1'(U)-C1'(Ψ)-C5(Ψ))** | **U(14)**<br>**(C1'(Ψ)-C1'(U)-N9(U))** |
| | 59.14 (14.99) | 46.83 (15.01) |
| Most populated cluster | 68.07 (5.43) | 38.64 (5.96) |
| Second Most populated cluster | 37.59 (5.69) | 69.04 (5.24) |
| **Duplex-GΨC (Ψ-C)** | **Ψ(5)** | **C(14)** |

| | (C1'(C)-C1'(Ψ)-C5(Ψ)) | (C1'(Ψ)-C1'(C)-N9(C)) |
|---|---|---|
| | 58.57 (20.01) | 36.48 (15.92) |
| Most populated cluster | 74.92 (5.73) | 26.44 (9.32) |
| Second Most populated cluster | 38.14 (7.84) | 44.15 (7.86) |
| r1 and r2 indicate values obtained from replicate 1 and replicate 2 respectively. | | |

**Table S3.** Fraction (in %) of NORTH sugar pucker and ANTI base orientation of the 5th and 14th residues in the duplexes (A) containing $m^1\Psi$-A pair; (B) containing $m^1\Psi$-G, $m^1\Psi$-U and $m^1\Psi$-C mismatches respectively; (C) containing U-G, U-U, U-C, Ψ-G, Ψ-U and Ψ-C mismatches respectively.

| **(A)** | **Duplex-Gm$^1$ΨC (m$^1$Ψ-A)** | | **Duplex-Cm$^1$ΨG (m$^1$Ψ-A)** | | **Duplex-Um$^1$ΨA (m$^1$Ψ-A)** | | **Duplex-Am$^1$ΨU (m$^1$Ψ-A)** | |
|---|---|---|---|---|---|---|---|---|
| Residues | **m$^1$Ψ5** | **A14** | **m$^1$Ψ5** | **A14** | **m$^1$Ψ5** | **A14** | **m$^1$Ψ5** | **A14** |
| **NORTH** | r1 100<br>r2 100 | r1 100<br>r2 100 | r1 100<br>r2 100 | r1 100<br>r2 100 | r1 100<br>r2 100 | r1 100<br>r2 100 | r1 100<br>r2 100 | r1 100<br>r2 100 |
| **ANTI** | r1 100<br>r2 99.97 | r1 99.94<br>r2 99.98 | r1 99.99<br>r2 99.99 | r1 99.98<br>r2 99.98 | r1 99.99<br>r2 99.98 | r1 99.98<br>r2 99.97 | r1 99.97<br>r2 99.98 | r1 99.98<br>r2 99.99 |
| r1 and r2 indicate values obtained from replicate 1 and replicate 2 respectively. | | | | | | | | |

| **(B)** | **Duplex-Gm$^1$ΨC (m$^1$Ψ-G)** | | **Duplex-Gm$^1$ΨC (m$^1$Ψ-U)** | | **Duplex-Gm$^1$ΨC (m$^1$Ψ-C)** | |
|---|---|---|---|---|---|---|
| Residues | **m$^1$Ψ5** | **G14** | **m$^1$Ψ5** | **U14** | **m$^1$Ψ5** | **C14** |
| **NORTH** | r1 100<br>r2 100 | r1 100<br>r2 100 | r1 99.99<br>r2 99.99 | r1 100<br>r2 100 | r1 100<br>r2 100 | r1 99.99<br>r2 100 |
| **ANTI** | r1 99.99<br>r2 99.83 | r1 99.64<br>r2 100 | r1 99.81<br>r2 99.98 | r1 99.99<br>r2 99.74 | r1 99.97<br>r2 99.98 | r1 99.98<br>r2 99.88 |
| r1 and r2 indicate values obtained from replicate 1 and replicate 2 respectively. | | | | | | |

| (C) | Duplex-GUC (U-G) | | Duplex-GUC (U-U) | | Duplex-GUC (U-C) | | Duplex-GΨC (Ψ-G) | | Duplex-GΨC (Ψ-U) | | Duplex-GΨC (Ψ-C) | |
|---|---|---|---|---|---|---|---|---|---|---|---|---|
| Residues | **U5** | **G14** | **U5** | **U14** | **U5** | **C14** | **Ψ5** | **G14** | **Ψ5** | **U14** | **Ψ5** | **C14** |
| **NORTH** | 99.99 | 100 | 100 | 99.99 | 98.69 | 95.61 | 100 | 100 | 100 | 99.99 | 100 | 100 |
| **ANTI** | 100 | 99.86 | 99.99 | 99.99 | 99.89 | 99.88 | 100 | 99.89 | 99.98 | 99.97 | 99.98 | 99.99 |

**Table S4.** Average values and standard deviations (in parentheses) of local base-pair parameters for (A) duplex-Gm$^{1}$ΨC, (B) duplex-Cm$^{1}$ΨG, (C) duplex-Um$^{1}$ΨA and (D) duplex-Am$^{1}$ΨU containing m$^{1}$Ψ-A pair.

(A) duplex-Gm$^{1}$ΨC

| Base pair | **Shear [Å]** | **Stretch [Å]** | **Stagger [Å]** | **Buckle [°]** | **Propeller [°]** | **Opening [°]** |
|---|---|---|---|---|---|---|
| U1-A18 | -0.19 (0.28)<br>-0.19 (0.30) | -0.03 (0.11)<br>-0.03 (0.22) | 0.12 (0.52)<br>0.11 (0.54) | -1.4 (15.4)<br>-1.1 (15.5) | -8.8 (10.8)<br>-8.6 11.2 | 2.7 (4.7)<br>2.7 (4.8) |
| C2-G17 | -0.01 (0.37)<br>-0.02 (0.36) | 0.04 (0.12)<br>0.04 (0.18) | -0.13 (0.41)<br>-0.12 (0.44) | 8.1 (11.0)<br>7.6 (11.1) | -14.1 (8.3 )<br>-13.7 (8.5) | 1.9 (4.0)<br>1.9 (3.7) |
| A3-U16 | 0.10 (0.31)<br>0.10 (0.36) | 0.03 (0.18)<br>0.03 (0.20) | -0.02 (0.45)<br>-0.01 (0.45) | 0.8 (9.2)<br>1.0 (9.2) | -12.7 (7.8)<br>-13.1 (7.8) | 2.6 (6.9)<br>2.6 (6.6) |
| G4-C15 | -0.04 (0.33)<br>-0.07 (0.73) | 0.02 (0.12)<br>0.02 (0.21) | 0.04 (0.41)<br>0.05 (0.42) | -2.0 (9.7)<br>-1.7 (9.5) | -12.4 (7.8)<br>-12.6 (7.8) | 1.9 (3.6)<br>2.0 (3.9) |
| m$^{1}$Ψ5-A14 | -0.02 (0.96)<br>-0.21 (0.31) | 0.01 (0.20)<br>0.02 (0.17) | 0.01 (0.48)<br>0.00 (0.48) | -1.1 (9.7)<br>0.0 (9.4) | -13.4 (7.3)<br>-13.7 (7.2) | 2.0 (6.5)<br>1.8 (6.2) |
| C6-G13 | -0.01 (0.35)<br>-0.01 (0.37) | 0.03 (0.12)<br>0.03 (0.14) | 0.09 (0.39)<br>0.07 (0.39) | 1.4 (9.1)<br>1.8 (9.3) | -10.5 (7.5)<br>-11.3 (7.6) | 1.5 (3.9)<br>1.6 (3.9) |
| A7-U12 | 0.11 (0.31)<br>0.10 (0.31) | 0.03 (0.17)<br>0.03 (0.17) | 0.04 (0.44)<br>0.04 (0.46) | 0.5 (9.3)<br>-1.2 (9.7) | -12.1 (7.9)<br>-13.6 (8.2) | 2.7 (6.5)<br>2.9 (6.7) |
| G8-C11 | -0.11 (0.40) | 0.02 (0.17) | 0.02 (0.44) | -1.3 (11.5) | -14.0 (9.5) | 2.8 (5.0) |

| | | | | | | |
|---|---|---|---|---|---|---|
| | -0.07 (0.40) | 0.02 (0.16) | 0.08 (0.45) | -3.4 (11.3) | -14.6 (9.5) | 2.4 (5.0) |
| U9-A10 | -0.16 (0.28)<br>-0.17 (0.28) | -0.04 (0.10)<br>-0.04 (0.11) | -0.16 (0.51)<br>-0.18 (0.51) | 1.3 (13.9)<br>2.2 (15.5) | -9.6 (11.8)<br>-6.5 (12.7) | 1.4 (5.1)<br>2.0 (5.0) |
| Values obtained from replicate 1 and replicate 2 are in the top and bottom respectively. | | | | | | |

(B) duplex-Cm$^{1}$ΨG

| Base pair | **Shear [Å]** | **Stretch [Å]** | **Stagger [Å]** | **Buckle [°]** | **Propeller [°]** | **Opening [°]** |
|---|---|---|---|---|---|---|
| U1-A18 | -0.21 (0.28)<br>-0.21 (0.28) | -0.03 (0.10)<br>-0.03 (0.10) | 0.13 (0.51)<br>0.14 (0.51) | 1.5 (0.51)<br>1.6 (14.4) | -7.7 (10.8)<br>-8.3 (10.8) | 2.6 (4.7)<br>2.5 (4.6) |
| C2-G17 | 0.00 (0.35)<br>-0.01 (0.35) | 0.04 (0.13)<br>0.04 (0.13) | -0.20 (0.43)<br>-0.17 (0.41) | 11.7 (11.1)<br>11.4 (10.9) | -14.5 (8.4)<br>-15.0 (8.2) | 2.0 (3.9)<br>1.9 (3.9) |
| A3-U16 | -0.13 (1.08)<br>0.12 (0.30) | -0.02 (0.22)<br>0.02 (0.15) | -0.11 (0.48)<br>-0.11 (0.48) | 0.8 (9.7)<br>0.5 (9.5) | -14.2 (7.8)<br>-14.5 (7.5) | 3.3 (6.7)<br>3.5 (6.4) |
| G4-C15 | 0.05 (0.33)<br>0.05 (0.33) | 0.01 (0.13)<br>0.01 (0.12) | 0.11 (0.40)<br>0.10 (0.40) | -0.0 (9.2)<br>0.1 (9.1) | -10.7 (7.6)<br>-10.7 (7.5) | 1.5 (3.6)<br>1.5 (3.4) |
| m$^{1}$Ψ5-A14 | -0.22 (0.29)<br>-0.22 (0.30) | 0.02 (0.13)<br>0.02 (0.14) | -0.01 (0.40)<br>-0.02 (0.40) | -0.1 (8.3)<br>0.3 (8.3) | -13.4 (7.1)<br>-13.6 (6.9) | 0.6 (5.6)<br>0.5 (5.6) |
| C6-G13 | 0.06 (0.32)<br>0.06 (0.32) | 0.03 (0.11)<br>0.04 (0.11) | -0.08 (0.39)<br>-0.09 (0.39) | -7.8 (9.0)<br>-7.8 (8.9) | -12.6 (7.6)<br>-12.8 (7.5) | 1.0 (3.4)<br>1.1 (3.3) |
| A7-U12 | 0.05 (0.30)<br>0.05 (0.30) | 0.03 (0.15)<br>0.03 (0.15) | 0.15 (0.47)<br>0.15 (0.46) | -2.5 (9.7)<br>-2.5 (9.7) | -12.3 (7.9)<br>-12.1 (8.0) | 4.7 (6.3)<br>4.6 (6.3) |
| G8-C11 | -0.12 (0.41)<br>-0.12 (0.40) | 0.02 (0.19)<br>0.02 (0.18) | 0.04 (0.44)<br>0.04 (0.44) | -3.5 (11.1)<br>-3.4 (11.0) | -15.8 (9.0)<br>-15.7 (9.1) | 3.0 (5.5)<br>2.9 (5.2) |
| U9-A10 | -0.16 (0.28)<br>-0.16 (0.28) | -0.04 (0.10)<br>-0.04 (0.10) | -0.11 (0.50)<br>-0.12 (0.50) | -0.3 (13.4)<br>-0.2 (13.4) | -10.0 (11.5)<br>-10.0 (11.6) | 1.1 (5.1)<br>1.2 (5.0) |

| Values obtained from replicate 1 and replicate 2 are in the top and bottom respectively. |
|---|

(C) duplex-Um¹ΨA

| Base pair | **Shear [Å]** | **Stretch [Å]** | **Stagger [Å]** | **Buckle [°]** | **Propeller [°]** | **Opening [°]** |
|---|---|---|---|---|---|---|
| U1-A18 | -0.21 (0.28)<br>-0.20 (0.28) | -0.03 (0.11)<br>-0.03 (0.11) | 0.12 (0.51)<br>0.13 (0.51) | 1.0 (15.0)<br>-0.0 (15.3) | -8.8 (10.9)<br>-8.9 (10.9) | 2.5 (4.7)<br>2.5 (4.7) |
| C2-G17 | 0.01 (0.36)<br>0.01 (0.38) | 0.03 (0.13)<br>0.04 (0.14) | -0.16 (0.42)<br>-0.14 (0.42) | 9.8 (11.3)<br>9.2 (11.6) | -15.6 (8.3)<br>-15.0 (8.4) | 2.0 (4.0)<br>2.2 (4..5) |
| A3-U16 | 0.10 (0.31)<br>0.10 (0.31) | 0.03 (0.17)<br>0.03 (0.17) | -0.15 (0.50)<br>-0.14 (0.49) | -2.9 (10.7)<br>-2.7 (11.0) | -16.6 (8.1)<br>-16.2 (8.6) | 4.1 (6.8)<br>4.1 (6.7) |
| U4-A15 | -0.07 (0.42 )<br>-0.10 (0.30) | 0.01 (0.14)<br>0.02 (0.13) | -0.08 (0.49)<br>-0.08 (0.49) | -0.2 (10.6)<br>0.3 (10.6) | -14.9 (8.0)<br>-15.4 (8.0) | 3.3 (6.0)<br>3.4 (6.0) |
| m¹Ψ5-A14 | -0.23 (0.28)<br>-0.23 (0.28) | 0.03 (0.12)<br>0.03 (0.12) | -0.05 (0.43)<br>-0.07 (0.43) | 2.7 (9.8)<br>3.3 (9.8) | -14.6 (7.3)<br>-15.2 (7.4) | 1.2 (5.2)<br>0.9 (5.3) |
| A6-U13 | 0.13 (0.28)<br>0.13 (0.29) | 0.04 (0.13)<br>0.04 (0.13) | -0.08 (0.46)<br>-0.09 (0.46) | -4.7 (9.7)<br>-4.9 (9.7) | -12.8 (7.9)<br>-12.9 (7.8) | 2.3 (5.4)<br>2.5 (5.4) |
| A7-U12 | -0.22 (1.15)<br>-0.12 (0.96) | -0.00 (0.22)<br>0.01 (0.21) | 0.04 (0.47)<br>0.03 (0.47) | 0.7 (10.2)<br>0.9 (9.9) | -13.2 (8.3)<br>-13.0 (8.2) | 3.9 (6.6)<br>3.6 (6.5) |
| G8-C11 | -0.13 (0.38)<br>-0.14 (0.42) | 0.01 (0.15)<br>0.02 (0.19) | 0.01 (0.44)<br>0.02 (0.44) | -2.8 (11.3)<br>-2.4 (11.2) | -14.2 (9.7)<br>-14.8 (9.2) | 2.7 (4.5)<br>3.0 (5.6) |
| U9-A10 | -0.15 (0.28)<br>-0.16 (0.28) | -0.04 (0.10)<br>-0.04 (0.10) | -0.15 (0.51)<br>-0.15 (0.51) | 0.4 (13.9)<br>0.1 (13.5) | -9.3 (11.7)<br>-9.6 (11.6) | 1.4 (5.0)<br>1.4 (5.1) |
| Values obtained from replicate 1 and replicate 2 are in the top and bottom respectively. | | | | | | |

(D) duplex-Am¹ΨU

| Base pair | **Shear [Å]** | **Stretch [Å]** | **Stagger [Å]** | **Buckle [°]** | **Propeller [°]** | **Opening [°]** |
|---|---|---|---|---|---|---|
| U1-A18 | -0.20 (0.28)<br>-0.20 (0.28) | -0.03 (0.11)<br>-0.03 (0.11) | 0.11 (0.51)<br>0.11 (0.52) | -0.2 (15.3)<br>-0.7 (15.6) | -8.7 (10.8)<br>-8.9 (10.9) | 2.7 (4.7)<br>2.7 (4.7) |
| C2-G17 | -0.01 (0.34)<br>-0.01 (0.34) | 0.03 (0.12)<br>0.03 (0.12) | -0.15 (0.41)<br>-0.14 (0.41) | 8.1 (11.0)<br>7.5 (11.3) | -14.5 (8.2)<br>-14.1 (8.5) | 2.0 (3.7)<br>1.9 (3.8) |
| A3-U16 | 0.12 (0.29) | 0.03 (0.14) | -0.18 (0.47) | -3.2 (10.2) | -14.6 (7.7) | 2.4 (5.8) |

| | | | | | | |
|---|---|---|---|---|---|---|
| | 0.12 (0.29) | 0.03 (0.14) | -0.18 (0.46) | -2.9 (10.4) | -14.3 (7.7) | 2.5 (5.7) |
| G4-C15 | 0.12 (0.36)<br>0.11 (0.32) | 0.04 (0.16)<br>0.04 (0.16) | -0.16 (0.52)<br>-0.14 (0.51) | -4.5 (11.1)<br>-3.7 (11.2) | -15.6 (8.4)<br>-15.5 (8.3) | 3.0 (6.6)<br>3.3 (6.5) |
| m1Ψ5-A14 | -0.23 (0.30)<br>-0.22 (0.30) | 0.02 (0.14)<br>0.02 (0.14) | -0.09 (0.46)<br>-0.12 (0.46) | -0.7 (10.5)<br>-0.1 (10.7) | -17.5 (8.0)<br>-18.0 (7.8) | 1.8 (5.9)<br>1.8 (5.9) |
| C6-G13 | -0.10 (0.28)<br>-0.09 (0.31) | 0.04 (0.13)<br>0.04 (0.13) | 0.03 (0.45)<br>0.02 (0.43) | 1.5 (10.2)<br>1.1 (9.7) | -12.6 (7.7)<br>-13.0 (7.6) | 3.2 (5.6)<br>3.2 (5.5) |
| A7-U12 | -0.24 (1.25)<br>0.07 (0.30) | -0.03 (0.23)<br>0.02 (0.13) | 0.08 (0.53)<br>0.09 (0.44) | -0.3 (9.4)<br>-0.5 (9.3) | -12.1 (9.2)<br>-12.8 (8.1) | 3.1 (6.1)<br>3.2 (5.7) |
| G8-C11 | -0.12 (0.38)<br>-0.12 (0.38) | 0.01 (0.16)<br>0.01 (0.16) | 0.05 (0.44)<br>0.06 (0.44) | -1.7 (11.4)<br>-1.9 (11.3) | -14.6 (9.3)<br>-15.0 (9.1) | 2.6 (4.9)<br>2.6 (4.7) |
| U9-A10 | -0.15 (0.28)<br>-0.15 (0.28) | -0.04 (0.10)<br>-0.04 (0.10) | -0.15 (0.52)<br>-0.14 (0.51) | 1.3 (13.9)<br>1.3 (13.9) | -9.7 (11.7)<br>-10.1 (11.7) | 1.4 (5.1)<br>1.4 (5.1) |
| Values obtained from replicate 1 and replicate 2 are in the top and bottom respectively. | | | | | | |

**Table S5.** Average values and standard deviations (in parentheses) of local base-pair parameters for duplexes (Gm$^1$ΨC context) containing (A) m$^1$Ψ-G, (B) m$^1$Ψ-U, (C) m$^1$Ψ-C mismatches.

(A) m$^1$Ψ-G

| Base pair | **Shear [Å]** | **Stretch [Å]** | **Stagger [Å]** | **Buckle [°]** | **Propeller [°]** | **Opening [°]** |
|---|---|---|---|---|---|---|
| U1-A18 | -0.22 (0.28)<br>-0.20 (0.28) | -0.02 (0.10)<br>-0.03 (0.10) | 0.04 (0.50)<br>0.13 (0.51) | -1.5 (14.7)<br>-1.2 (15.1) | -3.5 (11.7)<br>-8.8 (10.8) | 2.6 (4.7)<br>2.5 (4.7) |
| C2-G17 | 0.16 (0.39)<br>-0.02 (0.33) | 0.01 (0.13)<br>0.04 (0.12) | -0.34 (0.47)<br>-0.12 (0.41) | 11.8 (11.9)<br>8.5 (10.7) | -9.6 (8.8)<br>-14.3 (8.1) | 2.0 (3.8)<br>1.8 (3.5) |
| A3-U16 | -2.98 (2.25)<br>0.09 (0.31) | -0.39 (0.38)<br>0.03 (0.17) | -0.03 (0.57)<br>0.04 (0.44) | 4.9 (9.7)<br>0.6 (8.8) | -13.7 (8.7)<br>-14.2 (7.4) | 2.6 (7.8)<br>3.5 (6.7) |
| G4-C15 | -0.05 (0.33)<br>-0.01 (0.34) | 0.02 (0.13)<br>0.03 (0.19) | 0.03 (0.40)<br>0.07 (0.39) | -3.2 (8.6)<br>-4.7 (8.7) | -14.8 (7.8)<br>-15.4 (7.2) | 2.0 (4.1)<br>1.6 (4.8) |
| m1Ψ5-G14 | 2.30 (0.44)<br>2.24 (0.38) | 0.07 (0.67)<br>-0.13 (0.48) | 0.28 (0.53)<br>0.25 (0.47) | -4.4 (8.5)<br>-3.3 (8.2) | -11.9 (7.0)<br>-11.8 (6.8) | 9.8 (16.5)<br>4.2 (11.8) |
| C6-G13 | 0.04 (0.44)<br>-0.02 (0.34) | 0.02 (0.16)<br>0.01 (0.11) | 0.02 (0.39)<br>0.00 (0.36) | 0.3 (9.0)<br>2.1 (8.4) | -10.6 (7.7)<br>-12.0 (7.3) | 1.8 (5.4)<br>1.2 (3.5) |

| | | | | | | |
|---|---|---|---|---|---|---|
| A7-U12 | 0.01 (0.81)<br>0.13 (0.31) | 0.03 (0.19)<br>0.03 (0.14) | 0.05 (0.44)<br>0.04 (0.44) | 2.0 (10.2)<br>0.9 (9.1) | -9.2 (8.8)<br>-11.3 (7.8) | 2.5 (6.6)<br>1.8 (6.1) |
| G8-C11 | -0.13 (0.50)<br>-0.11 (0.39) | 0.05 (0.22)<br>0.02 (0.16) | 0.06 (0.47)<br>0.05 (0.44) | 4.4 (12.7)<br>-0.5 (11.4) | -11.4 (10.5)<br>-14.4 (9.2) | 3.4 (6.8)<br>2.8 (4.8) |
| U9-A10 | -0.17 (0.28)<br>-0.15 (0.28) | -0.04 (0.10)<br>-0.04 (0.11) | -0.16 (0.52)<br>-0.19 (0.52) | 2.0 (13.9)<br>2.3 (14.0) | -10.7 (12.1)<br>-9.9 (11.8) | 1.2 (5.1)<br>1.6 (5.1) |
| Values obtained from replicate 1 and replicate 2 are in the top and bottom respectively. | | | | | | |

(B) m$^1\Psi$-U

| Base pair | **Shear [Å]** | **Stretch [Å]** | **Stagger [Å]** | **Buckle [°]** | **Propeller [°]** | **Opening [°]** |
|---|---|---|---|---|---|---|
| U1-A18 | -0.19 (0.28)<br>-0.19 (0.28) | -0.02 (0.11)<br>-0.02 (0.11) | 0.08 (0.52)<br>0.10 (0.52) | -2.5 (16.8)<br>-3.8 (16.4) | -8.1 (11.6)<br>-9.0 (11.2) | 2.9 (4.7)<br>2.8 (4.7) |
| C2-G17 | -0.03 (0.33)<br>-0.01 (0.35) | 0.04 (0.12)<br>0.04 (0.12) | -0.10 (0.41)<br>-0.13 (0.43) | 6.7 (11.6)<br>6.8 (11.8) | -13.2 (8.5)<br>-12.3 (8.8) | 1.9 (3.5)<br>1.9 (3.6) |
| A3-U16 | 0.12 (0.31)<br>-0.36 (1.47) | 0.03 (0.16)<br>-0.03 (0.24) | -0.02 (0.44)<br>-0.04 (0.47) | 2.1 (9.2)<br>2.5 (9.6) | -11.2 (7.8)<br>-11.4 (8.1) | 2.0 (6.4)<br>1.7 (6.6) |
| G4-C15 | -0.07 (0.36)<br>-0.07 (0.35) | 0.00 (0.14)<br>0.00 (0.13) | -0.17 (0.44)<br>-0.19 (0.44) | -3.7 (11.0)<br>-4.4 (10.8) | -15.1 (8.0)<br>-15.6 (8.0) | 3.0 (4.2)<br>3.0 (4.0) |
| m$^1\Psi$5-U14 | 0.26 (2.65)<br>-0.08 (2.69) | -1.37 (0.62)<br>-1.32 (0.70) | 0.01 (0.61)<br>0.03 (0.62) | -2.0 (13.1)<br>-1.6 (13.5) | -16.2 (8.2)<br>-15.8 (8.1) | -1.3 (14.7)<br>-2.0 (15.5) |
| C6-G13 | 0.09 (0.46)<br>0.09 (0.50) | 0.02 (0.20)<br>0.03 (0.25) | -0.15 (0.45)<br>-0.13 (0.45) | 7.3 (10.5)<br>6.3 (10.4) | -17.4 (7.9)<br>-17.3 (7.9) | 3.6 (5.6)<br>3.7 (6.4) |
| A7-U12 | 0.03 (0.75)<br>0.14 (0.31) | 0.02 (0.20)<br>0.04 (0.17) | -0.04 (0.45)<br>-0.04 (0.45) | -1.9 (9.7)<br>-3.2 (9.6) | -12.1 (8.2)<br>-12.4 (8.1) | 2.2 (6.7)<br>2.2 (6.6) |
| G8-C11 | -0.07 (0.37)<br>-0.07 (0.37) | 0.03 (0.15)<br>0.03 (0.15) | 0.03 (0.44)<br>0.03 (0.44) | -2.6 (11.5)<br>-3.3 (11.7) | -13.5 (9.4)<br>-13.7 (9.7) | 2.6 (4.5)<br>2.6 (4.5) |
| U9-A10 | -0.16 (0.28)<br>-0.16 (0.28) | -0.04 (0.11)<br>-0.04 (0.11) | -0.17 (0.52)<br>-0.19 (0.52) | 2.3 (14.4)<br>0.8 (15.0) | -8.8 (12.0)<br>-7.4 (12.2) | 1.6 (5.1)<br>1.8 (5.0) |
| Values obtained from replicate 1 and replicate 2 are in the top and bottom respectively. | | | | | | |

(C) m$^1\Psi$-C

| Base pair | **Shear [Å]** | **Stretch [Å]** | **Stagger [Å]** | **Buckle [°]** | **Propeller [°]** | **Opening [°]** |
|---|---|---|---|---|---|---|
| U1-A18 | -0.19 (0.28)<br>-0.21 (0.28) | -0.03 (0.11)<br>-0.02 (0.11) | 0.12 (0.51)<br>0.08 (0.51) | -1.5 (15.6)<br>-2.0 (16.5) | -8.9 (10.8)<br>-6.7 (11.9) | 2.6 (4.7)<br>2.7 (4.7) |
| C2-G17 | -0.03 (0.33)<br>0.06 (0.37) | 0.04 (0.12)<br>0.03 (0.13) | -0.13 (0.41)<br>-0.20 (0.45) | 8.2 (11.2)<br>9.0 (12.7) | -14.0 (8.2)<br>-12.3 (8.9) | 2.0 (3.6)<br>2.0 (3.8) |
| A3-U16 | 0.10 (0.31)<br>-1.10 (2.08) | 0.03 (0.17)<br>-0.13 (0.34) | 0.02 (0.45)<br>-0.00 (0.50) | 1.2 (9.2)<br>3.0 (9.7) | -12.9 (7.9)<br>-12.4 (8.5) | 3.0 (6.8)<br>2.7 (7.2) |
| G4-C15 | -0.01 (0.36)<br>-0.02 (0.34) | 0.02 (0.13)<br>0.02 (0.12) | -0.01 (0.42)<br>0.01 (0.42) | -3.3 (10.6)<br>-1.4 (11.2) | -13.9 (8.4)<br>-13.4 (8.5) | 1.9 (4.1)<br>2.0 (3.9) |
| m[1]Ψ5-C14 | 1.71 (2.69)<br>1.58 (2.70) | -0.73 (0.83)<br>-0.75 (0.86) | -0.44 (0.80)<br>-0.45 (0.80) | -7.1 (13.1)<br>-6.5 (13.4) | -16.4 (9.0)<br>-16.4 (9.3) | -16.8 (19.0)<br>-17.0 (21.5) |
| C6-G13 | 0.07 (0.51)<br>0.13 (0.68) | 0.03 (0.19)<br>0.07 (0.29) | 0.07 (0.44)<br>0.09 (0.46) | 0.7 (9.7)<br>-0.1 (10.3) | -12.6 (7.6)<br>-12.6 (7.5) | 2.2 (6.8)<br>3.3 (9.9) |
| A7-U12 | -0.08 (1.01)<br>0.12 (0.31) | 0.01 (0.22)<br>0.03 (0.16) | 0.05 (0.46)<br>0.04 (0.46) | 0.7 (9.5)<br>-0.1 (9.7) | -11.6 (8.2)<br>-11.6 (8.0) | 2.5 (6.8)<br>2.4 (6.6) |
| G8-C11 | -0.11 (0.39)<br>-0.11 (0.40) | 0.02 (0.18)<br>0.02 (0.19) | 0.05 (0.44)<br>0.05 (0.44) | -0.6 (11.6)<br>-1.0 (11.6) | -14.4 (9.4)<br>-14.4 (9.4) | 2.9 (5.1)<br>2.9 (5.3) |
| U9-A10 | -0.15 (0.28)<br>-0.15 (0.28) | -0.04 (0.11)<br>-0.04 (0.11) | -0.18 (0.52)<br>-0.17 (0.52) | 2.1 (14.1)<br>2.4 (14.3) | -9.9 (11.8)<br>-9.7 (11.8) | 1.6 (5.1)<br>1.6 (5.1) |
| Values obtained from replicate 1 and replicate 2 are in the top and bottom respectively. | | | | | | |

**Table S6.** Average values and standard deviations (in parentheses) of local base-pair step parameters for (A) duplex-Gm[1]ΨC, (B) duplex-Cm[1]ΨG, (C) duplex-Um[1]ΨA and (D) duplex-Am[1]ΨU containing m[1]Ψ-A pair.

(A) duplex-Gm[1]ΨC

| Base-pair step | Shift [Å] | Slide [Å] | Rise [Å] | Tilt [°] | Roll [°] | Twist [°] |
|---|---|---|---|---|---|---|
| U1-A18/C2-G17 | -0.17 (0.62)<br>-0.18 (0.61) | -1.35 (0.53)<br>-1.34 (0.54) | 3.11 (0.37)<br>3.13 (0.38) | 1.1 (5.2)<br>0.9 (5.3) | 6.6 (7.1)<br>6.2 (7.2) | 30.6 (4.1)<br>30.6 (4.2) |
| C2-G17/A3-U16 | 0.02 (0.76)<br>-0.02 (0.72) | -1.33 (0.40)<br>-1.30 (0.40) | 3.48 (0.39)<br>3.46 (0.39) | -0.8 (4.3)<br>-0.9 (4.3) | 14.0 (6.9)<br>14.4 (6.9) | 32.2 (3.5)<br>32.2 (3.4) |

| | | | | | | |
|---|---|---|---|---|---|---|
| A3-U16/G4-C15 | -0.01 (0.85)<br>-0.04 (0.78) | -1.76 (0.39)<br>-1.74 (0.40) | 3.34 (0.32)<br>3.33 (0.32) | -1.3 (4.5)<br>-1.4 (4.5) | 9.4 (6.1)<br>9.6 (6.1) | 29.1 (3.9)<br>29.3 (3.4) |
| G4-C15/m[1]Ψ5-A14 | -0.06 (0.72)<br>-0.07 (0.73) | -1.98 (0.51)<br>-1.97 (0.51) | 3.31 (0.25)<br>3.29 (0.25) | 0.8 (4.0)<br>0.7 (4.0) | 5.0 (4.8)<br>4.8 (4.7) | 29.5 (5.1)<br>28.6 (3.9) |
| m[1]Ψ5-A14/C6-G13 | 0.20 (0.71)<br>0.19 (0.69) | -1.78 (0.42)<br>-1.73 (0.42) | 3.24 (0.34)<br>3.26 (0.33) | 0.3 (4.4)<br>0.1 (4.3) | 7.3 (5.6)<br>7.4 (5.6) | 28.2 (4.8)<br>29.3 (3.5) |
| C6-G13/A7-U12 | -0.05 (0.72)<br>-0.02 (0.73) | -1.42 (0.39)<br>-1.41 (0.39) | 3.31 (0.36)<br>3.36 (0.37) | 0.5 (4.2)<br>0.7 (4.3) | 10.3 (6.9)<br>11.2 (7.0) | 31.7 (3.4)<br>31.9 (3.4) |
| A7-U12/G8-C11 | -0.02 (0.80)<br>-0.01 (0.82) | -1.67 (0.43)<br>-1.61 (0.44) | 3.32 (0.34)<br>3.32 (0.35) | -0.7 (4.6)<br>-0.6 (4.6) | 9.5 (6.4)<br>9.4 (6.6) | 28.8 (3.8)<br>29.9 (4.2) |
| G8-C11/U9-A10 | 0.15 (0.66)<br>0.04 (0.74) | -1.78 (0.62)<br>-1.49 (0.74) | 3.28 (0.30)<br>3.23 (0.30) | 1.3 (4.6)<br>2.5 (4.9) | 6.1 (6.3)<br>5.1 (6.4) | 30.2 (4.6)<br>28.4 (6.0) |
| Values obtained from replicate 1 and replicate 2 are in the top and bottom respectively. | | | | | | |

(B) duplex-Cm[1]ΨG

| Base-pair step | Shift [Å] | Slide [Å] | Rise [Å] | Tilt [°] | Roll [°] | Twist [°] |
|---|---|---|---|---|---|---|
| U1-A18/C2-G17 | -0.19 (0.59)<br>-0.18 (0.58) | -1.37 (0.49)<br>-1.37 (0.49) | 3.09 (0.37)<br>3.10 (0.37) | 1.5 (5.2)<br>1.3 (5.2) | 6.1 (7.0)<br>6.0 (7.0) | 30.6 (4.1)<br>30.8 (4.1) |
| C2-G17/A3-U16 | 0.04 (0.73)<br>0.06 (0.71) | -1.40 (0.43)<br>-1.36 (0.40) | 3.58 (0.43)<br>3.58 (0.41) | -1.2 (4.8)<br>-0.7 (4.4) | 15.0 (7.1)<br>15.1 (7.0) | 31.5 (5.0)<br>32.5 (3.4) |
| A3-U16/G4-C15 | -0.05 (0.74)<br>-0.06 (0.74) | -1.86 (0.50)<br>-1.87 (0.50) | 3.34 (0.25)<br>3.33 (0.25) | -1.9 (4.2)<br>-1.7 (4.1) | 7.2 (5.4)<br>7.2 (5.3) | 29.5 (5.2)<br>28.6 3.6 |
| G4-C15/m[1]Ψ5-A14 | -0.04 (0.67)<br>-0.04 (0.65) | -1.80 (0.37)<br>-1.78 (0.36) | 3.30 (0.31)<br>3.29 (0.31) | 2.3 (3.9)<br>2.3 (3.9) | 6.9 (5.5)<br>6.8 (5.4) | 29.1 (3.5)<br>29.1 (3.3) |
| m[1]Ψ5-A14/C6-G13 | 0.15 (0.62)<br>0.14 (0.62) | -1.43 (0.38)<br>-1.41 (0.37) | 3.49 (0.35)<br>3.49 (0.35) | 0.0 (4.1)<br>-0.1 (4.0) | 12.4 (6.7)<br>12.8 (6.6) | 32.6 (3.3)<br>32.9 (3.1) |
| C6-G13/A7-U12 | 0.20 (0.69)<br>0.21 (0.69) | -1.61 (0.40)<br>-1.61 (0.40) | 3.16 (0.33)<br>3.16 (0.33) | -1.4 (4.4)<br>-1.4 (4.3) | 6.7 (6.1)<br>6.8 (6.1) | 29.1 (3.3)<br>29.0 (3.2) |
| A7-U12/G8-C11 | -0.22 (0.72)<br>-0.22 (0.72) | -1.70 (0.42)<br>-1.71 (0.42) | 3.30 (0.34)<br>3.30 (0.34) | 0.1 (4.7)<br>0.1 (4.7) | 8.2 (6.0)<br>8.2 (6.0) | 29.2 (3.8)<br>29.1 (3.9) |
| G8-C11/U9-A10 | 0.15 (0.66)<br>0.15 (0.65) | -1.73 (0.60)<br>-1.74 (0.60) | 3.25 (0.28)<br>3.26 (0.28) | 1.4 (4.5)<br>1.5 (4.5) | 6.1 (6.2)<br>6.1 (6.2) | 30.3 (4.4)<br>30.3 (4.4) |
| Values obtained from replicate 1 and replicate 2 are in the top and bottom respectively. | | | | | | |

(C) duplex-Um$^{1}\Psi$A

| Base-pair step | Shift [Å] | Slide [Å] | Rise [Å] | Tilt [°] | Roll [°] | Twist [°] |
|---|---|---|---|---|---|---|
| U1-A18/C2-G17 | -0.20 (0.60)<br>-0.16 (0.61) | -1.35 (0.51)<br>-1.34 (0.52) | 3.12 (0.37)<br>3.11 (0.37) | 1.2 (5.2)<br>1.2 (5.2) | 6.5 (7.0)<br>6.2 (7.0) | 31.0 (4.1)<br>30.8 (4.3) |
| C2-G17/A3-U16 | 0.12 (0.72)<br>0.11 (0.74) | -1.31 (0.40)<br>-1.29 (0.42) | 3.61 (0.43)<br>3.60 (0.44) | 0.1 (4.6)<br>0.3 (4.9) | 16.4 (7.1)<br>15.9 (7.4) | 32.4 (3.5)<br>32.4 (3.7) |
| A3-U16/U4-A15 | -0.01 (0.82)<br>-0.00 (0.81) | -1.58 (0.51)<br>-1.55 (0.51) | 3.23 (0.28)<br>3.23 (0.29) | -0.4 (4.6)<br>-0.3 (4.7) | 8.4 (6.0)<br>8.7 (6.1) | 28.3 (3.5)<br>28.2 (3.5) |
| U4-A15/m$^{1}\Psi$5-A14 | -0.01 (0.64)<br>0.03 (0.69) | -1.54 (0.43)<br>-1.53 (0.44) | 3.30 (0.35)<br>3.29 (0.35) | 0.8 (4.4)<br>0.9 (4.4) | 7.5 (5.9)<br>7.7 (5.9) | 29.0 (3.5)<br>28.6 (4.3) |
| m$^{1}\Psi$5-A14/A6-U13 | 0.17 (0.53)<br>0.16 (0.53) | -1.38 (0.35)<br>-1.37 (0.34) | 3.47 (0.39)<br>3.48 (0.39) | -0.3 (4.6)<br>-0.5 (4.6) | 15.1 (7.1)<br>15.5 (7.1) | 31.7 (3.3)<br>32.2 (3.2) |
| A6-U13/A7-U12 | -0.03 (0.71)<br>-0.05 (0.70) | -1.57 (0.44)<br>-1.56 (0.43) | 3.15 (0.35)<br>3.15 (0.34) | -1.6 (4.9)<br>-1.5 (4.9) | 7.3 (6.4)<br>7.4 (6.3) | 27.8 (5.1)<br>27.9 (4.6) |
| A7-U12/G8-C11 | -0.14 (0.72)<br>-0.12 (0.74) | -1.68 (0.44)<br>-1.70 (0.43) | 3.37 (0.35)<br>3.36 (0.35) | -0.9 (5.1)<br>-1.1 (4.8) | 8.7 (6.4)<br>9.4 (6.2) | 30.5 (5.5)<br>29.9 (5.1) |
| G8-C11/U9-A10 | 0.15 (0.64)<br>0.16 (0.65) | -1.77 (0.64)<br>-1.80 (0.61) | 3.26 (0.29)<br>3.27 (0.28) | 1.5 (4.5)<br>1.6 (4.5) | 6.1 (6.2)<br>6.0 (6.2) | 30.1 (4.7)<br>30.4 (4.5) |
| Values obtained from replicate 1 and replicate 2 are in the top and bottom respectively. | | | | | | |

(D) duplex-Am$^{1}\Psi$U

| Base-pair step | Shift [Å] | Slide [Å] | Rise [Å] | Tilt [°] | Roll [°] | Twist [°] |
|---|---|---|---|---|---|---|
| U1-A18/C2-G17 | -0.18 (0.59)<br>-0.19 (0.61) | -1.37 (0.52)<br>-1.37 (0.53) | 3.13 (0.37)<br>3.14 (0.38) | 1.2 (5.2)<br>1.1 (5.2) | 6.6 (7.0)<br>6.7 (7.1) | 30.9 (4.1)<br>30.7 (4.1) |
| C2-G17/A3-U16 | -0.02 (0.67)<br>-0.02 (0.67) | -1.29 (0.38)<br>-1.28 (0.40) | 3.60 (0.42)<br>3.58 (0.42) | 0.2 (4.4)<br>0.2 (4.4) | 15.8 (7.1 )<br>15.8 (7.1) | 32.6 (3.3)<br>32.4 (3.4) |
| A3-U16/A4-U15 | -0.01 (0.72)<br>-0.02 (0.72) | -1.50 (0.43)<br>-1.50 (0.44) | 3.35 (0.37)<br>3.33 (0.37) | -0.6 (5.0)<br>-0.8 (5.0) | 10.5 (6.5)<br>10.4 (6.6) | 29.3 (3.4)<br>29.2 (3.4) |
| A4-U15/m$^{1}\Psi$5-A14 | -0.10 (0.83)<br>-0.17 (0.83) | -1.70 (0.52)<br>-1.66 (0.50) | 3.23 (0.28)<br>3.23 (0.28) | 0.1 (4.4)<br>0.2 (4.4) | 6.1 (5.4)<br>6.3 (5.4) | 28.3 (4.0)<br>28.7 (3.7) |

| | | | | | | |
|---|---|---|---|---|---|---|
| m[1]Ψ5-A14/U6-A13 | 0.35 (0.61)<br>0.33 (0.61) | -1.54 (0.44)<br>-1.51 (0.42) | 3.27 (0.33)<br>3.29 (0.33) | -0.4 (4.7)<br>-0.5 (4.7) | 8.3 (6.0)<br>9.0 (5.9) | 29.0 (3.6)<br>29.3 (3.5) |
| U6-A13/A7-U12 | -0.03 (0.66)<br>-0.01 (0.56) | -1.42 (0.39)<br>-1.38 (0.35) | 3.28 (0.40)<br>3.29 (0.36) | -0.7 (5.1)<br>-0.5 (4.5) | 12.6 (7.3)<br>13.3 (7.0) | 30.2 (5.0)<br>31.1 (3.1) |
| A7-U12/G8-C11 | -0.12 (0.71)<br>-0.12 (0.70) | -1.64 (0.44)<br>-1.66 (0.43) | 3.32 (0.35)<br>3.31 (0.34) | -0.9 (4.8)<br>-0.6 (4.6) | 9.1 (6.3)<br>9.2 (6.2) | 30.1 (5.5)<br>29.1 (3.7) |
| G8-C11/U9-A10 | 0.18 (0.64)<br>0.19 (0.64) | -1.76 (0.61)<br>-1.73 (0.61) | 3.27 (0.28)<br>3.26 (0.28) | 1.6 (4.5)<br>1.6 (4.5) | 6.3 (6.3)<br>6.3 (6.2) | 30.2 (4.4)<br>30.3 (4.5) |
| Values obtained from replicate 1 and replicate 2 are in the top and bottom respectively. | | | | | | |

**Table S7.** Average values and standard deviations (in parentheses) of local base-pair step parameters for duplexes (Gm[1]ΨC context) containing (A) m[1]Ψ-G, (B) m[1]Ψ-U, (C) m[1]Ψ-C mismatches respectively.

(A) m[1]Ψ-G

| Base-pair step | Shift [Å] | Slide [Å] | Rise [Å] | Tilt [°] | Roll [°] | Twist [°] |
|---|---|---|---|---|---|---|
| U1-A18/C2-G17 | -0.19 (0.59)<br>-0.15 (0.60) | -1.30 (0.51)<br>-1.35 (0.52) | 3.05 (0.35)<br>3.10 (0.36) | 2.1 (5.1)<br>1.1 (5.1) | 7.2 (6.9)<br>6.3 (7.0) | 28.7 (4.3)<br>30.6 4.1 |
| C2-G17/A3-U16 | 0.02 (0.71)<br>0.12 (0.73) | -1.71 (0.53)<br>-1.35 (0.39) | 3.33 (0.54)<br>3.48 (0.38) | -5.5 (5.9)<br>-1.0 (4.2) | 14.1 (7.0)<br>13.6 (6.6) | 21.7 (8.6)<br>32.2 3.3 |
| A3-U16/G4-C15 | -0.12 (0.63)<br>-0.16 (0.78) | -1.63 (0.39)<br>-1.74 (0.39) | 3.56 (0.34)<br>3.38 (0.31) | -4.3 (5.0)<br>-0.9 (4.4) | 9.3 (5.5)<br>9.8 (5.9) | 40.6 (8.2)<br>30.3 (3.4) |
| G4-C15/m[1]Ψ5-G14 | 0.54 (0.90)<br>0.32 (0.72) | -1.76 (0.44)<br>-1.69 (0.43) | 3.36 (0.24)<br>3.31 (0.23) | 1.2 (4.0)<br>1.4 (4.0) | 4.7 (4.0)<br>5.1 (4.2) | 38.3 (3.8)<br>37.7 (3.6) |
| m[1]Ψ5-G14/C6-G13 | -0.65 (1.15)<br>-0.28 (0.89) | -2.06 (0.43)<br>-1.96 (0.40) | 3.00 (0.35)<br>2.97 (0.32) | 4.2 (4.3)<br>4.3 (4.1) | 7.6 (5.4)<br>7.8 (5.3) | 20.3 (4.1)<br>21.3 (3.5) |
| C6-G13/A7-U12 | -0.04 (0.71)<br>-0.04 (0.67) | -1.30 (0.44)<br>-1.33 (0.38) | 3.27 (0.38)<br>3.33 (0.34) | -0.3 (4.6)<br>-0.3 (4.1) | 11.1 (7.0)<br>11.6 (6.7) | 29.8 (4.6)<br>30.9 (3.3) |
| A7-U12/G8-C11 | 0.14 (0.86)<br>0.03 (0.78) | -1.61 (0.49)<br>-1.65 (0.42) | 3.25 (0.38)<br>3.32 (0.34) | -1.2 (5.1)<br>-0.9 (4.5) | 8.6 (6.9)<br>9.6 (6.5) | 27.9 (5.4)<br>28.9 (3.6) |
| G8-C11/U9-A10 | -0.05 (0.75)<br>0.18 (0.66) | -1.87 (0.68)<br>-1.77 (0.62) | 3.39 (0.35)<br>3.27 (0.29) | 0.2 (5.2)<br>1.6 (4.6) | 4.9 (6.7)<br>6.5 (6.3) | 31.5 (5.2)<br>30.2 (4.6) |

| Values obtained from replicate 1 and replicate 2 are in the top and bottom respectively. |
|---|

(B) m$^1$Ψ-U

| Base-pair step | Shift [Å] | Slide [Å] | Rise [Å] | Tilt [°] | Roll [°] | Twist [°] |
|---|---|---|---|---|---|---|
| U1-A18/C2-G17 | -0.19 (0.61)<br>-0.14 (0.59) | -1.29 (0.58)<br>-1.29 (0.58) | 3.14 (0.37)<br>3.10 (0.37) | 0.4 (5.4)<br>1.0 (5.2) | 6.2 (7.5)<br>6.8 (6.9) | 30.5 (4.2)<br>30.0 (4.3) |
| C2-G17/A3-U16 | -0.04 (0.72)<br>-0.08 (0.71) | -1.26 (0.41)<br>-1.31 (0.45) | 3.43 (0.40)<br>3.41 (0.43) | -0.8 (4.3)<br>-1.3 (5.0) | 13.6 (7.0)<br>13.9 (7.0) | 32.1 (3.4)<br>30.3 (6.1) |
| A3-U16/G4-C15 | 0.08 (0.81)<br>0.07 (0.76) | -1.69 (0.41)<br>-1.65 (0.40) | 3.46 (0.35)<br>3.50 (0.35) | 0.0 (4.6)<br>-0.4 (4.9) | 11.2 (6.2)<br>11.5 (6.2) | 28.0 (4.0)<br>30.0 (6.1) |
| G4-C15/m$^1$Ψ5-U14 | -0.07 (1.12)<br>-0.02 (1.19) | -1.77 (0.61)<br>-1.83 (0.62) | 3.30 (0.53)<br>3.25 (0.54) | -1.3 (6.3)<br>-1.5 (6.3) | 7.1 (5.6)<br>6.9 (5.7) | 31.6 (12.5)<br>30.0 (13.3) |
| m$^1$Ψ5-U14/C6-G13 | 0.25 (1.03)<br>0.24 (1.06) | -1.47 (0.63)<br>-1.55 (0.58) | 3.05 (0.46)<br>3.08 (0.48) | 3.1 (6.0)<br>2.5 (6.1) | 6.2 (5.2)<br>5.8 (5.2) | 29.6 (11.7)<br>31.2 (12.1) |
| C6-G13/A7-U12 | -0.21 (0.72)<br>-0.21 (0.75) | -1.36 (0.41)<br>-1.36 (0.41) | 3.52 (0.40)<br>3.53 (0.39) | -0.9 (4.5)<br>-0.4 (4.4) | 14.5 (7.1)<br>14.5 (6.9) | 30.8 (4.4)<br>31.0 (3.9) |
| A7-U12/G8-C11 | 0.04 (0.81)<br>0.08 (0.81) | -1.61 (0.42)<br>-1.57 (0.43) | 3.30 (0.35)<br>3.29 (0.35) | -0.9 (4.6)<br>-0.6 (4.6) | 9.7 (6.6)<br>9.9 (6.6) | 29.0 (4.4)<br>28.8 (3.8) |
| G8-C11/U9-A10 | 0.17 (0.67)<br>0.20 (0.71) | -1.69 (0.65)<br>-1.57 (0.70) | 3.24 (0.29)<br>3.27 (0.30) | 1.6 (4.6)<br>2.3 (5.0) | 6.4 (6.3)<br>5.6 (6.5) | 29.5 (4.7)<br>29.6 (4.8) |
| Values obtained from replicate 1 and replicate 2 are in the top and bottom respectively. | | | | | | |

(C) m$^1$Ψ-C

| Base-pair step | Shift [Å] | Slide [Å] | Rise [Å] | Tilt [°] | Roll [°] | Twist [°] |
|---|---|---|---|---|---|---|
| U1-A18/C2-G17 | -0.14 (0.59)<br>-0.17 (0.60) | -1.33 (0.53)<br>-1.27 (0.58) | 3.11 (0.37)<br>3.09 (0.37) | 1.1 (5.1)<br>1.2 (5.3) | 6.5 (7.0)<br>6.5 (7.1) | 30.5 (4.1)<br>29.9 (4.3) |
| C2-G17/A3-U16 | 0.06 (0.73)<br>0.01 (0.73) | -1.33 (0.40)<br>-1.43 (0.52) | 3.47 (0.39)<br>3.40 (0.47) | -1.0 (4.3)<br>-2.7 (5.5) | 13.7 (6.8)<br>13.9 (6.9) | 32.2 (3.4)<br>28.1 (7.9) |
| A3-U16/G4-C15 | -0.05 (0.80) | -1.76 (0.40) | 3.39 (0.34) | -0.6 (4.6) | 10.1 (6.1) | 29.7 (3.7) |

| | | | | | | |
|---|---|---|---|---|---|---|
| | -0.04 (0.74) | -1.68 (0.44) | 3.42 (0.36) | -2.1 (5.1) | 9.7 (5.9) | 33.6 (8.0) |
| G4-C15/m$^1$Ψ5-C14 | -1.31 (1.46)<br>-1.42 (1.63) | -1.66 (0.47)<br>-1.70 (0.48) | 3.55 (0.42)<br>3.58 (0.43) | 3.3 (5.2)<br>2.9 (5.6) | 8.3 (5.5)<br>8.7 (6.6) | 37.8 (11.0)<br>37.8 (10.8) |
| m$^1$Ψ5-C14/C6-G13 | 1.38 (1.34)<br>1.42 (1.44) | -1.84 (0.50)<br>-1.85 (0.48) | 3.07 (0.52)<br>3.10 (0.53) | 0.8 (6.6)<br>0.5 (6.5) | 6.5 (5.5)<br>6.5 (5.6) | 22.8 (11.3)<br>24.2 (11.8) |
| C6-G13/A7-U12 | -0.10 (0.79)<br>-0.19 (0.88) | -1.36 (0.46)<br>-1.36 (0.47) | 3.29 (0.37)<br>3.29 (0.37) | -0.3 (4.8)<br>0.3 (4.9) | 11.4 (6.9)<br>11.6 (6.9) | 29.9 (5.4)<br>30.1 (4.6) |
| A7-U12/G8-C11 | 0.02 (0.83)<br>0.01 (0.81) | -1.64 (0.44)<br>-1.63 (0.43) | 3.31 (0.34)<br>3.30 (0.34) | -1.0 (4.7)<br>-0.9 (4.6) | 9.6 (6.4)<br>9.9 (6.4) | 29.2 (5.4)<br>28.6 (3.9) |
| G8-C11/U9-A10 | 0.17 (0.65)<br>0.17 (0.67) | -1.76 (0.63)<br>-1.73 (0.63) | 3.27 (0.29)<br>3.26 (0.29) | 1.6 (4.6)<br>1.6 (4.6) | 6.5 (6.3)<br>6.6 (6.3) | 30.3 (4.6)<br>30.1 (4.6) |
| Values obtained from replicate 1 and replicate 2 are in the top and bottom respectively. | | | | | | |

**Table S8.** Average values and standard deviations (in parentheses) of local base-pair parameters for duplexes (GUC context) containing (A) U-G, (B) U-U, (C) U-C respectively and duplexes (GΨC context) containing (D) Ψ-G, (E) Ψ-U, (F) Ψ-C mismatches respectively.

(A) U-G

| Base pair | **Shear [Å]** | **Stretch [Å]** | **Stagger [Å]** | **Buckle [°]** | **Propeller [°]** | **Opening [°]** |
|---|---|---|---|---|---|---|
| U1-A18 | -0.22 (0.28) | -0.03 (0.10) | 0.12 (0.51) | 0.1 (14.1) | -7.3 (11.1) | 2.5 (4.7) |
| C2-G17 | 0.04 (0.36) | 0.02 (0.12) | -0.19 (0.44) | 10.7 (10.8) | -13.6 (8.4) | 1.7 (3.5) |
| A3-U16 | -0.99 (1.96) | -0.13 (0.31) | 0.07 (0.47) | 2.6 (9.0) | -15.8 (7.5) | 2.7 (6.7) |
| G4-C15 | -0.02 (0.32) | 0.02 (0.11) | 0.12 (0.38) | -3.9 (8.7) | -16.1 (7.1) | 1.6 (3.5) |
| U5-G14 | 2.25 (0.34) | -0.29 (0.26) | 0.22 (0.45) | -2.0 (8.7) | -12.6 (7.3) | 2.9 (7.1) |
| C6-G13 | -0.03 (0.32) | 0.00 (0.11) | -0.02 (0.36) | 3.7 (8.3) | -13.7 (7.0) | 1.0 (3.2) |
| A7-U12 | 0.14 (0.31) | 0.03 (0.14) | 0.05 (0.44) | 1.1 (8.8) | -12.4 (7.7) | 1.1 (6.1) |
| G8-C11 | -0.12 (0.41) | 0.01 (0.18) | 0.05 (0.44) | -1.7 (11.6) | -15.6 (9.2) | 2.7 (5.3) |
| U9-A10 | -0.15 (0.28) | -0.04 (010) | -0.14 (0.52) | 0.5 (13.8) | -10.9 (11.5) | 1.6 (5.1) |

(B) U-U

| Base pair | **Shear [Å]** | **Stretch [Å]** | **Stagger [Å]** | **Buckle [°]** | **Propeller [°]** | **Opening [°]** |
|---|---|---|---|---|---|---|
| U1-A18 | -0.20 (0.28) | -0.03 (0.11) | 0.14 (0.52) | -1.5 (15.3) | -9.8 (10.6) | 2.7 (4.7) |
| C2-G17 | -0.02 (0.35) | 0.04 (0.12) | -0.09 (0.41) | 7.9 (11.0) | -14.3 (8.3) | 1.7 (3.8) |
| A3-U16 | 0.13 (0.31) | 0.02 (0.13) | 0.01 (0.44) | 1.7 (8.9) | -13.5 (7.5) | 1.5 (5.9) |
| G4-C15 | -0.05 (0.38) | 0.01 (0.16) | -0.11 (0.45) | -5.3 (10.8) | -18.5 (7.6) | 3.2 (5.0) |
| U5-U14 | -0.14 (2.67) | -1.31 (0.64) | -0.06 (0.64) | 0.0 (14.0) | -17.7 (8.7) | -4.9 (13.7) |
| C6-G13 | 0.02 (0.34) | 0.01 (0.12) | -0.10 (0.42) | 7.3 (10.3) | -17.8 (7.5) | 2.7 (3.9) |
| A7-U12 | 0.15 (0.31) | 0.02 (0.14) | -0.04 (0.44) | -1.7 (9.3) | -13.4 (7.8) | 1.1 (6.2) |
| G8-C11 | -0.07 (0.38) | 0.02 (0.16) | 0.01 (0.44) | -4.7 (11.7) | -15.3 (9.3) | 2.4 (4.6) |
| U9-A10 | -0.15 (0.28) | -0.04 (0.10) | -0.11 (0.51) | -0.2 (13.9) | -10.4 (11.6) | 1.7 (5.1) |

(C) U-C

| Base pair | **Shear [Å]** | **Stretch [Å]** | **Stagger [Å]** | **Buckle [°]** | **Propeller [°]** | **Opening [°]** |
|---|---|---|---|---|---|---|
| U1-A18 | -0.19 (0.28) | -0.03 (0.11) | 0.13 (0.51) | -2.1 (16.3) | -10.5 (10.9) | 2.5 (4.8) |
| C2-G17 | -0.01 (0.33) | 0.04 (0.12) | -0.05 (0.43) | 6.3 (12.8) | -15.0 (8.2) | 1.9 (3.6) |
| A3-U16 | 0.12 (0.31) | 0.01 (0.13) | 0.13 (0.46) | 1.0 (9.4) | -15.1 (7.5) | 2.3 (6.2) |
| G4-C15 | 0.01 (0.35) | 0.00 (0.14) | 0.19 (0.54) | -0.9 (12.6) | -17.4 (7.8) | 1.1 (4.4) |
| U5-C14 | 1.13 (2.90) | -0.80 (2.86) | -1.48 (3.02) | -0.6 (18.9) | -22.2 (26.0) | -31.5 (32.5) |
| C6-G13 | -0.05 (0.34) | 0.02 (0.12) | -0.03 (0.44) | 4.4 (10.3) | -14.6 (7.9) | 1.3 (3.7) |
| A7-U12 | 0.12 (0.31) | 0.02 (0.15) | 0.08 (0.45) | 1.0 (9.5) | -13.6 (8.0) | 2.2 (6.6) |
| G8-C11 | -0.13 (0.40) | 0.01 (0.18) | 0.05 (0.44) | -2.6 (11.8) | -16.3 (9.2) | 2.8 (5.2) |
| U9-A10 | -0.14 (0.28) | -0.04 (0.10) | -0.12 (0.51) | 0.2 (13.9) | -11.1 (11.6) | 1.6 (5.1) |

(D) Ψ-G

| Base pair | **Shear [Å]** | **Stretch [Å]** | **Stagger [Å]** | **Buckle [°]** | **Propeller [°]** | **Opening [°]** |
|---|---|---|---|---|---|---|
| U1-A18 | -0.21 (0.27) | -0.03 (0.10) | 0.14 (0.50) | 0.1 (14.5) | -8.2 (10.9) | 2.3 (4.7) |
| C2-G17 | -0.03 (0.33) | 0.04 (0.12) | -0.11 (0.40) | 9.7 (10.3) | -14.8 (8.0) | 1.6 (3.5) |
| A3-U16 | 0.03 (0.56) | 0.00 (0.15) | 0.11 (0.42) | 2.2 (8.5) | -15.8 (7.1) | 3.1 (6.1) |
| G4-C15 | -0.04 (0.32) | -0.00 (0.11) | 0.07 (0.38) | -4.8 (8.8) | -17.2 (6.9) | 1.1 (3.5) |
| Ψ5-G14 | 2.26 (0.35) | -0.20 (0.35) | 0.39 (0.45) | -5.6 (8.6) | -10.9 (6.9) | 3.1 (9.4) |
| C6-G13 | -0.03 (0.32) | 0.01 (0.11) | 0.02 (0.36) | 3.1 (8.1) | -13.5 (6.9) | 1.0 (3.2) |
| A7-U12 | 0.14 (0.31) | 0.02 (0.13) | 0.07 (0.44) | 1.0 (8.9) | -12.5 (7.6) | 1.3 (5.9) |
| G8-C11 | -0.10 (0.37) | 0.01 (0.15) | 0.05 (0.44) | -1.9 (11.5) | -15.7 (9.1) | 2.5 (4.5) |
| U9-A10 | -0.14 (0.28) | -0.04 (0.10) | -0.13 (0.51) | 0.5 (13.7) | -10.8 (11.5) | 1.6 (5.1) |

(E) Ψ-U

| Base pair | **Shear [Å]** | **Stretch [Å]** | **Stagger [Å]** | **Buckle [°]** | **Propeller [°]** | **Opening [°]** |
|---|---|---|---|---|---|---|
| U1-A18 | -0.20 (0.28) | -0.02 (0.11) | 0.09 (0.51) | -2.6 (15.3) | -7.5 (11.2) | 2.6 (4.7) |
| C2-G17 | 0.04 (0.36) | 0.03 (0.12) | -0.18 (0.45) | 9.0 (11.3) | -11.7 (8.7) | 1.7 (3.6) |
| A3-U16 | -1.26 (2.16) | -0.16 (0.34) | 0.03 (0.49) | 5.2 (9.5) | -12.6 (8.1) | 1.3 (6.9) |
| G4-C15 | -0.09 (0.36) | -0.01 (0.12) | -0.13 (0.44) | -3.7 (10.6) | -17.8 (7.6) | 2.8 (4.2) |
| Ψ5-U14 | 1.11 (2.32 ) | -1.45 (0.43) | 0.05 (0.57) | -7.1 (13.0) | -16.0 (8.3) | -2.6 (11.6) |
| C6-G13 | 0.05 (0.39) | 0.00 (0.15) | -0.12 (0.42) | 4.3 (10.0) | -16.9 (7.5) | 3.2 (4.8) |
| A7-U12 | 0.17 (0.31) | 0.03 (0.14) | -0.01 (0.43) | -2.6 (9.1) | -11.5 (7.8) | 1.1 (6.1) |
| G8-C11 | -0.07 (0.36) | 0.02 (0.14) | 0.04 (0.43) | -2.7 (11.5) | -14.3 (9.3) | 2.4 (4.3) |

| | | | | | | |
|---|---|---|---|---|---|---|
| U9-A10 | -0.15 (0.28) | -0.04 (0.10) | -0.14 (0.51) | 1.8 (14.1) | -9.9 (11.7) | 1.6 (5.1) |

(F) Ψ-C

| Base pair | **Shear [Å]** | **Stretch [Å]** | **Stagger [Å]** | **Buckle [°]** | **Propeller [°]** | **Opening [°]** |
|---|---|---|---|---|---|---|
| U1-A18 | -0.20 (0.28) | -0.03 (0.10) | 0.15 (0.51) | -1.2 (14.9) | -9.3 (10.6) | 2.5 (4.7) |
| C2-G17 | -0.03 (0.34) | 0.04 (0.12) | -0.10 (0.41) | 8.9 (10.7) | -14.5 (8.2) | 1.8 (3.6) |
| A3-U16 | 0.10 (0.30) | 0.01 (0.13) | 0.09 (0.43) | 2.6 (8.8) | -14.2 (7.4) | 2.6 (6.2) |
| G4-C15 | -0.04 (0.38) | 0.01 (0.15) | 0.03 (0.44) | -3.4 (10.9) | -16.5 (7.8) | 2.0 (4.8) |
| Ψ5-C14 | 2.07 (2.89) | -0.97 (0.72) | -0.27 (0.86) | -8.5 (15.2) | -15.5 (10.0) | -14.4 (16.9) |
| C6-G13 | 0.01 (0.35) | 0.01 (0.12) | 0.05 (0.41) | 1.2 (9.9) | -13.4 (7.7) | 1.6 (3.7) |
| A7-U12 | 0.15 (0.31) | 0.03 (0.15) | 0.06 (0.44) | 0.4 (9.1) | -12.7 (7.9) | 1.2 (6.3) |
| G8-C11 | -0.10 (0.38) | 0.01 (0.14) | 0.05 (0.44) | -2.4 (11.7) | -15.9 (9.2) | 2.5 (4.5) |
| U9-A10 | -0.15 (0.28) | -0.04 (0.10) | -0.12 (0.51) | 0.2 (13.8) | -11.0 (11.6) | 1.6 (5.1) |

**Table S9.** Average values and standard deviations (in parentheses) of local base-pair step parameters for duplexes (GUC context) containing (A) U-G, (B) U-U, (C) U-C respectively and duplexes (GΨC context) containing (D) Ψ-G, (E) Ψ-U, (F) Ψ-C mismatches respectively.

(A) U-G

| Base-pair step | Shift [Å] | Slide [Å] | Rise [Å] | Tilt [°] | Roll [°] | Twist [°] |
|---|---|---|---|---|---|---|
| U1-A18/C2-G17 | -0.24 (0.59) | -1.34 (0.47) | 3.07 (0.36) | 1.5 (5.1) | 6.3 (6.8) | 30.3 (4.1) |
| C2-G17/A3-U16 | 0.02 (0.67) | -1.45 (0.44) | 3.38 (0.43) | -3.0 (5.4) | 14.6 (6.5) | 29.1 (7.2) |
| A3-U16/G4-C15 | -0.16 (0.65) | -1.67 (0.39) | 3.41 (0.32) | -2.5 (4.8) | 9.9 (5.6) | 34.4 (7.2) |
| G4-C15/U5-G14 | 0.20 (0.52) | -1.59 (0.42) | 3.28 (0.22) | 1.6 (4.1) | 5.7 (4.4) | 38.4 (3.3) |
| U5-G14/C6-G13 | -0.25 (0.62) | -1.86 (0.39) | 2.94 (0.32) | 4.7 (4.1) | 9.0 (5.5) | 21.9 (3.5) |

| | | | | | | |
|---|---|---|---|---|---|---|
| C6-G13/A7-U12 | -0.07 (0.63) | -1.27 (0.35) | 3.33 (0.33) | -0.6 (4.1) | 13.3 (6.6) | 31.5 (3.0) |
| A7-U12/G8-C11 | 0.15 (0.81) | -1.62 (0.42) | 3.32 (0.34) | -0.7 (4.5) | 10.5 (6.8) | 29.4 (3.6) |
| G8-C11/U9-A10 | 0.24 (0.64) | -1.76 (0.61) | 3.26 (0.28) | 1.7 (4.6) | 6.3 (6.2) | 30.7 (4.4) |

(E) U-U

| Base-pair step | Shift [Å] | Slide [Å] | Rise [Å] | Tilt [°] | Roll [°] | Twist [°] |
|---|---|---|---|---|---|---|
| U1-A18/C2-G17 | -0.22 (0.62) | -1.34 (0.52) | 3.10 (0.37) | 0.9 (5.2) | 6.4 (7.0) | 30.7 (4.0) |
| C2-G17/A3-U16 | -0.12 (0.66) | -1.27 (0.37) | 3.43 (0.38) | -0.9 (4.2) | 14.6 (6.7) | 32.5 (3.3) |
| A3-U16/G4-C15 | 0.06 (0.75) | -1.59 (0.40) | 3.45 (0.34) | 0.1 (4.5) | 12.2 (6.1) | 28.9 (3.6) |
| G4-C15/U5-U14 | -0.42 (0.97) | -1.58 (0.56) | 3.13 (0.55) | -1.7 (6.9) | 8.2 (6.3) | 31.0 (11.8) |
| U5-U14/C6-G13 | 0.36 (0.98) | -1.44 (0.56) | 3.11 (0.52) | 1.5 (6.3) | 8.3 (5.9) | 31.7 (11.6) |
| C6-G13/A7-U12 | -0.21 (0.64) | -1.27 (0.36) | 3.48 (0.37) | -0.4 (4.3) | 15.9 (6.9) | 31.9 (3.3) |
| A7-U12/G8-C11 | 0.21 (0.80) | -1.58 (0.41) | 3.33 (0.34) | -0.4 (4.6) | 10.9 (6.8) | 29.5 (3.5) |
| G8-C11/U9-A10 | 0.27 (0.64) | -1.68 (0.61) | 3.22 (0.28) | 1.5 (4.6) | 6.4 (6.3) | 30.1 (4.4) |

(C) U-C

| Base-pair step | Shift [Å] | Slide [Å] | Rise [Å] | Tilt [°] | Roll [°] | Twist [°] |
|---|---|---|---|---|---|---|
| U1-A18/C2-G17 | -0.19 (0.59) | -1.31 (0.56) | 3.13 (0.37) | 0.7 (5.2) | 6.6 (7.0) | 31.0 (4.0) |
| C2-G17/A3-U16 | -0.05 (0.67) | -1.23 (0.42) | 3.39 (0.39) | -0.9 (4.4) | 14.4 (6.6) | 32.5 (3.3) |
| A3-U16/G4-C15 | -0.07 (0.72) | -1.50 (0.51) | 3.30 (0.36) | -0.5 (4.5) | 10.0 (6.5) | 30.4 (3.4) |
| G4-C15/U5-C14 | -2.31 (2.14) | -1.34 (0.63) | 3.32 (0.65) | 6.3 (17.8) | 13.8 (9.6) | 36.0 (13.1) |
| U5-C14/C6-G13 | 2.24 (2.11) | -1.60 (0.58) | 3.03 (0.88) | -3.0 (18.9) | 12.4 (9.1) | 25.4 (15.5) |
| C6-G13/A7-U12 | -0.08 (0.64) | -1.25 (0.38) | 3.34 (0.38) | -1.2 (4.5) | 14.2 (7.0) | 31.7 (3.4) |
| A7-U12/G8-C11 | 0.05 (0.82) | -1.61 (0.43) | 3.33 (0.34) | -0.7 (4.6) | 10.8 (6.6) | 29.5 (3.7) |

| | | | | | | |
|---|---|---|---|---|---|---|
| G8-C11/U9-A10 | 0.23 (0.64) | -1.69 (0.61) | 3.25 (0.28) | 1.5 (4.6) | 6.5 (6.2) | 30.7 (4.4) |

(D) Ψ-G

| Base-pair step | Shift [Å] | Slide [Å] | Rise [Å] | Tilt [°] | Roll [°] | Twist [°] |
|---|---|---|---|---|---|---|
| U1-A18/C2-G17 | -0.24 (0.61) | -1.33 (0.50) | 3.10 (0.36) | 0.8 (5.2) | 5.4 (7.1) | 30.8 (4.0) |
| C2-G17/A3-U16 | 0.05 (0.66) | -1.34 (0.36) | 3.42 (0.37) | -1.5 (4.2) | 13.9 (6.3) | 32.3 (3.6) |
| A3-U16/G4-C15 | -0.29 (0.69) | -1.67 (0.38) | 3.38 (0.30) | -1.2 (4.3) | 10.5 (5.7) | 31.5 (3.6) |
| G4-C15/Ψ5-G14 | 0.35 (0.62) | -1.59 (0.39) | 3.33 (0.23) | 0.3 (4.0) | 6.8 (4.4) | 38.5 (3.3) |
| Ψ5-G14/C6-G13 | -0.21 (0.73) | -1.92 (0.37) | 2.84 (0.31) | 5.6 (4.0) | 7.7 (5.3) | 21.6 (3.4) |
| C6-G13/A7-U12 | -0.05 (0.63) | -1.29 (0.35) | 3.31 (0.33) | -0.4 (4.0) | 12.8 (6.5) | 31.6 (3.0) |
| A7-U12/G8-C11 | 0.11 (0.77) | -1.62 (0.41) | 3.33 (0.34) | -0.6 (4.5) | 10.3 (6.6) | 29.5 (3.4) |
| G8-C11/U9-A10 | 0.25 (0.62) | -1.75 (0.61) | 3.26 (0.28) | 1.6 (4.5) | 6.3 (6.2) | 30.7 (4.4 |

(E) Ψ-U

| Base-pair step | Shift [Å] | Slide [Å] | Rise [Å] | Tilt [°] | Roll [°] | Twist [°] |
|---|---|---|---|---|---|---|
| U1-A18/C2-G17 | -0.21 (0.59) | -1.29 (0.52) | 3.07 (0.36) | 1.3 (5.1) | 6.6 (6.9) | 29.7 (4.1) |
| C2-G17/A3-U16 | -0.15 (0.68) | -1.42 (0.45) | 3.31 (0.45) | -3.4 (5.6) | 14.1 (6.7) | 27.6 (8.1) |
| A3-U16/G4-C15 | -0.04 (0.70) | -1.60 (0.40) | 3.53 (0.35) | -1.7 (5.1) | 11.3 (5.8) | 34.1 (8.0) |
| G4-C15/Ψ5-U14 | -0.21 (0.86) | -1.59 (0.56) | 3.43 (0.51) | -1.4 (6.3) | 8.8 (5.7) | 37.2 (10.5) |
| Ψ5-U14/C6-G13 | 0.25 (0.86) | -1.67 (0.56) | 2.89 (0.45) | 4.4 (6.1) | 6.3 (5.6) | 26.6 (10.5) |
| C6-G13/A7-U12 | 0.05 (0.39) | 0.00 (0.15) | -0.12 (0.42) | 4.3 (10.0) | -16.9 (7.5) | 31.0 (3.5) |
| A7-U12/G8-C11 | 0.16 (0.76) | -1.59 (0.41) | 3.28 (0.34) | -0.5 (4.5) | 9.7 (6.6) | 28.9 (3.5) |
| G8-C11/U9-A10 | 0.25 (0.63) | -1.72 (0.62) | 3.23 (0.28) | 1.6 (4.5) | 6.2 (6.2) | 29.9 (4.5) |

(F) Ψ-C

| Base-pair step | Shift [Å] | Slide [Å] | Rise [Å] | Tilt [°] | Roll [°] | Twist [°] |
|---|---|---|---|---|---|---|
| U1-A18/C2-G17 | -0.20 (0.59) | -1.33 (0.50) | 3.08 (0.36) | 1.0 (5.1) | 6.0 (6.9) | 30.7 (4.0) |
| C2-G17/A3-U16 | -0.02 (0.68) | -1.31 (0.37) | 3.41 (0.37) | -1.4 (4.2) | 14.0 (6.5) | 32.4 (3.3) |
| A3-U16/G4-C15 | -0.10 (0.76) | -1.68 (0.40) | 3.38 (0.34) | -0.7 (4.5) | 10.9 (5.9) | 30.5 (3.7) |
| G4-C15/Ψ5-C14 | -1.08 (1.35) | -1.55 (0.46) | 3.54 (0.49) | 2.4 (5.7) | 10.4 (5.8) | 39.9 (11.7) |
| Ψ5-C14/C6-G13 | 1.07 (1.20) | -1.77 (0.54) | 2.96 (0.59) | 2.1 (7..4) | 7.6 (6.8) | 22.1 (12.1) |
| C6-G13/A7-U12 | -0.13 (0.65) | -1.25 (0.37) | 3.28 (0.36) | -0.0 (4.2) | 13.1 (6.9) | 31.2 (3.4) |
| A7-U12/G8-C11 | 0.15 (0.78) | -1.59 (0.42) | 3.32 (0.34) | -0.5 (4.5) | 10.6 (6.6) | 29.4 (3.6) |
| G8-C11/U9-A10 | 0.26 (0.62) | -1.72 (0.61) | 3.25 (0.29) | 1.7 (4.6) | 6.4 (6.3) | 30.6 (4.4) |

**Table S10.** Frequency (in %) of hydrogen bonds between $m^1\Psi$ (5) and A (14) for the duplexes containing $m^1\Psi$-A pair.

| Duplex | Hydrogen bonds | Hydrogen bonding atoms | | | Frequency (in %) |
|---|---|---|---|---|---|
| | | Donor | Donor-H | Acceptor | |
| **Duplex-Gm$^1\Psi$C** | N3-HN3- - - N1 | N3 ($m^1\Psi$5) | HN3 ($m^1\Psi$5) | N1 (A14) | r1 65.85<br>r2 67.86 |
| | N6-H61- - - O2 | N6 (A14) | H61 (A14) | O2 ($m^1\Psi$5) | r1 53.87<br>r2 56.13 |
| **Duplex-Cm$^1\Psi$G** | N3-HN3- - - N1 | N3 ($m^1\Psi$5) | HN3 ($m^1\Psi$5) | N1 (A14) | r1 66.73<br>r2 66.59 |
| | N6-H61- - - O2 | N6 (A14) | H61 (A14) | O2 ($m^1\Psi$5) | r1 62.33<br>r2 62.89 |
| **Duplex-Um$^1\Psi$A** | N3-HN3- - - N1 | N3 ($m^1\Psi$5) | HN3 ($m^1\Psi$5) | N1 (A14) | r1 65.57<br>r2 67.64 |
| | N6-H61- - - O2 | N6 (A14) | H61 (A14) | O2 ($m^1\Psi$5) | r1 56.74 |

| | | | | | |
|---|---|---|---|---|---|
| | | | | | r2 59.66 |
| **Duplex-Am$^1\Psi$U** | N3-HN3- - - N1 | N3 (m$^1\Psi$5) | HN3 (m$^1\Psi$5) | N1 (A14) | r1 67.73<br>r2 64.50 |
| | N6-H61- - - O2 | N6 (A14) | H61 (A14) | O2 (m$^1\Psi$5) | r1 59.61<br>r2 57.43 |

**Table S11.** Frequency (in %) of hydrogen bonds (A) between m$^1\Psi$ (5) and G/U/C (14) for the duplexes containing m$^1\Psi$-G, m$^1\Psi$-U, and m$^1\Psi$-C mismatches respectively, (B) between U (5) and G/U/C (14) for the duplexes containing U-G, U-U, and U-C mismatches respectively, and (C) between Ψ (5) and G/U/C (14) for the duplexes containing Ψ-G, Ψ-U, and Ψ-C mismatches respectively,

| (A) | Hydrogen bonds | Hydrogen bonding atoms | | | Frequency (in %) |
|---|---|---|---|---|---|
| | | Donor | Donor-H | Acceptor | |
| **Duplex-Gm$^1\Psi$C (m$^1\Psi$-G)** | N1-H1- - - O4 | N1 (G14) | H1 (G14) | O4 (m$^1\Psi$5) | r1 81.32<br>r2 83.56 |
| | N3-HN3- - - O6 | N3 (m$^1\Psi$5) | HN3 (m$^1\Psi$5) | O6 (G14) | r147.76<br>r2 58.62 |
| **Duplex-Gm$^1\Psi$G (m$^1\Psi$-U)** | N3-HN3- - - O4 | N3 (m$^1\Psi$5) | HN3 (m$^1\Psi$5) | O4 (U14) | r1 39.33<br>r2 33.36 |
| | N3-H3- - - O2 | N3 (U14) | H3 (U14) | O2 (m$^1\Psi$5) | r1 25.49<br>r2 30.53 |
| | N3-H3- - - O4 | N3 (U14) | H3 (U14) | O4 (m$^1\Psi$5) | r1 11.58<br>r2 7.50 |
| | N3-HN3- - - O2 | N3 (m$^1\Psi$5) | HN3 (m$^1\Psi$5) | O2 (U14) | r1 4.59<br>r2 4.70 |
| **Duplex-Gm$^1\Psi$C (m$^1\Psi$-C)** | N4-H41- - - O2 | N4 (C14) | H41 (C14) | O2 (m$^1\Psi$5) | r1 29.18<br>r2 32.42 |
| | N4-H41- - - O4 | N4 (C14) | H41 (C14) | O4 (m$^1\Psi$5) | r1 11.17<br>r2 11.85 |

| (B) | Hydrogen bonds | Hydrogen bonding atoms | | | Frequency |
|---|---|---|---|---|---|

| | | Donor | Donor-H | Acceptor | (in %) |
|---|---|---|---|---|---|
| **Duplex-GUC (U-G)** | N1-H1- - - O2 | N1 (G14) | H1 (G14) | O2 (U5) | 83.26 |
| | N3-HN3- - - O6 | N3 (U5) | H3 (U5) | O6 (G14) | 68.90 |
| **Duplex-GUG (U-U)** | N3-H3- - - O4 | N3 (U14) | H3 (U14) | O4 (U5) | 33.07 |
| | N3-H3- - - O4 | N3 (U5) | H3 (U5) | O4 (U14) | 30.12 |
| | N3-H3- - - O2 | N3 (U5) | H3 (U5) | O2 (U14) | 2.97 |
| | N3-H3- - - O2 | N3 (U14) | H3 (U14) | O2 (U5) | 2.73 |
| **Duplex-GUC (U-C)** | N4-H41- - - O4 | N4 (C14) | H41 (C14) | O4 (U5) | 30.34 |
| | N4-H41- - - O2 | N4 (C14) | H41 (C14) | O2 (U5) | 7.32 |

| (C) | Hydrogen bonds | Hydrogen bonding atoms | | | Frequency (in %) |
|---|---|---|---|---|---|
| | | Donor | Donor-H | Acceptor | |
| **Duplex-GΨC (Ψ-G)** | N1-H1- - - O4 | N1 (G14) | H1 (G14) | O4 (Ψ5) | 84.48 |
| | N3-HN3- - - O6 | N3 (Ψ5) | HN3 (Ψ5) | O6 (G14) | 57.98 |
| **Duplex-GΨG (Ψ-U)** | N3-HN3- - - O4 | N3 (Ψ5) | HN3 (Ψ5) | O4 (U14) | 49.02 |
| | N3-H3- - - O2 | N3 (U14) | H3 (U14) | O2 (Ψ5) | 18.81 |
| | N3-H3- - - O4 | N3 (U14) | H3 (U14) | O4 (Ψ5) | 9.82 |
| | N3-HN3- - - O2 | N3 (Ψ5) | HN3 (Ψ5) | O2 (U14) | 3.64 |
| **Duplex-GΨC (Ψ-C)** | N4-H41- - - O2 | N4 (C14) | H41 (C14) | O2 (Ψ5) | 23.92 |
| | N4-H41- - - O4 | N4 (C14) | H41 (C14) | O4 (Ψ5) | 15.52 |

**Table S12.** Water bridges (occupancy in %) formed by the backbone phosphate oxygen atoms of $m^1\Psi$ with the phosphate oxygen atoms of its immediate neighboring residues for the duplexes of four different sequence contexts containing $m^1\Psi$-A pair.

| Water-bridges | Duplex-Gm¹ΨC | Duplex-Cm¹ΨG | Duplex-Um¹ΨA | Duplex-Am¹ΨU |
|---|---|---|---|---|
| OP2(5)-W-OP1(4) | 17.82<br>16.71 | 16.32<br>14.87 | 17.13<br>17.82 | 14.69<br>14.81 |
| OP2(5)-W-OP2(4) | 10.64<br>11.09 | 13.94<br>13.43 | 16.67<br>15.54 | 10.66<br>10.25 |
| OP2(6)-W-OP1(5) | 16.88<br>15.23 | 8.62<br>8.32 | 6.70<br>6.38 | 13.11<br>12.94 |
| OP2(6)-W-OP2(5) | 11.60<br>9.55 | 7.73<br>6.67 | 5.84<br>5.62 | 10.24<br>9.74 |
| Values obtained from replicate 1 and replicate 2 are in the top and bottom respectively. | | | | |

**Table S13.** Water bridges (occupancy in %) formed by (A) the backbone phosphate oxygen atoms of $m^1\Psi$ with the phosphate oxygen atoms of its immediate neighboring residues for the duplexes ($Gm^1\Psi C$ context) containing $m^1\Psi$-G, $m^1\Psi$-U, and $m^1\Psi$-C mismatches respectively; (B) backbone phosphate oxygen atoms of U with the phosphate oxygen atoms of its immediate neighboring residues for the duplexes (GUC context) containing U-G, U-U, and U-C mismatches respectively; (C) backbone phosphate oxygen atoms of Ψ with the phosphate oxygen atoms of its immediate neighboring residues for the duplexes (GΨC context) containing Ψ-G, Ψ-U, and Ψ-C mismatches respectively.

(A)

| Water-bridges | Duplex-Gm¹ΨC (m1Ψ-G) | Duplex-Gm¹ΨC (m1Ψ-U) | Duplex-Gm¹ΨC (m¹Ψ-C) |
|---|---|---|---|
| OP2(5)-W-OP1(4) | 50.61<br>30.02 | 15.67<br>20.49 | 20.47<br>26.02 |
| OP2(5)-W-OP2(4) | 10.39<br>13.04 | 7.34<br>7.54 | 7.09<br>6.52 |
| OP1(6)-W-OP1(5) | <5<br><5 | 5.07<br><5 | <5<br><5 |
| OP2(6)-W-OP1(5) | <5<br><5 | 27.19<br>23.90 | 19.07<br>20.54 |
| OP2(6)-W-OP2(5) | <5<br><5 | 14.47<br>11.01 | 11.68<br>10.91 |

| Values obtained from replicate 1 and replicate 2 are in the top and bottom respectively. |
|---|

(B)

| **Water-bridges** | Duplex-GUC (U-A) [5] | Duplex-GUC (U-G) | Duplex-GUC (U-U) | Duplex-GUC (U-C) |
|---|---|---|---|---|
| OP2(5)-W-OP1(4) | 16 | 31.05 | 11.84 | 13.40 |
| OP2(5)-W-OP2(4) | 11 | 14.72 | 7.6 | 7.67 |
| OP2(6)-W-OP1(5) | 15 | <5 | 32.37 | 28.32 |
| OP2(6)-W-OP2(5) | 12 | 5.47 | 18.02 | 21.47 |

(C)

| **Water-bridges** | Duplex-GΨC (Ψ-A) [6] | Duplex-GΨC (Ψ-G) | Duplex-GΨC (Ψ-U) | Duplex-GΨC (Ψ-C) |
|---|---|---|---|---|
| OP2(5)-W-HN1(5) | 28 | 18.22 | 15.59 | 23.08 |
| OP2(5)-W-OP1(4) | 12 | 37.8 | 26.76 | 25.01 |
| OP2(5)-W-OP2(4) | 7 | 18.53 | 10.94 | 13.4 |
| OP2(6)-W-OP1(5) | 21 | <5 | 30.86 | 24.53 |
| OP2(6)-W-OP2(5) | 16 | <5 | 19.15 | 19.19 |

**Table S14.** Water bridging interaction between the bases of 5th and 14th residues in the duplexes (containing mismatches) under this study.

| Base pair | Water bridges | Occupancy (%) |
|---|---|---|
| U-G | O4(5)-W-O6(14) | 10.92 |
| | O2(5)-W-H21(14) (indicated as O2(5)-W-N2(14) in the main text) | 12.14 |
| | O2’-O2(5)-W-H21(14) (indicated as O2’-O2(5)-W-N2(14) in the main text) i.e a water molecule bridging O2’ and O2 atoms of U and | 2.94 |

| | | |
|---|---|---|
| | N2 atom of G | |
| Ψ-G | O2(5)-W-O6(14) | 9.01 |
| | O4(5)-W-H21(14)<br>(indicated as O4(5)-W-N2(14) in the main text) | 20.96 |
| | O2'-O4(5)-W-H21(14)<br>(indicated as O2'-O2(5)-W-N2(14) in the main text) i.e a water molecule bridging O2' and O4 atoms of Ψ and N2 atom of G | 4.96 |
| $m^1$Ψ-G | O2(5)-W-O6(14) | 9.31<br>9.71 |
| | O4(5)-W-H22(14)<br>(indicated as O4(5)-W-N2(14) in the main text) | 10.88<br>14.64 |
| | O2'-O4(5)-W-H22(14)<br>(indicated as O2'-O2(5)-W-N2(14) in the main text) i.e a water molecule bridging O2' and O4 atoms of $m^1$Ψ and N2 atom of G | 3.37<br>4.18 |
| U-U | O2(5)-W-O2(14) | 9.36 |
| | O4(5)-W-O4(14) | 6.60 |
| | O2(5)-W-H3(14)<br>(indicated as O2(5)-W-N3(14) in the main text) | 6.19 |
| | O2(14)-W-H3(5)<br>(indicated as O2(14)-W-N3(5) in the main text) | 4.37 |
| | O4(5)-W-H3(14)<br>(indicated as O4(5)-W-N3(14) in the main text) | 0.48 |
| | O4(14)-W-H3(5)<br>(indicated as O4(14)-W-N3(5) in the main text) | 0.44 |
| Ψ-U | O4(5)-W-O2(14) | 15.48 |
| | O2(5)-W-O4(14) | 8.44 |
| | O4(5)-W-H3(14)<br>(indicated as O4(5)-W-N3(14) in the main text) | 4.88 |
| | O4(14)-W-HN3(5)<br>(indicated as O4(14)-W-N3(5) in the main text) | 0.44 |

| | | |
|---|---|---|
| | O2(14)-W-HN3(5)<br>(indicated as O2(14)-W-N3(5) in the main text) | 1.69 |
| | O2(5)-W-H3(14)<br>(indicated as O2(5)-W-N3(14) in the main text) | 0.25 |
| m[1]Ψ-U | O4(5)-W-O2(14) | 10.60<br>10.82 |
| | O2(5)-W-O4(14) | 7.67<br>9.02 |
| | O4(5)-W-H3(14)<br>(indicated as O4(5)-W-N3(14) in the main text) | 3.77<br>3.76 |
| | O4(14)-W-HN3(5)<br>(indicated as O4(14)-W-N3(5) in the main text) | 0.41<br>0.48 |
| | O2(14)-W-HN3(5)<br>(indicated as O2(14)-W-N3(5) in the main text) | 4.03<br>5.10 |
| | O2(5)-W-H3(14)<br>(indicated as O2(5)-W-N3(14) in the main text) | 1.12<br>0.95 |
| U-C | O4(5)-W-N3(14) | 5.24 |
| | O2(5)-W-N3(14) | 8.08 |
| | H3(5)-W-N3(14)<br>(indicated as N3(5)-W-N3(14) in the main text) | 10.82 |
| | O4(5)-W-H42(14) | 1.80 |
| Ψ-C | O2(5)-W-N3(14) | 0.42 |
| | O4(5)-W-N3(14) | 16.47 |
| | HN3(5)-W-N3(14)<br>(indicated as N3(5)-W-N3(14) in the main text) | 10.82 |
| | O2(5)-W-H42(14)<br>(indicated as O2(5)-W-N4(14) in the main text) | 1.64 |
| m[1]Ψ-C | O2(5)-W-N3(14) | 0.44<br>1.02 |
| | O4(5)-W-N3(14) | 10.47<br>11.54 |

| | | |
|---|---|---|
| | H3(5)-W-N3(14)<br>(indicated as N3(5)-W-N3(14) in the main text) | 17.17<br>15.48 |
| | O2(5)-W-H42(14)<br>(indicated as O2(5)-W-N4(14) in the main text) | 1.78<br>1.74 |

**Table S15.** Interaction energies [kcal/mol] of the base-pair steps containing the m¹Ψ-A base pair obtained from the last 400 ns of MD simulations at 300 K for (A) duplex-Gm¹ΨC, (B) duplex-Cm¹ΨG, (C) duplex-Um¹ΨA and (D) duplex-Am¹ΨU respectively

| **(A)** | **Coulomb (kcal/mol)** | **VDW (kcal/mol)** | **Total [a] (kcal/mol)** | **U-A pair [b]** | **Ψ-A pair [c]** |
|---|---|---|---|---|---|
| **Stacking energies** | | | | | |
| **Step 1**<br>**4 5**<br>**5'- G m¹Ψ -3'**<br>**3'- C A -5'**<br>**15 14** | -1.07<br>(1.53)<br><br>-0.99<br>(1.52) | -14.91<br>(1.03)<br><br>-14.96<br>(0.99) | **-15.98**<br>**(1.84)**<br><br>**-15.95**<br>**(1.81)** | -9.57 (2.6)<br>-12.00 | -10.58 (2.51)<br>-12.48 |
| **Step 2**<br>**5 6**<br>**5'- m¹Ψ C -3'**<br>**3'- A G -5'**<br>**14 13** | 4.14<br>(1.46)<br><br>4.32<br>(1.43) | -14.47<br>(1.04)<br><br>-14.62<br>(0.94) | **-10.33**<br>**(3.21)**<br><br>**-10.3**<br>**(1.71)** | -9.45 (2.33)<br>-12.66 | -9.84 (2.3)<br>-11.99 |
| **Base pairing energies** | | | | | |
| **4 15**<br>**G - C** | -26.51<br>(2.53)<br><br>-26.48<br>(2.57) | 0.42<br>(1.70)<br><br>0.43<br>(1.71) | **-26.09**<br>**(3.04)**<br><br>**-26.05**<br>**(3.08)** | -26.0 (3.10)<br>-23.31 | -25.93 (3.10)<br>-23.18 |
| **5 14**<br>**m¹Ψ - A** | -9.66<br>(2.44)<br><br>-9.95<br>(1.93) | -0.56<br>(1.29)<br><br>-0.58<br>(1.31) | **-10.22**<br>**(2.76)**<br><br>**-10.53**<br>**(2.33)** | -9.24 (2.48)<br>-12.66 | -8.99 (2.45)<br>-12.33 |
| **6 13**<br>**C - G** | -26.38<br>(2.58) | 0.32<br>(1.69) | **-26.06**<br>**(3.08)** | -26.08 (3.03)<br>-24.02 | -26.10 (3.0)<br>-23.93 |

| | | | |
|---|---|---|---|
| | -26.35<br>(2.58) | 0.29<br>(1.67) | **-26.06<br>(2.59)** |

[a] The values obtained in this study from the last 400 ns of MD simulations
[b] The Values reported in Deb et al [5] for the U-A pair
[c] The Values reported in Deb et al [5] for the Ψ-A pair
The values corresponding to the QM energies reported in Deb et al [5] are shown in blue.
For the energies corresponding to $m^1\Psi$-A, the values obtained from replicate 1 and replicate 2 are in the top and bottom respectively.

| **(B)** | **Coulomb (kcal/mol)** | **VDW (kcal/mol)** | **Total [a] (kcal/mol)** | **U-A pair [b]** | **Ψ-A pair [c]** |
|---|---|---|---|---|---|
| **Stacking energies** | | | | | |
| **Step 1**<br>**4 5**<br>**5'- C $m^1\Psi$ -3'**<br>**3'- G A -5'**<br>**15 14** | 3.16<br>(1.22)<br><br>3.20<br>(1.22) | -14.94<br>(0.95)<br><br>-14.97<br>(0.94) | **-11.78<br>(1.54)**<br><br>**-11.74<br>(1.54)** | -8.34 (2.18)<br>-12.31 | -7.9 (2.03)<br>-12.23 |
| **Step 2**<br>**5 6**<br>**5'- $m^1\Psi$ G -3'**<br>**3'- A C -5'**<br>**14 13** | 1.03<br>(1.53)<br><br>0.94<br>(1.52) | -14.34<br>(0.96)<br><br>-14.36<br>(0.95) | **-13.31<br>(1.81)**<br><br>**-13.42<br>(1.79)** | -8.64 (2.35)<br>-12.71 | -10.24 (2.16)<br>-14.32 |
| **Base pairing energies** | | | | | |
| **4 15**<br>**C - G** | -26.61<br>(2.48)<br><br>-26.62<br>(2.47) | 0.4<br>(1.69)<br><br>0.39<br>(1.69) | **-26.21<br>(3.00)**<br><br>**-26.23<br>(2.99)** | -25.99 (3.08)<br>-23.42 | -25.94 (3.08)<br>-23.57 |
| **5 14**<br>**$m^1\Psi$ - A** | -10.28<br>(1.80)<br><br>-10.29<br>(1.79) | -0.52<br>(1.31)<br><br>-0.52<br>(1.31) | **-10.80<br>(2.23)**<br><br>**-10.81<br>(2.22)** | -9.65 (2.37)<br>-12.77 | -9.46 (2.24)<br>-12.38 |
| **6 13**<br>**G - C** | -26.32<br>(2.52) | 0.26<br>(1.69) | **-26.06<br>(3.03)** | -26.05 (3.01)<br>-24.06 | -26.24 (2.50)<br>-24.06 |

| | | | |
|---|---|---|---|
| | -26.29<br>(2.51) | 0.25<br>(1.67) | **-26.04<br>(3.01)** |

[a] The values obtained in this study from the last 400 ns of MD simulations
[b] The values reported in Deb et al [5] for the U-A pair
[c] The values reported in Deb et al, 2019 [5] for the Ψ-A pair
The values corresponding to the QM energies reported in Deb et al [5] are shown in blue.
For the energies corresponding to $m^1\Psi$-A, the values obtained from replicate 1 and replicate 2 are in the top and bottom respectively.

| **(C)** | **Coulomb (kcal/mol)** | **VDW (kcal/mol)** | **Total [a] (kcal/mol)** | **U-A pair [b]** | **Ψ-A pair [c]** |
|---|---|---|---|---|---|
| **Stacking energies** | | | | | |
| **Step 1**<br>**4 5**<br>**5'- U $m^1\Psi$ -3'**<br>**3'- A A -5'**<br>**15 14** | 2.96<br>(1.18)<br><br>2.98<br>(1.19) | -14.13<br>(1.01)<br><br>-14.09<br>(1.05) | **-11.17<br>(1.55)**<br><br>**-11.11<br>(1.59)** | -6.5 (1.93)<br>-9.62 | -6.13 (1.88)<br>-9.54 |
| **Step 2**<br>**5 6**<br>**5'- $m^1\Psi$ A -3'**<br>**3'- A U -5'**<br>**14 13** | -0.26<br>(1.66)<br><br>-0.37<br>(1.65) | -13.48<br>(0.98)<br><br>-13.49<br>(0.98) | **-13.74<br>(1.93)**<br><br>**-13.86<br>(1.92)** | -10.42 (2.37)<br>-12.66 | -11.29 (3.16)<br>-13.62 |
| **Base pairing energies** | | | | | |
| **4 15**<br>**U - A** | -9.01<br>(1.94)<br><br>-9.02<br>(1.89) | -0.54<br>(1.31)<br><br>-0.54<br>(1.31) | **-9.55<br>(2.34)**<br><br>**-9.56<br>(2.29)** | -9.38 (2.37)<br>-12.19 | -9.36 (2.37)<br>-12.23 |
| **5 14**<br>**$m^1\Psi$ - A** | -10.06<br>(1.78)<br><br>-10.04<br>(1.77) | -0.68<br>(1.27)<br><br>-0.69<br>(1.26) | **-10.74<br>(2.19)**<br><br>**-10.73<br>(2.17)** | -9.63 (2.29)<br>-12.63 | -9.32 (2.25)<br>-12.17 |
| **6 13**<br>**A - U** | -9.06<br>(1.88)<br><br>-9.02 | -0.64<br>(1.26)<br><br>-0.64 | **-9.70<br>(2.26)**<br><br>**-9.66** | -9.70 (2.27)<br>-12.69 | -9.68 (2.29)<br>-12.65 |

| | | | |
|---|---|---|---|
| | (1.89) | (1.27) | **(2.28)** |

[a] The values obtained in this study from the last 400 ns of MD simulations
[b] The Values reported in Deb et al [5] for the U-A pair
[c] The Values reported in Deb et al [5] for the Ψ-A pair
The values corresponding to the QM energies reported in Deb et al [5] are shown in blue.
For the energies corresponding to m[1]Ψ-A, the values obtained from replicate 1 and replicate 2 are in the top and bottom respectively.

| **(D)** | **Coulomb (kcal/mol)** | **VDW (kcal/mol)** | **Total [a] (kcal/mol)** | **U-A pair [b]** | **Ψ-A pair [c]** |
|---|---|---|---|---|---|
| **Stacking energies** | | | | | |
| **Step 1**<br>**4 5**<br>**5'- A m[1]Ψ -3'**<br>**3'- U A -5'**<br>**15 14** | 0.24<br>(1.50)<br><br>0.26<br>(1.48) | -14.24<br>(1.04)<br><br>-14.28<br>(1.02) | **-14.00**<br>**(1.82)**<br><br>**-14.02**<br>**(1.79)** | -8.12 (2.16)<br>-11.44 | -8.39 (2.12)<br>-11.14 |
| **Step 2**<br>**5 6**<br>**5'- m[1]Ψ U -3'**<br>**3'- A A -5'**<br>**14 13** | 4.35<br>(1.03)<br><br>4.38<br>(1.04) | -13.75<br>(0.93)<br><br>-13.79<br>(0.92) | **-9.4**<br>**(1.39)**<br><br>**-9.41**<br>**(1.38)** | -6.59 (1.92)<br>-9.64 | -7.68 (1.83)<br>-10.69 |
| **Base pairing energies** | | | | | |
| **4 15**<br>**A - U** | -8.85<br>(2.03)<br><br>-8.89<br>(1.99) | -0.61<br>(1.28)<br><br>-0.59<br>(1.29) | **-9.46**<br>**(2.39)**<br><br>**–9.48**<br>**(2.37)** | -9.31 (2.49)<br>-12.39 | -9.38 (2.40)<br>-12.47 |
| **5 14**<br>**m[1]Ψ - A** | -10.08<br>(1.82)<br><br>-10.07<br>(1.83) | -0.57<br>(1.30)<br><br>-0.55<br>(1.32) | **-10.65**<br>**(2.23)**<br><br>**-10.62**<br>**(2.26)** | -9.41 (2.39)<br>-12.19 | -9.15 (2.32)<br>-12.02 |
| **6 13**<br>**U - A** | -8.92<br>(1.93)<br><br>-8.93<br>(1.92) | -0.69<br>(1.26)<br><br>-0.69<br>(1.25) | **-9.61**<br>**(2.30)**<br><br>**-9.62**<br>**(2.29)** | -9.67 (2.27)<br>-12.65 | -9.71 (2.26)<br>-12.77 |

| |
|---|
| [a] The values obtained in this study from the last 400 ns of MD simulations<br>[b] The Values reported in Deb et al [5] for U-A pair<br>[c] The Values reported in Deb et al [5] for Ψ-A pair<br>The values corresponding to the QM energies reported in Deb et al [5] are shown in blue.<br>For the energies corresponding to m[1]Ψ-A, the values obtained from replicate 1 and replicate 2 are in the top and bottom respectively. |

**Table S16.** Interaction energies of (A) the base-pair steps containing the m[1]Ψ-G mismatch for duplex-Gm[1]ΨC [kcal/mol]; (B) the base-pair steps containing the U-G mismatch for duplex-GUC [kcal/mol]; (C) the base-pair steps containing the Ψ-G mismatch for duplex-GΨC [kcal/mol]

| (A) | Coulomb (kcal/mol) | VDW (kcal/mol) | Total [a] (kcal/mol) |
|---|---|---|---|
| **Stacking energies** | | | |
| **Step 1**<br>**4 5**<br>**5'- G m[1]Ψ -3'**<br>**3'- C G -5'**<br>**15 14** | -1.58 (2.90)<br>-0.63 (2.64) | -15.18 (1.22)<br>-15.38 (1.12) | **-16.76 (3.14)**<br>**-16.01 (2.86)** |
| **Step 2**<br>**5 6**<br>**5'- m[1]Ψ C -3'**<br>**3'- G G -5'**<br>**14 13** | 7.34 (1.88)<br>7.19 (1.77) | -14.05 (1.06)<br>-14.26 (0.96) | **-6.71 (2.16)**<br>**-7.07 (2.02)** |
| **Base pairing energies** | | | |
| **4 15**<br>**G - C** | -26.23 (2.55)<br>-26.24 (2.63) | 0.23 (1.67)<br>0.28 (1.67) | **-26.00 (3.05)**<br>**-25.96 (3.12)** |
| **5 14**<br>**m[1]Ψ - G** | -12.21 (2.36)<br>-12.66 (2.13) | -0.48 (1.20)<br>-0.50 (1.24) | **-12.69 (2.64)**<br>**-13.16 (2.46)** |
| **6 13**<br>**C - G** | -26.63 (2.72)<br>-26.66 (2.46) | 0.55 (1.74)<br>0.51 (1.70) | **-26.08 (3.23)**<br>**-26.15 (2.99)** |
| [a] The values obtained in this study from the last 400 ns of MD simulations<br>The values obtained from replicate 1 and replicate 2 are in the top and bottom respectively. | | | |

| (B) | Coulomb (kcal/mol) | VDW (kcal/mol) | Total [a] |
|---|---|---|---|

| | | | (kcal/mol) |
|---|---|---|---|
| **Stacking energies** | | | |
| **Step 1**<br><br>**4 5**<br>**5'- G U -3'**<br>**3'- C G -5'**<br>**15 14** | 5.24 (2.40) | -14.94 (0.94) | **-9.7 (2.57)** |
| **Step 2**<br>**5 6**<br>**5'- U C -3'**<br>**3'- G G -5'**<br>**14 13** | 7.98 (2.02) | -14.27 (0.89) | **-6.29 (2.21)** |
| **Base pairing energies** | | | |
| **4 15**<br>**G - C** | -26.38 (2.49) | 0.34 (1.69) | **-26.04 (3.01)** |
| **5 14**<br>**U - G** | -12.51 (1.95) | -0.46 (1.29) | **-12.97 (2.33)** |
| **6 13**<br>**C - G** | -26.68 (2.43) | 0.55 (1.74) | **-26.13 (2.99)** |
| [a] The values obtained in this study from the last 400 ns of MD simulations | | | |

| **(C)** | **Coulomb (kcal/mol)** | **VDW (kcal/mol)** | **Total [a]**<br>**(kcal/mol)** |
|---|---|---|---|
| **Stacking energies** | | | |
| **Step 1**<br><br>**4 5**<br>**5'- G Ψ -3'**<br>**3'- C G -5'**<br>**15 14** | 4.21 (2.62) | -14.68 (1.03) | **-10.47 (2.81)** |
| **Step 2**<br>**5 6**<br>**5'- Ψ C -3'**<br>**3'- G G -5'**<br>**14 13** | 7.45 (1.88) | -14.16 (0.89) | **-6.71 (2.08)** |
| **Base pairing energies** | | | |

| | | | |
|---|---|---|---|
| 4 15<br>G - C | -26.46 (2.46) | 0.38 (1.69) | **-26.08 (2.98)** |
| 5 14<br>Ψ - G | -13.12 (1.95) | -0.44 (1.25) | **-13.56 (2.32)** |
| 6 13<br>C - G | -26.69 (2.41) | 0.53 (1.72) | **-26.16 (2.96)** |
| [a] The values obtained in this study from the last 400 ns of MD simulations | | | |

**Table S17.** Interaction energies of the base-pair steps containing the m$^1$Ψ-U mismatch for duplex-Gm$^1$ΨC [kcal/mol]. (A) The values obtained in this study from the last 400 ns of MD simulations. (B) Those obtained from the most populated cluster in the last 400 ns of MD simulations and (C) Those obtained from the second most populated cluster in the last 400 ns of MD simulations.

| **(A)** | **Coulomb (kcal/mol)** | **VDW (kcal/mol)** | **Total (kcal/mol)** |
|---|---|---|---|
| **Stacking energies** | | | |
| **Step 1**<br>4 5<br>5'- G m$^1$Ψ -3'<br>3'- C U -5'<br>15 14 | 1.21 (2.96)<br>0.88 (3.04) | -13.10 (1.26)<br>-13.01 (1.33) | **-11.89 (3.22)**<br>**-12.13 (3.32)** |
| **Step 2**<br>5 6<br>5'- m$^1$Ψ C -3'<br>3'- U G -5'<br>14 13 | 6.25 (2.36)<br>6.19 (2.31) | -12.87 (1.20)<br>-12.93 (1.25) | **-6.62 (2.65)**<br>**-6.74 (2.63)** |
| **Base pairing energies** | | | |
| 4 15<br>G - C | -26.37 (2.65)<br>-26.36 (2.62) | 0.56 (1.75)<br>0.56 (1.76) | **-25.81 (3.18)**<br>**-25.80 (3.16)** |
| 5 14<br>m$^1$Ψ - U | -6.21 (2.82)<br>-6.02 (2.84) | -1.05 (0.92)<br>-1.07 (0.89) | **-7.26 (2.97)**<br>**-7.09 (2.97)** |
| 6 13<br>C - G | -25.89 (2.95)<br>-25.87 (3.17) | 0.46 (1.73)<br>0.49 (1.75) | **-25.43 (3.42)**<br>**-25.38 (3.62)** |

| The values obtained in this study from the last 400 ns of MD simulations of replicate 1 and replicate 2 are in the top and bottom respectively. |
| --- |

| (B) | Coulomb (kcal/mol) | VDW (kcal/mol) | Total (kcal/mol) |
| --- | --- | --- | --- |
| **Stacking energies** | | | |
| **Step 1**<br><br>**4 5**<br>**5'- G m$^1$Ψ -3'**<br>**3'- C U -5'**<br>**15 14** | 2.91 (2.25) | -13.39 (1.04) | **-10.48 (2.47)** |
| **Step 2**<br>**5 6**<br>**5'- m$^1$Ψ C -3'**<br>**3'- U G -5'**<br>**14 13** | 5.32 (2.23) | -12.58 (0.97) | **-7.26 (2.43)** |
| **Base pairing energies** | | | |
| **4 15**<br>**G - C** | -26.34 (2.62) | 0.52 (1.74) | **-25.82 (3.14)** |
| **5 14**<br>**m$^1$Ψ - U** | -6.81 (2.36) | -1.00 (0.96) | **-7.81 (2.54)** |
| **6 13**<br>**C - G** | -25.95 (2.83) | 0.49 (1.73) | **-25.46 (3.32)** |
| The values obtained in this study from the most populated cluster in the last 400 ns of MD simulations of replicate 1. | | | |

| (C) | Coulomb (kcal/mol) | VDW (kcal/mol) | Total (kcal/mol) |
| --- | --- | --- | --- |
| **Stacking energies** | | | |
| **Step 1**<br><br>**4 5**<br>**5'- G m$^1$Ψ -3'**<br>**3'- C U -5'**<br>**15 14** | -1.17 (2.28) | -12.96 (1.29) | **-14.13 (2.61)** |
| **Step 2**<br>**5 6** | 7.71 (1.76) | -13.51 (1.08) | **-5.8 (2.06)** |

| | | | |
|---|---|---|---|
| **5'- m¹Ψ C -3'<br>3'- U G -5'<br>14 13** | | | |
| **Base pairing energies** | | | |
| **4 15<br>G - C** | -26.49 (2.63) | 0.68 (1.79) | **-25.81 (3.18)** |
| **5 14<br>m¹Ψ - U** | -6.19 (2.71) | -1.24 (0.87) | **-7.43 (2.84)** |
| **6 13<br>C - G** | -25.88 (2.77) | 0.44 (1.73) | **-25.44 (3.26)** |
| The values obtained in this study from the second most populated cluster in the last 400 ns of MD simulations of replicate 1. | | | |

**Table S18.** Interaction energies of the base-pair steps containing the U-U mismatch for duplex-GUC [kcal/mol]. (A) The values obtained in this study from the last 400 ns of MD simulations. (B) Those obtained from the most populated cluster in the last 400 ns of MD simulations and (C) Those obtained from the second most populated cluster in the last 400 ns of MD simulations.

| (A) | Coulomb (kcal/mol) | VDW (kcal/mol) | Total (kcal/mol) |
|---|---|---|---|
| **Stacking energies** | | | |
| **Step 1**<br><br>**4 5<br>5'- G U-3'<br>3'- C U -5'<br>15 14** | 7.37 (2.98) | -12.35 (1.12) | **-4.98 (3.18)** |
| **Step 2<br>5 6<br>5'- U C -3'<br>3'- U G -5'<br>14 13** | 7.93 (2.88) | -12.41 (1.15) | **-4.48 (3.10)** |
| **Base pairing energies** | | | |
| **4 15<br>G - C** | -26.07 (2.78) | 0.51 (1.76) | **-25.56 (3.29)** |
| **5 14** | -3.60 (2.79) | -0.99 (0.81) | **-4.59 (2.91)** |

| | | | |
|---|---|---|---|
| **U - U** | | | |
| **6 13<br>C - G** | -26.11 (2.61) | 0.48 (1.74) | **-25.63 (3.14)** |

| **(B)** | **Coulomb (kcal/mol)** | **VDW (kcal/mol)** | **Total (kcal/mol)** |
|---|---|---|---|
| **Stacking energies** | | | |
| **Step 1**<br><br>**4 5<br>5'- G U-3'<br>3'- C U -5'<br>15 14** | 5.51 (2.26) | -12.30 (0.96) | **-6.79 (2.46)** |
| **Step 2<br>5 6<br>5'- U C -3'<br>3'- U G -5'<br>14 13** | 9.71 (2.02) | -12.75 (1.04) | **-3.04 (2.27)** |
| **Base pairing energies** | | | |
| **4 15<br>G - C** | -26.00 (2.79) | 0.55 (1.75) | **-25.45 (3.29)** |
| **5 14<br>U - U** | -4.12 (2.24) | -1.06 (0.82) | **-5.18 (2.38)** |
| **6 13<br>C - G** | -26.23 (2.51) | 0.48 (1.74) | **-25.75 (3.05)** |

| **(C)** | **Coulomb (kcal/mol)** | **VDW (kcal/mol)** | **Total (kcal/mol)** |
|---|---|---|---|
| **Stacking energies** | | | |
| **Step 1**<br><br>**4 5<br>5'- G U-3'<br>3'- C U -5'<br>15 14** | 9.60 (2.07) | -12.68 (1.05) | **-3.08 (2.32)** |
| **Step 2<br>5 6<br>5'- U C -3'** | 6.01 (2.46) | -12.29 (1.00) | **-6.28 (2.66)** |

| | | | |
|---|---|---|---|
| 3'- U G -5'<br>14 13 | | | |
| **Base pairing energies** | | | |
| **4 15**<br>**G - C** | -26.24 (2.75) | 0.54 (1.76) | **-25.7 (3.26)** |
| **5 14**<br>**U - U** | -4.09 (2.18) | -0.99 (0.84) | **-5.08 (2.34)** |
| **6 13**<br>**C - G** | -26.05 (2.69) | 0.56 (1.76) | **-25.49 (3.21)** |

**Table S19.** Interaction energies of the base-pair steps containing the Ψ-U mismatch for duplex-GΨC [kcal/mol]. (A) The values obtained in this study from the last 400 ns of MD simulations. (B) Those obtained from the most populated cluster in the last 400 ns of MD simulations and (C) Those obtained from the second most populated cluster in the last 400 ns of MD simulations.

| **(A)** | **Coulomb (kcal/mol)** | **VDW (kcal/mol)** | **Total (kcal/mol)** |
|---|---|---|---|
| **Stacking energies** | | | |
| **Step 1**<br><br>**4 5**<br>**5'- G Ψ -3'**<br>**3'- C U -5'**<br>**15 14** | 6.78 (2.80) | -12.43 (1.04) | **-5.65 (2.98)** |
| **Step 2**<br>**5 6**<br>**5'- Ψ C -3'**<br>**3'- U G -5'**<br>**14 13** | 6.34 (2.62) | -12.48 (1.02) | **-6.14 (2.81)** |
| **Base pairing energies** | | | |
| **4 15**<br>**G - C** | -26.38 (2.65) | 0.58 (1.76) | **-25.8 (3.18)** |
| **5 14**<br>**Ψ - U** | -4.61 (2.59) | -1.05 (0.89) | **-5.66 (2.74)** |
| **6 13**<br>**C - G** | -26.20 (2.72) | 0.61 (1.76) | **-25.59 (3.24)** |

| **(B)** | **Coulomb (kcal/mol)** | **VDW (kcal/mol)** | **Total (kcal/mol)** |
|---|---|---|---|
| **Stacking energies** | | | |
| **Step 1**<br>**4 5**<br>**5'- G Ψ -3'**<br>**3'- C U -5'**<br>**15 14** | 7.71 (2.32) | -12.57 (1.01) | **-4.86 (2.53)** |
| **Step 2**<br>**5 6**<br>**5'- Ψ C -3'**<br>**3'- U G -5'**<br>**14 13** | 5.42 (2.25) | -12.37 (0.95) | **-6.95 (2.44)** |
| **Base pairing energies** | | | |
| **4 15**<br>**G - C** | -26.46 (2.62) | 0.52 (1.75) | **-25.94 (3.15)** |
| **5 14**<br>**Ψ - U** | -4.91 (2.26) | -1.03 (0.96) | **-5.94 (2.46)** |
| **6 13**<br>**C - G** | -26.18 (2.79) | 0.75 (1.77) | **-25.43 (3.30)** |

| **(C)** | **Coulomb (kcal/mol)** | **VDW (kcal/mol)** | **Total (kcal/mol)** |
|---|---|---|---|
| **Stacking energies** | | | |
| **Step 1**<br>**4 5**<br>**5'- G Ψ -3'**<br>**3'- C U -5'**<br>**15 14** | 4.22 (2.39) | -12.22 (0.94) | **-8.0 (2.56)** |
| **Step 2**<br>**5 6**<br>**5'- Ψ C -3'**<br>**3'- U G -5'**<br>**14 13** | 8.72 (2.02) | -12.92 (0.99) | **-4.2 (2.25)** |
| **Base pairing energies** | | | |

| | | | |
|---|---|---|---|
| **4 15<br>G - C** | -26.24 (2.72) | 0.62 (1.77) | **-25.62 (3.24)** |
| **5 14<br>Ψ - U** | -4.75 (2.31) | -1.14 (0.90) | **-5.89 (2.48)** |
| **6 13<br>C - G** | -26.29 (2.51) | 0.54 (1.73) | **-25.75 (3.05)** |

**Table S20.** Interaction energies of the base-pair steps containing the m[1]Ψ-C mismatch for duplex-Gm[1]ΨC [kcal/mol]. (A) The values obtained in this study from the last 400 ns of MD simulations. (B) Those obtained from the most populated cluster in the last 400 ns of MD simulations and (C) Those obtained from the second most populated cluster in the last 400 ns of MD simulations.

| **(A)** | **Coulomb (kcal/mol)** | **VDW (kcal/mol)** | **Total (kcal/mol)** |
|---|---|---|---|
| **Stacking energies** | | | |
| **Step 1**<br>**4 5**<br>**5'- G m[1]Ψ -3'**<br>**3'- C C -5'**<br>**15 14** | -1.89 (1.89)<br>-2.01 (1.89) | -12.57 (1.24)<br>-12.51 (1.49) | **-14.46 (2.26)**<br>**-14.52 (2.41)** |
| **Step 2**<br>**5 6**<br>**5'- m[1]Ψ C -3'**<br>**3'- C G -5'**<br>**14 13** | 2.62 (2.32)<br>2.54 (2.38) | -11.76 (1.28)<br>-11.85 (1.32) | **-9.14 (2.65)**<br>**-9.31 (2.72)** |
| **Base pairing energies** | | | |
| **4 15<br>G - C** | -26.19 (2.63)<br>-26.22 (2.62) | 0.33 (1.69)<br>0.32 (1.70) | **-25.86 (3.12)**<br>**-25.90 (3.12)** |
| **5 14<br>m[1]Ψ - C** | -2.80 (2.79)<br>-3.02 (2.95) | -0.68 (0.70)<br>-0.71 (0.75) | **-3.48 (2.87)**<br>**-3.72 (3.04)** |
| **6 13<br>C - G** | -26.02 (3.01)<br>-25.76 (2.51) | 0.30 (1.71)<br>0.32 (1.69) | **-25.72 (3.46)**<br>**-25.44 (3.03)** |
| The values obtained in this study from the last 400 ns of MD simulations of replicate 1 and replicate 2 are in the top and bottom respectively. | | | |

| (B) | Coulomb (kcal/mol) | VDW (kcal/mol) | Total (kcal/mol) |
|---|---|---|---|
| **Stacking energies** | | | |
| **Step 1**<br><br>**4 5**<br>**5'- G $m^1\Psi$ -3'**<br>**3'- C C -5'**<br>**15 14** | -1.41 (1.58) | -12.54 (1.08) | **-13.95 (1.91)** |
| **Step 2**<br>**5 6**<br>**5'- $m^1\Psi$ C -3'**<br>**3'- C G -5'**<br>**14 13** | 1.18 (1.14) | -11.04 (1.06) | **-9.86 (1.56)** |
| **Base pairing energies** | | | |
| **4 15**<br>**G - C** | -26.15 (2.57) | 0.28 (1.69) | **-25.87 (3.08)** |
| **5 14**<br>**$m^1\Psi$ - C** | -0.55 (1.14) | -0.79 (0.52) | **-1.34 (1.25)** |
| **6 13**<br>**C - G** | -26.44 (2.55) | 0.48 (1.76) | **-25.96 (3.09)** |
| The values obtained in this study from the most populated cluster in the last 400 ns of MD simulations of replicate 1. | | | |

| (C) | Coulomb (kcal/mol) | VDW (kcal/mol) | Total (kcal/mol) |
|---|---|---|---|
| **Stacking energies** | | | |
| **Step 1**<br><br>**4 5**<br>**5'- G $m^1\Psi$ -3'**<br>**3'- C C -5'**<br>**15 14** | -2.94 (1.36) | -12.48 (1.32) | **-15.48 (1.89)** |
| **Step 2**<br>**5 6**<br>**5'- $m^1\Psi$ C -3'**<br>**3'- C G -5'**<br>**14 13** | 3.85 (1.67) | -12.25 (1.08) | **-8.4 (1.99)** |

| | | | |
|---|---|---|---|
| **Base pairing energies** | | | |
| **4 15**<br>**G - C** | -26.18 (2.57) | 0.30 (1.66) | **-25.88 (3.06)** |
| **5 14**<br>**m$^1$Ψ - C** | -4.56 (1.59) | -0.49 (0.73) | **-5.05 (1.75)** |
| **6 13**<br>**C - G** | -26.04 (2.54) | 0.08 (1.64) | **-26.32 (3.02)** |
| The values obtained in this study from the second most populated cluster in the last 400 ns of MD simulations of replicate 1. | | | |

**Table S21.** Interaction energies of the base-pair steps containing the U-C mismatch for duplex-GUC [kcal/mol]. (A) The values obtained in this study from the last 400 ns of MD simulations. (B) Those obtained from the most populated cluster in the last 400 ns of MD simulations and (C) Those obtained from the second most populated cluster in the last 400 ns of MD simulations.

| **(A)** | **Coulomb (kcal/mol)** | **VDW (kcal/mol)** | **Total (kcal/mol)** |
|---|---|---|---|
| **Stacking energies** | | | |
| **Step 1**<br><br>**4 5**<br>**5'- G U-3'**<br>**3'- C C -5'**<br>**15 14** | 1.81 (3.26) | -10.48 (2.67) | **-8.67 (4.21)** |
| **Step 2**<br>**5 6**<br>**5'- U C -3'**<br>**3'- C G -5'**<br>**14 13** | 3.14 (2.49) | -10.28 (2.25) | **-7.14 (3.36)** |
| **Base pairing energies** | | | |
| **4 15**<br>**G - C** | -25.86 (2.80) | 0.29 (1.72) | **-25.57 (3.28)** |
| **5 14**<br>**U - C** | -2.07 (2.65) | -0.46 (0.77) | **-2.53 (2.75)** |
| **6 13**<br>**C - G** | -26.14 (2.59) | 0.29 (1.70) | **-25.85 (3.09)** |

| (B) | Coulomb (kcal/mol) | VDW (kcal/mol) | Total (kcal/mol) |
|---|---|---|---|
| **Stacking energies** | | | |
| **Step 1**<br><br>**4 5**<br>**5'- G U-3'**<br>**3'- C C -5'**<br>**15 14** | 0.14 (2.29) | -10.22 (1.70) | **-10.08 (2.85)** |
| **Step 2**<br>**5 6**<br>**5'- U C -3'**<br>**3'- C G -5'**<br>**14 13** | 3.95 (2.08) | -10.33 (1.70) | **-6.38 (2.69)** |
| **Base pairing energies** | | | |
| **4 15**<br>**G - C** | -25.66 (2.91) | 0.24 (1.70) | **-25.42 (3.37)** |
| **5 14**<br>**U - C** | -3.79 (2.17) | -0.23 (0.87) | **-4.02 (2.34)** |
| **6 13**<br>**C - G** | -26.02 (2.58) | 0.16 (1.66) | **-25.86 (3.07)** |

| (C) | Coulomb (kcal/mol) | VDW (kcal/mol) | Total (kcal/mol) |
|---|---|---|---|
| **Stacking energies** | | | |
| **Step 1**<br><br>**4 5**<br>**5'- G U-3'**<br>**3'- C C -5'**<br>**15 14** | 4.24 (2.11) | -11.92 (1.08) | **-6.96 (2.37)** |
| **Step 2**<br>**5 6**<br>**5'- U C -3'**<br>**3'- C G -5'**<br>**14 13** | 2.62 (2.35) | -11.11 (1.14) | **-8.49 (2.61)** |
| **Base pairing energies** | | | |

| | | | |
|---|---|---|---|
| **4 15<br>G - C** | -25.98 (2.69) | 0.29 (1.72) | **-25.69 (3.19)** |
| **5 14<br>U - C** | -0.50 (1.95) | -0.81 (0.52) | **-1.31 (2.02)** |
| **6 13<br>C - G** | -26.26 (2.59) | 0.44 (1.74) | **-25.82 (3.12)** |

**Table S22.** Interaction energies of the base-pair steps containing the Ψ-C mismatch for duplex-GΨC [kcal/mol]. (A) The values obtained in this study from the last 400 ns of MD simulations. (B) The values obtained in this study from the most populated cluster in the last 400 ns of MD simulations and (C) the values obtained in this study from the second most populated cluster in the last 400 ns of MD simulations.

| **(A)** | **Coulomb (kcal/mol)** | **VDW (kcal/mol)** | **Total (kcal/mol)** |
|---|---|---|---|
| **Stacking energies** | | | |
| **Step 1**<br><br>**4 5**<br>**5'- G Ψ -3'**<br>**3'- C C -5'**<br>**15 14** | 2.37 (2.26) | -11.64 (1.13) | **-9.27 (2.53)** |
| **Step 2**<br>**5 6**<br>**5'- Ψ C -3'**<br>**3'- C G -5'**<br>**14 13** | 3.19 (3.11) | -11.34 (1.19) | **-8.15 (3.33)** |
| **Base pairing energies** | | | |
| **4 15<br>G - C** | -26.11 (2.73) | 0.40 (1.72) | **-25.71 (3.23)** |
| **5 14<br>Ψ - C** | -1.63 (2.80) | -0.68 (0.72) | **-2.31 (2.89)** |
| **6 13<br>C - G** | -26.25 (2.60) | 0.39 (1.72) | **-25.86 (3.12)** |

| **(B)** | **Coulomb (kcal/mol)** | **VDW (kcal/mol)** | **Total (kcal/mol)** |
|---|---|---|---|

| | | | |
|---|---|---|---|
| **Stacking energies** | | | |
| **Step 1**<br><br>**4 5**<br>**5'- G Ψ -3'**<br>**3'- C C -5'**<br>**15 14** | 2.66 (1.65) | -11.78 (1.00) | **-9.12 (1.39)** |
| **Step 2**<br>**5 6**<br>**5'- Ψ C -3'**<br>**3'- C G -5'**<br>**14 13** | 1.63 (1.96) | -10.96 (1.04) | **-9.33 (2.22)** |
| **Base pairing energies** | | | |
| **4 15**<br>**G - C** | -26.28 (2.54) | 0.42 (1.73) | **-25.86 (3.07)** |
| **5 14**<br>**Ψ - C** | -0.32 (1.25) | -0.80 (0.53) | **-1.12 (1.36)** |
| **6 13**<br>**C - G** | -26.45 (2.53) | 0.52 (1.73) | **-25.93 (3.06)** |

| **(C)** | **Coulomb (kcal/mol)** | **VDW (kcal/mol)** | **Total (kcal/mol)** |
|---|---|---|---|
| **Stacking energies** | | | |
| **Step 1**<br><br>**4 5**<br>**5'- G Ψ -3'**<br>**3'- C C -5'**<br>**15 14** | 1.53 (2.12) | -11.66 (1.18) | **-10.13 (2.43)** |
| **Step 2**<br>**5 6**<br>**5'- Ψ C -3'**<br>**3'- C G -5'**<br>**14 13** | 4.52 (2.18) | -11.94 (1.12) | **-7.42 (2.45)** |
| **Base pairing energies** | | | |
| **4 15**<br>**G - C** | -25.94 (2.86) | 0.38 (1.71) | **-25.56 (3.33)** |
| **5 14** | -4.43 (1.72) | -0.50 (0.91) | **-4.93 (1.94)** |

| | | | |
|---|---|---|---|
| **Ψ - C** | | | |
| **6 13 C - G** | -26.14 (2.59) | 0.25 (1.68) | **-25.89 (3.09)** |
| [a] The values obtained in this study from the last 400 ns of MD simulations<br>[b] The values obtained in this study from the most populated cluster in the last 400 ns of MD simulations<br>[c] The values obtained in this study from the second most populated cluster in the last 400 ns of MD simulations | | | |

**Table S23.** Average and standard deviations of MM-GBSA binding free energies [in kcal/mol] of (A) Duplex-Gm$^1$ΨC, (B) Duplex-Cm$^1$ΨG, (C) Duplex-Um$^1$ΨA and (D) Duplex-Am$^1$ΨU containing the m$^1$Ψ-A pair. $\sigma_M$ is the standard error of mean.

| Energy component | (A) Duplex-Gm$^1$ΨC | | |
|---|---|---|---|
| | Avg | stdev | $\sigma_M$ |
| E_VDW | -48.26<br>-48.22 | 4.72<br>4.72 | 0.075<br>0.075 |
| E_EL | 1056.21<br>1053.84 | 30.98<br>30.33 | 0.49<br>0.48 |
| E_GB | -1089.72<br>-1087.43 | 29.96<br>29.44 | 0.47<br>0.46 |
| E_SURF | -5.01<br>-4.99 | 0.13<br>0.1286 | 0.002<br>0.002 |
| E(EL+GB) | -33.51<br>-33.58 | | |
| ΔGbind | -86.78<br>-86.80 | 3.97<br>3.76 | 0.063<br>0.059 |
| ΔΔG (w.r.t U-A) | -1.06<br>-1.08 | | |
| ΔS | -37.66<br>-38.38 | | |
| The values obtained in this study from the last 400 ns of MD simulations of replicate 1 and replicate 2 are in the top and bottom respectively. | | | |

| Energy component | (B) Duplex-Cm¹ΨG | | |
| --- | --- | --- | --- |
| | Avg | stdev | σM |
| E_VDW | -48.59<br>-48.59 | 4.63<br>4.73 | 0.073<br>0.075 |
| E_EL | 1049.65<br>1048.71 | 29.47<br>28.98 | 0.47<br>0.46 |
| E_GB | -1082.44<br>-1081.78 | 28.42<br>27.98 | 0.45<br>0.44 |
| E_SURF | -4.99<br>-4.99 | 0.15<br>0.14 | 0.002<br>0.002 |
| E(EL+GB) | -32.79<br>-33.07 | | |
| ΔGbind | -86.37<br>-86.65 | 3.87<br>3.70 | 0.061<br>0.059 |
| ΔS | -37.37<br>-37.57 | | |
| The values obtained in this study from the last 400 ns of MD simulations of replicate 1 and replicate 2 are in the top and bottom respectively. | | | |

| Energy component | (C) Duplex-Um¹ΨA | | |
| --- | --- | --- | --- |
| | Avg | stdev | σM |
| E_VDW | -47.69<br>-47.59 | 4.44<br>4.48 | 0.070<br>0.071 |
| E_EL | 1097.46<br>1098.91 | 29.0665<br>29.9767 | 0.46<br>0.47 |
| E_GB | -1120.87<br>-1122.39 | 28.31<br>29.13 | 0.45<br>0.46 |
| E_SURF | -4.94<br>-4.94 | 0.14<br>0.13 | 0.002<br>0.002 |

| | | | |
|---|---|---|---|
| E(EL+GB) | -23.41<br>-23.48 | | |
| ΔGbind | -76.04<br>-76.02 | 3.73<br>3.68 | 0.059<br>0.058 |
| ΔS | -36.54<br>-36.64 | | |
| The values obtained in this study from the last 400 ns of MD simulations of replicate 1 and replicate 2 are in the top and bottom respectively. | | | |

| Energy component | (D) Duplex-Am$^1$ΨU | | |
|---|---|---|---|
| | Avg | stdev | σM |
| E_VDW | -47.45<br>-47.59 | 4.53<br>4.41 | 0.07<br>0.07 |
| E_EL | 1091.70<br>1094.92 | 28.81<br>28.55 | 0.46<br>0.45 |
| E_GB | -1115.21<br>-1118.37 | 28.29<br>27.81 | 0.45<br>0.44 |
| E_SURF | -4.95<br>-4.94 | 0.13<br>0.12 | 0.002<br>0.002 |
| E(EL+GB) | -23.51<br>-23.44 | | |
| ΔGbind | -75.90<br>-75.98 | 3.81<br>3.57 | 0.06<br>0.06 |
| ΔS | -36.25<br>-36.70 | | |
| The values obtained in this study from the last 400 ns of MD simulations of replicate 1 and replicate 2 are in the top and bottom respectively. | | | |

**Table S24.** Average and standard deviations of MM-GBSA binding free energies [in kcal/mol] of (A) Duplex-GUC (U-A) and (B) Duplex-GΨC (Ψ-A) respectively. $\sigma_M$ is the standard error of mean.

| Energy component | (A) Duplex-GUC (U-A) | | |
|---|---|---|---|
| | Avg | stdev | σM |
| E_VDW | -47.98 | 4.61 | 0.073 |
| E_EL | 1083.08 | 33.27 | 0.53 |
| E_GB | -1115.84 | 32.28 | 0.51 |
| E_SURF | -4.98 | 0.15 | 0.002 |
| E(EL+GB) | -32.76 | | |
| ΔGbind | -85.72 | 3.89 | 0.062 |
| ΔS | -37.19 | | |

| Energy component | (B) Duplex-GΨC (Ψ-A) | | |
|---|---|---|---|
| | Avg | stdev | σM |
| E_VDW | -47.96 | 4.63 | 0.073 |
| E_EL | 1081.71 | 33.48 | 0.53 |
| E_GB | -1114.69\ | 32.27 | 0.51 |
| E_SURF | -4.98 | 0.14 | 0.002 |
| E(EL+GB) | -32.98 | | |
| ΔGbind | -85.92 | 3.89 | 0.061 |
| ΔΔG (w.r.t U-A) | -0.19 | | |
| ΔS | -37.68 | | |

**Table S25.** Average and standard deviations of MM-GBSA binding free energies [in kcal/mol] of Duplexes (Gm$^1$ΨC context) containing (A) m$^1$Ψ-G (B) m$^1$Ψ-U, (C) m$^1$Ψ-C mismatches respectively. $\sigma_M$ is the standard error of mean.

| Energy component | (A) Duplex-Gm$^1$ΨC (m$^1$Ψ-G) |
|---|---|

| | Avg | stdev | σM |
|---|---|---|---|
| E_VDW | -48.67<br>-48.96 | 4.74<br>4.74 | 0.075<br>0.075 |
| E_EL | 1071.20<br>1067.63 | 35.00<br>33.67 | 0.55<br>0.53 |
| E_GB | -1102.29<br>-1100.66 | 34.18<br>32.74 | 0.54<br>0.52 |
| E_SURF | -5.19<br>-5.06 | 0.17<br>0.13 | 0.003<br>0.002 |
| E(EL+GB) | -31.09<br>-33.03 | | |
| ΔGbind | -84.95<br>-87.06 | 4.17<br>3.78 | 0.066<br>0.059 |
| ΔΔG (w.r.t U-G) | 1.31<br>-0.79 | | |
| ΔS | -37.11<br>-37.43 | | |

The values obtained in this study from the last 400 ns of MD simulations of replicate 1 and replicate 2 are in the top and bottom respectively.

| Energy component | (B) Duplex-G$m^1\Psi$C ($m^1\Psi$-U) | | |
|---|---|---|---|
| | Avg | stdev | σM |
| E_VDW | -47.38<br>-47.25 | 4.85<br>4.85 | 0.077<br>0.076 |
| E_EL | 1090.89<br>1091.55 | 38.29<br>36.73 | 0.61<br>0.58 |
| E_GB | -1121.23<br>-1121.74 | 36.88<br>35.66 | 0.58<br>0.56 |
| E_SURF | -5.05<br>-5.06 | 0.16<br>0.15 | 0.002<br>0.002 |
| E(EL+GB) | -30.34 | | |

| | | | |
|---|---|---|---|
| | -30.19 | | |
| ΔGbind | -82.77<br>-82.50 | 4.51<br>4.56 | 0.071<br>0.072 |
| ΔΔG (w.r.t U-U) | -1.97<br>-1.69 | | |
| ΔS | -36.96<br>-36.98 | | |
| The values obtained in this study from the last 400 ns of MD simulations of replicate 1 and replicate 2 are in the top and bottom respectively. | | | |

| Energy component | (C) Duplex-Gm[1]ΨC (m[1]Ψ-C) | | |
|---|---|---|---|
| | Avg | stdev | σM |
| E_VDW | -45.82<br>-45.83 | 4.71<br>4.83 | 0.074<br>0.076 |
| E_EL | 1087.79<br>1098.58 | 33.30<br>42.66 | 0.53<br>0.67 |
| E_GB | -1116.22<br>-1126.03 | 32.34<br>40.14 | 0.51<br>0.63 |
| E_SURF | -5.00<br>-5.08 | 0.14<br>0.23 | 0.002<br>0.004 |
| E(EL+GB) | -28.42<br>-27.44 | | |
| ΔGbind | -79.24<br>-78.36 | 4.24<br>4.83 | 0.067<br>0.076 |
| ΔΔG (w.r.t U-C) | -3.34<br>-2.46 | | |
| ΔS | -36.46<br>-36.61 | | |
| The values obtained in this study from the last 400 ns of MD simulations of replicate 1 and replicate | | | |

| |
|---|
| 2 are in the top and bottom respectively. |

**Table S26.** Average and standard deviations of MM-GBSA binding free energies [in kcal/mol] of Duplexes (GUC context) containing (A) U-G, (C) U-U (D) U-C mismatches respectively. $\sigma_M$ is the standard error of mean.

| Energy component | (A) Duplex-GUC (U-G) | | |
|---|---|---|---|
| | Avg | stdev | σM |
| E_VDW | -49.06 | 4.69 | 0.074 |
| E_EL | 1101.68 | 39.76 | 0.63 |
| E_GB | -1133.78 | 37.67 | 0.59 |
| E_SURF | -5.09 | 0.24 | 0.004 |
| E(EL+GB) | -32.10 | | |
| ΔGbind | -86.26 | 4.32 | 0.068 |
| ΔS | -36.71 | | |

| Energy component | (B) Duplex-GUC (U-U) | | |
|---|---|---|---|
| | Avg | stdev | σM |
| E_VDW | -47.27 | 4.69 | 0.074 |
| E_EL | 1133.19 | 44.61 | 0.71 |
| E_GB | -1161.66 | 42.41 | 0.67 |
| E_SURF | -5.06 | 0.25 | 0.004 |
| E(EL+GB) | -28.48 | | |
| ΔGbind | -80.80 | 4.91 | 0.078 |
| ΔS | -36.61 | | |

| Energy component | (C) Duplex-GUC (U-C) | | |
|---|---|---|---|
| | Avg | stdev | σM |
| E_VDW | -47.39 | 5.48 | 0.087 |
| E_EL | 1176.96 | 91.58 | 1.45 |
| E_GB | -1200.21 | 85.64 | 1.35 |
| E_SURF | -5.26 | 0.48 | 0.007 |
| E(EL+GB) | -23.25 | | |
| ΔGbind | -75.90 | 5.5206 | 0.087 |
| ΔS | -37.84 | | |

**Table S27.** Average and standard deviations of MM-GBSA binding free energies [in kcal/mol] of Duplexes (GΨC context) containing (A) Ψ-G (B) Ψ-U, (C) Ψ-C mismatches respectively. $\sigma_M$ is the standard error of mean.

| Energy component | (A) Duplex-GΨC (Ψ-G) | | |
|---|---|---|---|
| | Avg | stdev | σM |
| E_VDW | -48.92 | 4.57 | 0.07 |
| E_EL | 1096.54 | 0.61 | 0.61 |
| E_GB | -1129.29 | 36.81 | 0.58 |
| E_SURF | -5.04 | 0.17 | 0.003 |
| E(EL+GB) | -32.75 | | |
| ΔGbind | -86.71 | 3.96 | 0.063 |
| ΔΔG (w.r.t U-G) | -0.45 | | |
| ΔS | -36.86 | | |

| Energy component | (B) Duplex-GΨC (Ψ-U) |
|---|---|

| | Avg | stdev | σM |
|---|---|---|---|
| E_VDW | -47.19 | 4.61 | 0.073 |
| E_EL | 1131.75 | 41.63 | 0.66 |
| E_GB | -1160.55 | 39.49 | 0.62 |
| E_SURF | -5.08 | 0.19 | 0.003 |
| E(EL+GB) | -28.80 | | |
| ΔGbind | -81.07 | 4.64 | 0.073 |
| ΔΔG (w.r.t U-U) | -0.27 | | |
| ΔS | -35.99 | | |

| Energy component | (C) Duplex-GΨC (Ψ-C) | | |
|---|---|---|---|
| | Avg | stdev | σM |
| E_VDW | -46.02 | 4.59 | 0.073 |
| E_EL | 1124.56 | 42.74 | 0.68 |
| E_GB | -1151.49 | 40.78 | 0.64 |
| E_SURF | -5.03 | 0.23 | 0.004 |
| E(EL+GB) | -26.93 | | |
| ΔGbind | -77.98 | 4.53 | 0.072 |
| ΔΔG (w.r.t U-C) | -2.08 | | |
| ΔS | -36.67 | | |

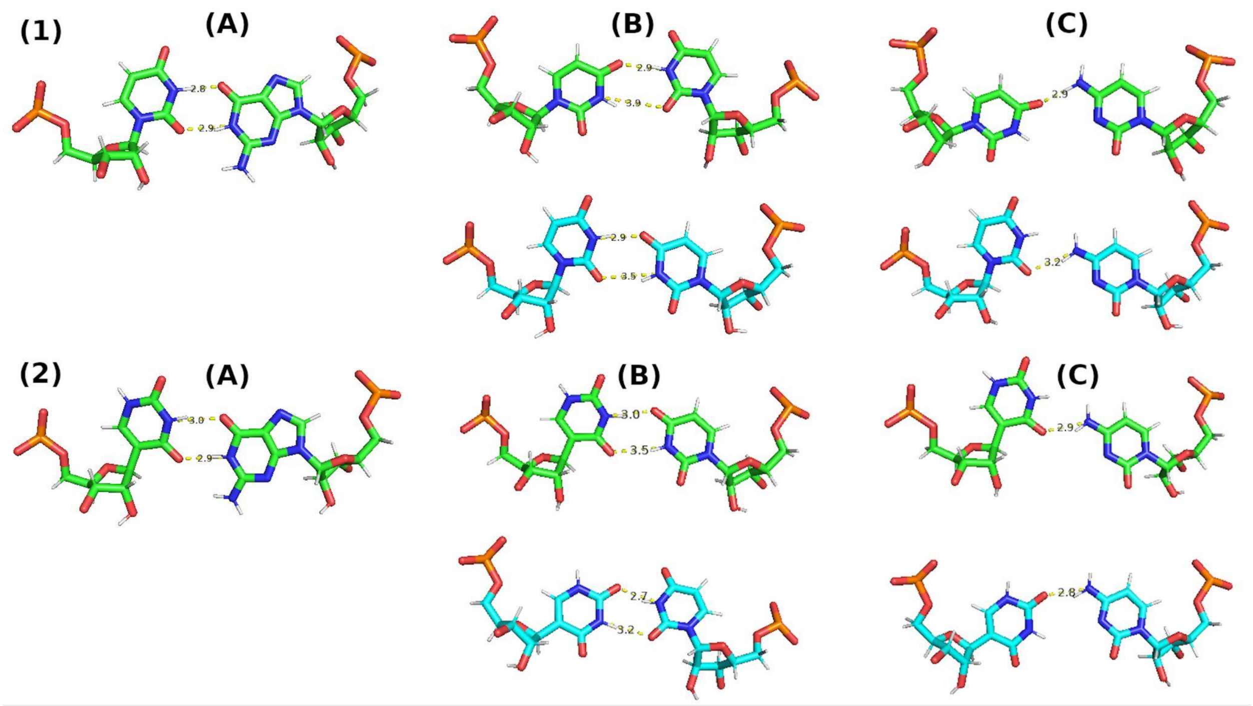

**Figure S1.** (1) Observed geometries of the (A) U-G, (B) U-U and (C) U-C mismatches respectively within Duplex-GUC. (2) Observed geometries of the (A) Ψ-G, (B) Ψ-U and (C) Ψ-C mismatches respectively within Duplex-GΨC. For U-G/Ψ-G mismatch, the geometry corresponds to the centroid structure obtained from the most populated cluster. For the U-U/Ψ-U and U-C/Ψ-C mismatches, the geometries corresponding to the centroid structures obtained from the most populated cluster (top) and the second most populated cluster (bottom) are shown.

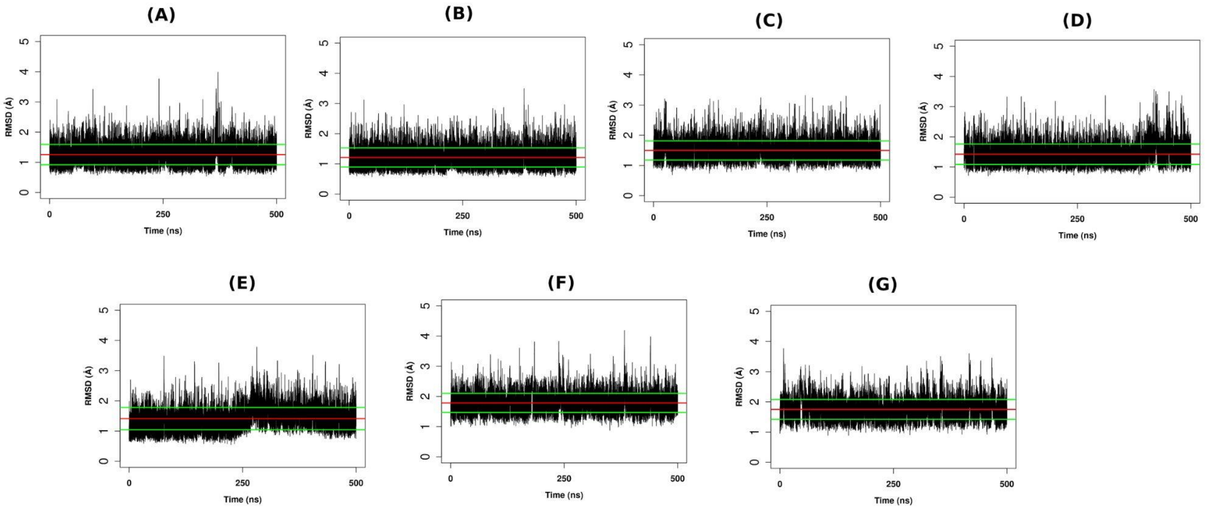

**Figure S2.** Time evolution plots of the root-mean-square deviations (RMSDs) (heavy atoms) of the duplexes (A) Duplex-Gm$^{1}$ΨC (m$^{1}$Ψ-A), (B) Duplex-Cm$^{1}$ΨG (m$^{1}$Ψ-A), (C) Duplex-Um$^{1}$ΨA (m$^{1}$Ψ-A), Duplex-Am$^{1}$ΨU (m$^{1}$Ψ-A), (E) Duplex-Gm$^{1}$ΨC (m$^{1}$Ψ-G), (F) Duplex-Gm$^{1}$ΨC (m$^{1}$Ψ-U), (G) Duplex-Gm$^{1}$ΨC (m$^{1}$Ψ-C) respectively for the first set of 500 ns simulations. The average value of the RMSD is shown as a red line and the average±standard deviation values are shown as green lines.

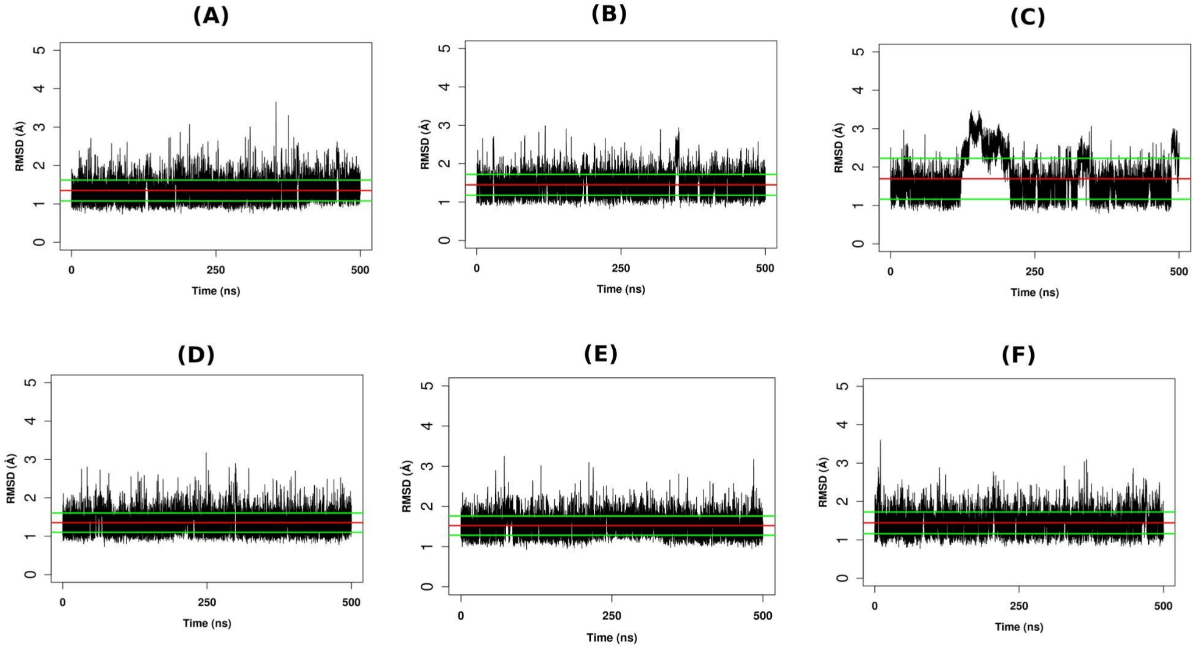

**Figure S3.** Time evolution plots of the root-mean-square deviations (RMSDs) (heavy atoms) of the duplexes (A) Duplex-GUC (U-G), (B) Duplex-GUC (U-U), (C) Duplex-GUC (U-C), Duplex-GΨC (Ψ-G), (E) Duplex-GΨC (Ψ-U), (F) Duplex-GΨC (Ψ-C) respectively. The average value of the RMSD is shown as a red line and the average±standard deviation values are shown as green lines.

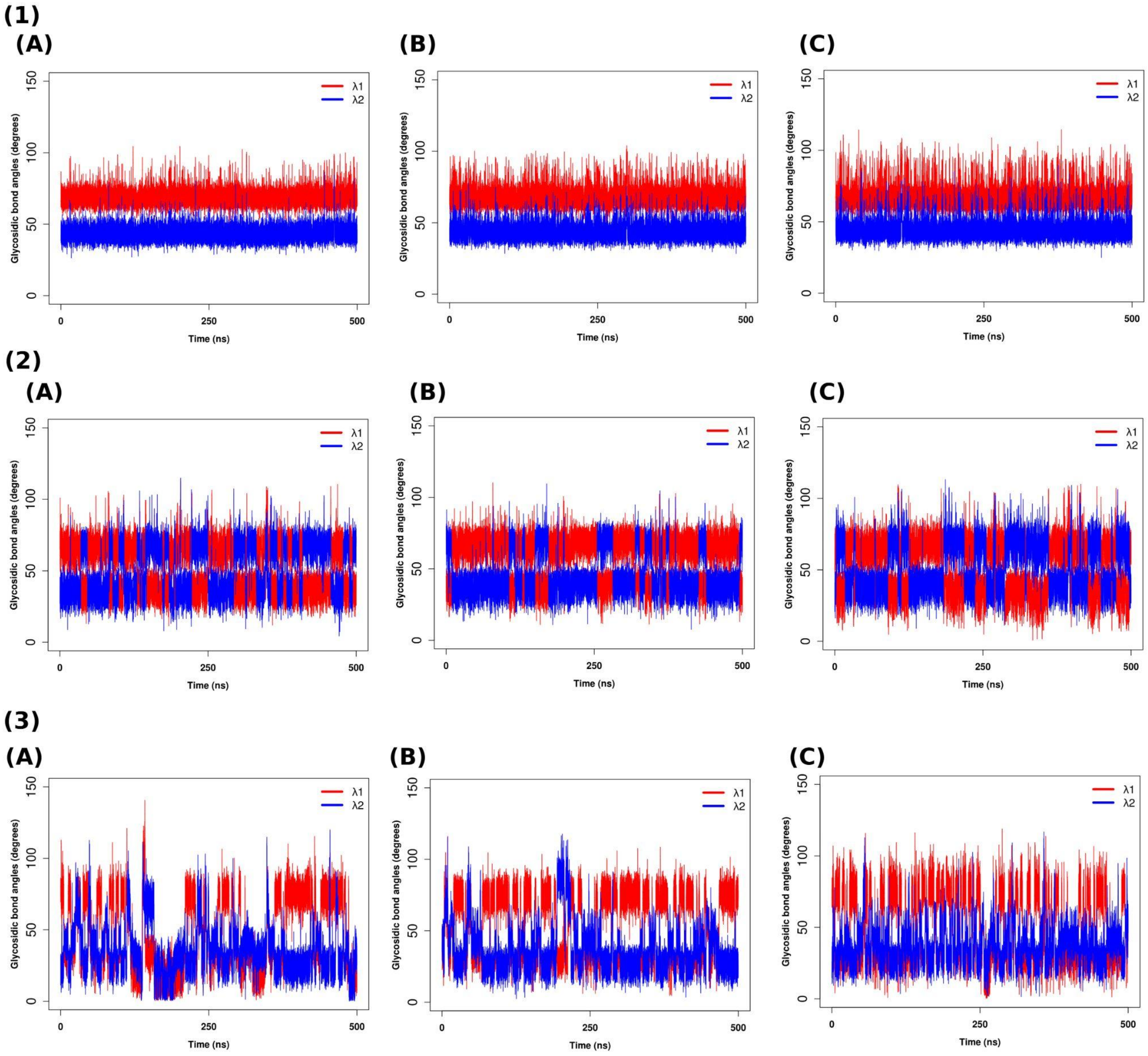

**Figure S4.** Time evolution of λ angles for (1) Y-G, (2) Y-U and (3) Y-C corresponding to (A) Y=U, (B) Y=Ψ and (C) Y=m$^1$Ψ respectively (λ1 and λ2 are indicating the values for the 5$^{th}$ and 14$^{th}$ residues respectively.

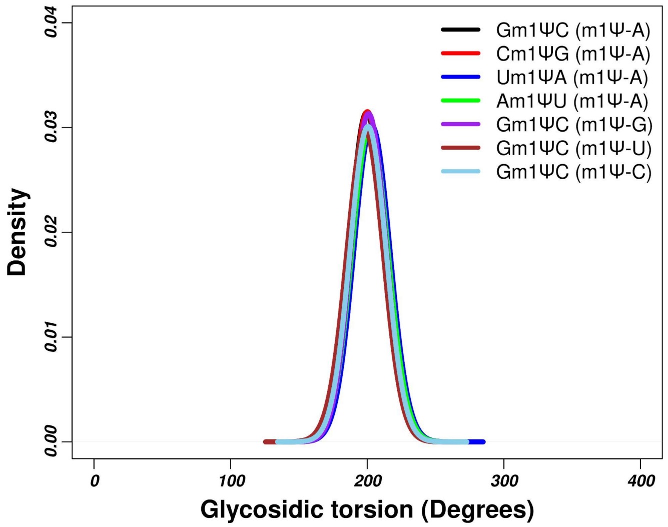

**Figure S5.** Population distribution of glycosidic torsion angle (**χ**) for m[1]Ψ(5) in the simulated duplexes for the last 400ns trajectory (for the first set of simulation).

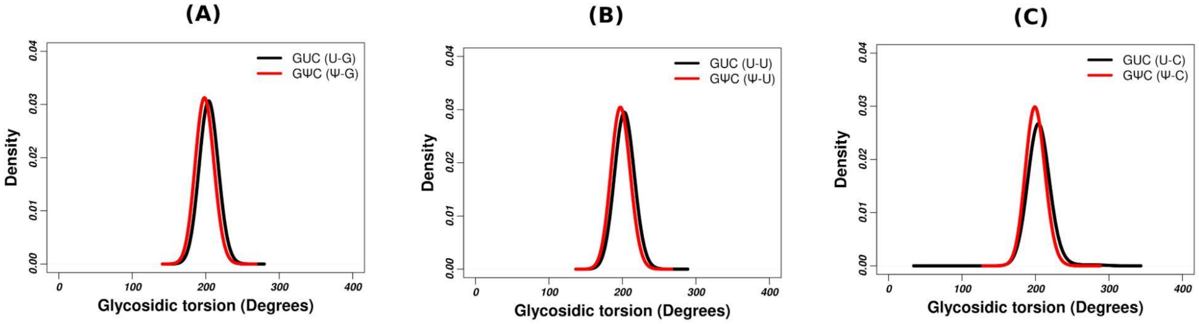

**Figure S6.** Population Distribution of glycosidic torsion angle (**χ**) for U(5) and Ψ(5) in the simulated duplexes for the last 400ns trajectory. (A) For the duplexes Duplex-GUC (U-G) and Duplex-GΨC (Ψ-G); (B) for the duplexes Duplex-GUC (U-U) and Duplex-GΨC (Ψ-U); (C) For the duplexes Duplex-GUC (U-C) and Duplex-GΨC (Ψ-C) respectively.

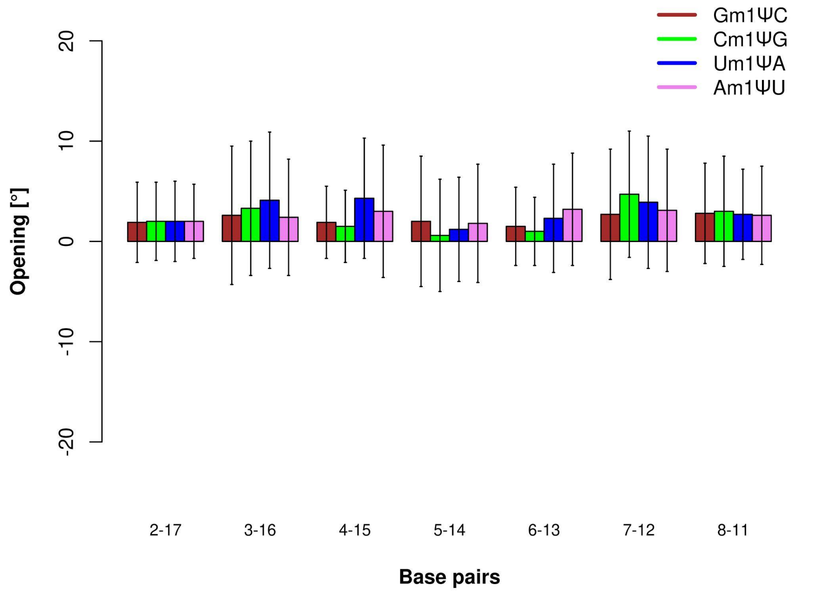

**Figure S7.** Opening [°] of the central 7 base pairs for the duplexes of four different sequence contexts containing $m^1\Psi$-A pair (for the first set of simulation).

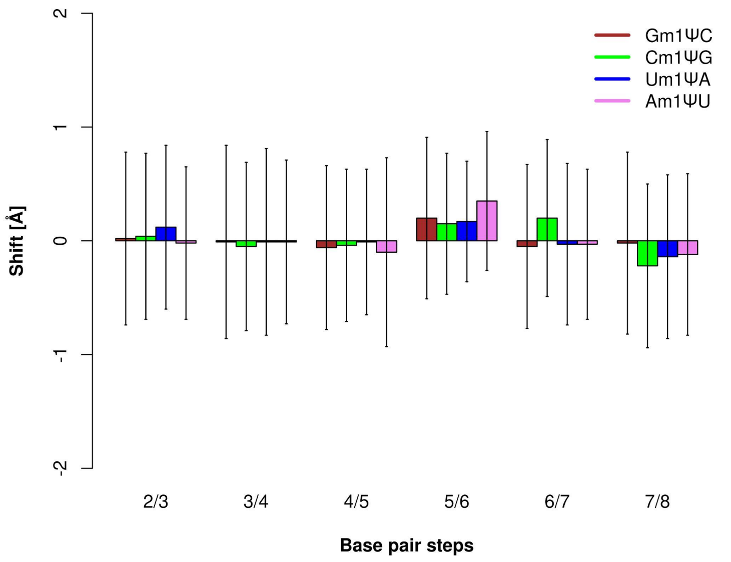

**Figure S8.** Shift [Å] of the central 6 base pair steps for the duplexes of four different sequence contexts containing $m^1\Psi$-A pair (for the first set of simulation).

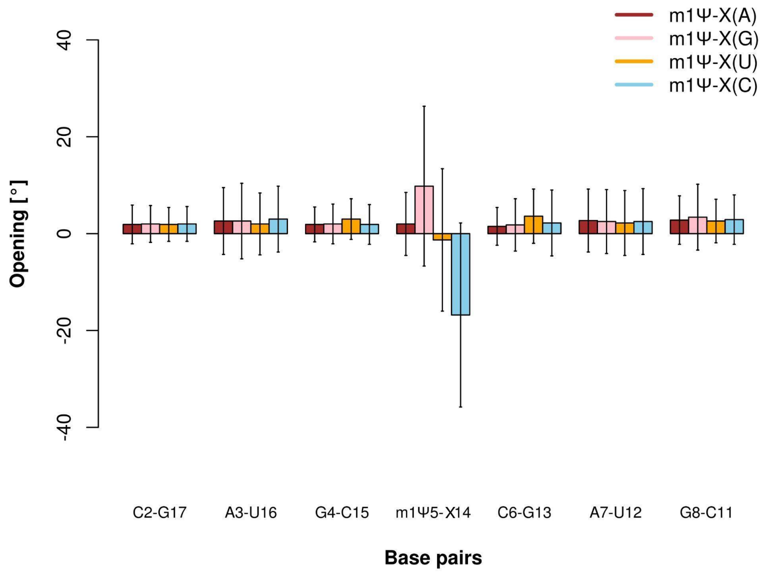

**Figure S9.** Opening [°] of the central 7 base pairs for the duplexes (context Gm[1]ΨC) containing m[1]Ψ-A pair and mismatches m[1]Ψ-G, m[1]Ψ-U and m[1]Ψ-C respectively (for the first set of simulation).

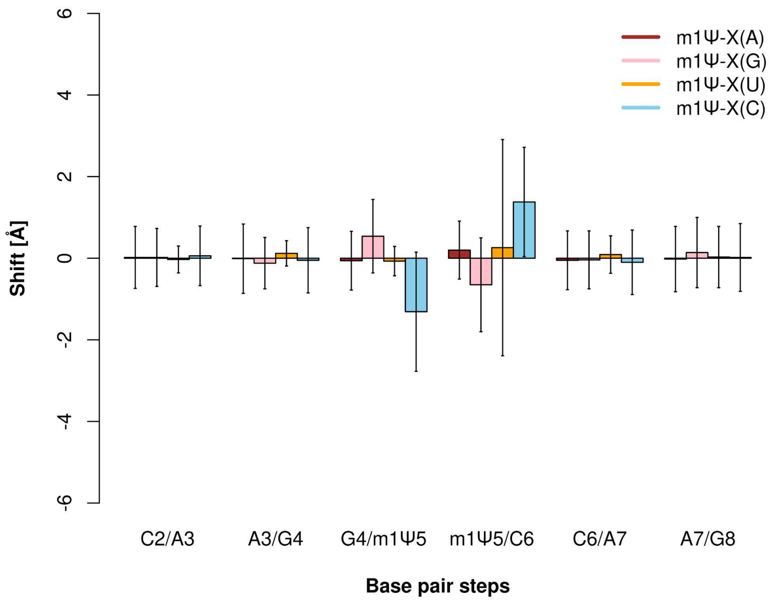

**Figure S10.** Shift [Å] of the central 6 base pair steps for the duplexes (context Gm[1]ΨC) containing m[1]Ψ-A pair and mismatches m[1]Ψ-G, m[1]Ψ-U and m[1]Ψ-C respectively (for the first set of simulation).

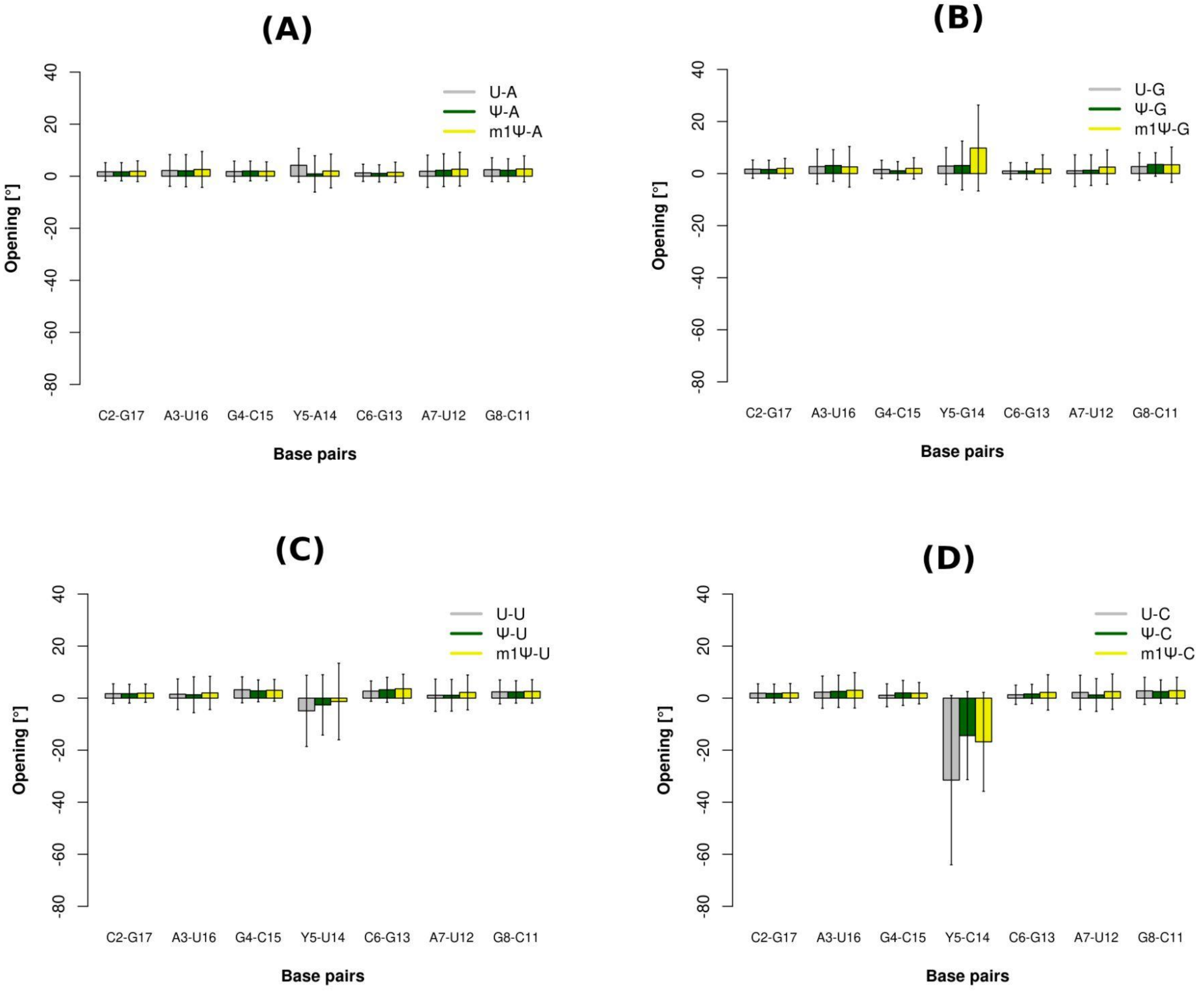

**Figure S11.** Opening [°] of the central 7 base pairs for the duplexes (context GUC/GΨC/Gm$^1$ΨC) (A) containing U-A, Ψ-A and m$^1$Ψ-A pairs; (B) containing U-G, Ψ-G and m$^1$Ψ-G; (C) containing U-U, Ψ-U and m$^1$Ψ-U mismatches; (D) containing U-C, Ψ-C and m$^1$Ψ-C mismatches respectively (values corresponding to the first set of simulation for m$^1$Ψ-modified duplexes are plotted). Y5 in the X axis represents U5/Ψ5/m$^1$Ψ5.

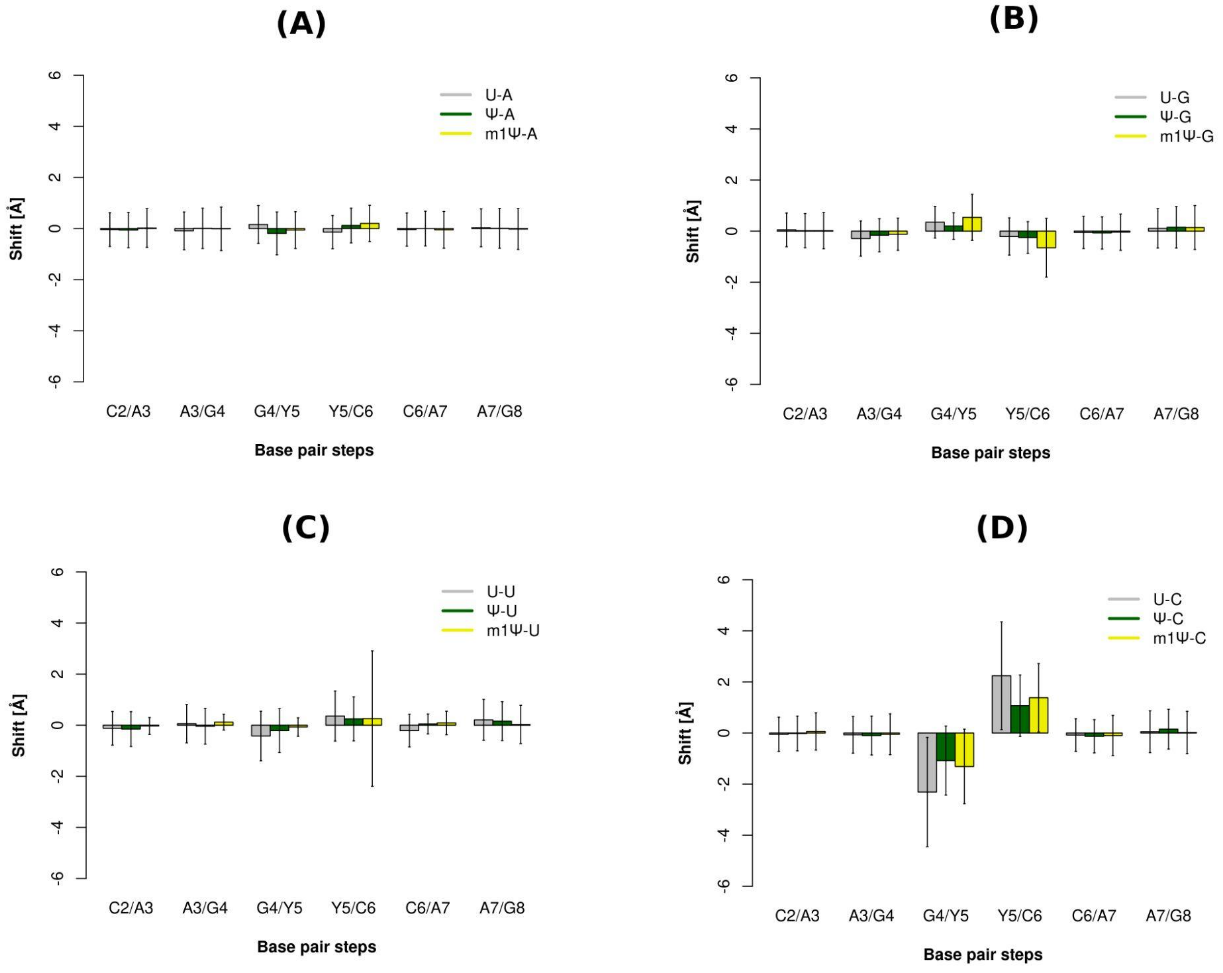

**Figure S12.** Shift [Å] of the central 6 base pair steps for the duplexes (context GUC/GΨC/Gm$^{1}$ΨC) (A) containing U-A, Ψ-A and m$^{1}$Ψ-A pairs; (B) containing U-G, Ψ-G and m$^{1}$Ψ-G; (C) containing U-U, Ψ-U and m$^{1}$Ψ-U mismatches; (D) containing U-C, Ψ-C and m$^{1}$Ψ-C mismatches respectively (values corresponding to the first set of simulation for m$^{1}$Ψ-modified duplexes are plotted). Y5 in the X axis represents U5/Ψ5/m$^{1}$Ψ5.

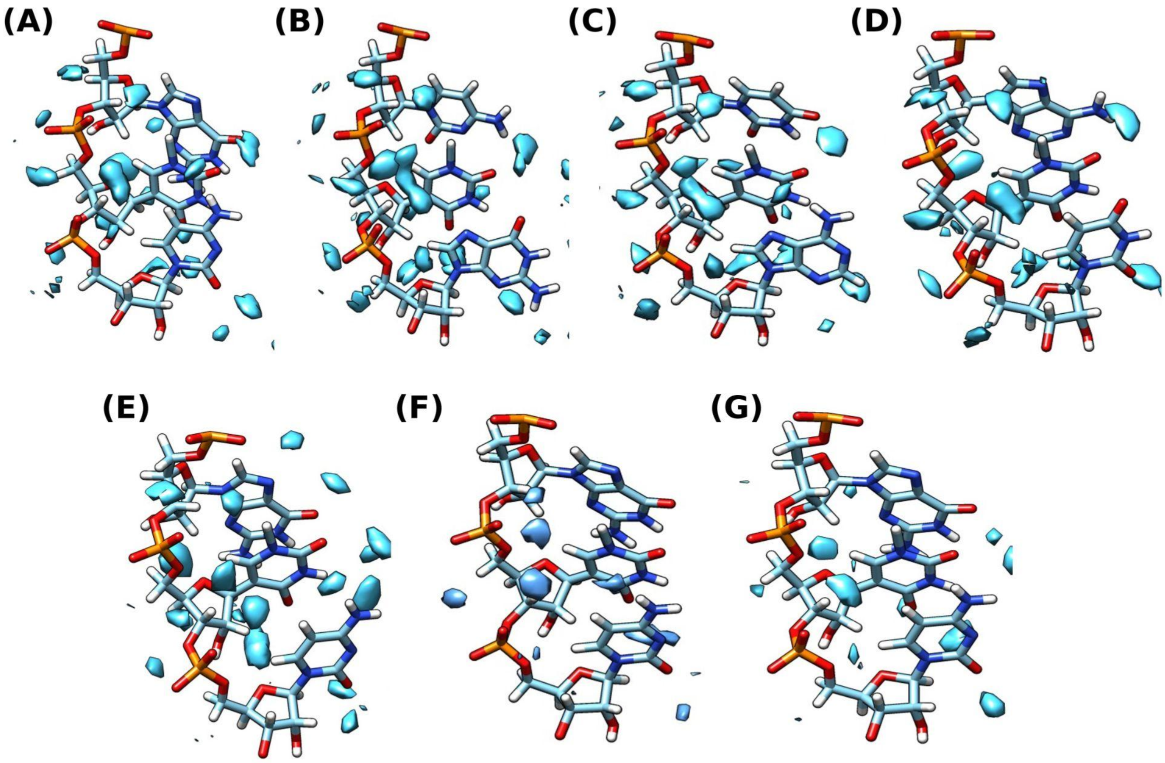

**Figure S13.** Water occupancy maps for the duplexes (residues 4-6) (A) Duplex-Gm$^1$ΨC (m$^1$Ψ-A), (B) Duplex-Cm$^1$ΨG (m$^1$Ψ-A) and (C) Duplex-Um$^1$ΨA (m$^1$Ψ-A), (D) Duplex-Am$^1$ΨU (m$^1$Ψ-A), (E) Duplex-Gm$^1$ΨC (m$^1$Ψ-G), (F) Duplex-Gm$^1$ΨC (m$^1$Ψ-U) and (G) Duplex-Gm$^1$ΨC (m$^1$Ψ-C) respectively (for the first set of simulation).

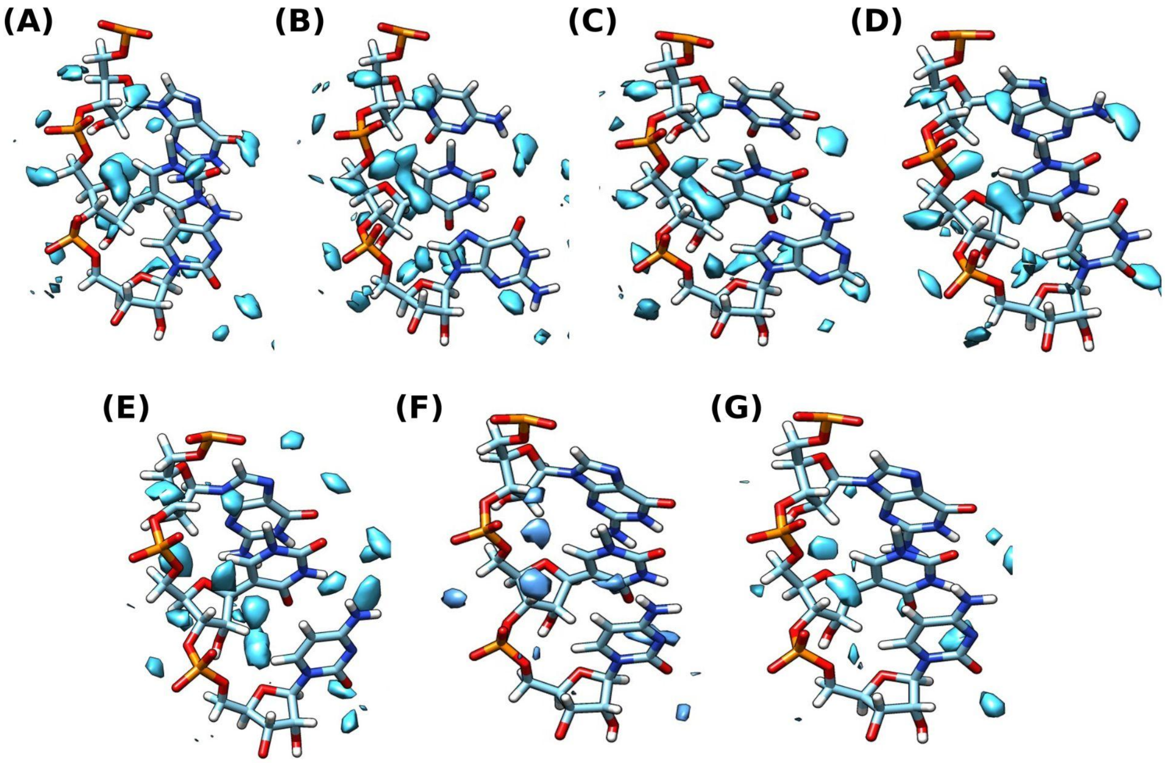

**Figure S14.** Water occupancy maps for the duplexes (residues 4-6) (A) Duplex-GUC (U-G), (B) Duplex-GUC (U-U), (C) Duplex-GUC (U-C), Duplex-GΨC (Ψ-G), (E) Duplex-GΨC (Ψ-U), (F) Duplex-GΨC (Ψ-C) respectively.

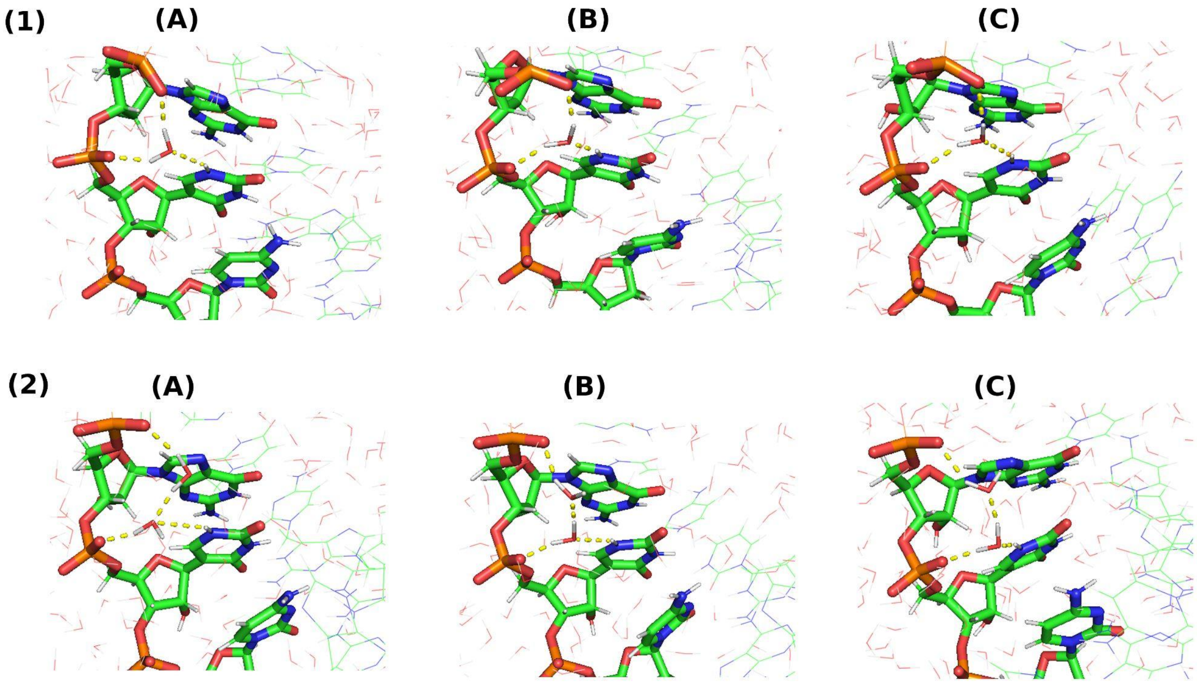

**Figure S15.** Snapshots of water bridges formed between the HN1 and OP2 atoms of Ψ(5) and the OP2 atom of G(4). Panel (1) shows the water bridge interactions involving one water molecule and panel (2) shows the water bridge interactions involving two water molecules for (A) Duplex-GΨC (Ψ-G), (B) Duplex-GΨC (Ψ-U) and (C) Duplex-GΨC (Ψ-C) respectively.

**(1)**

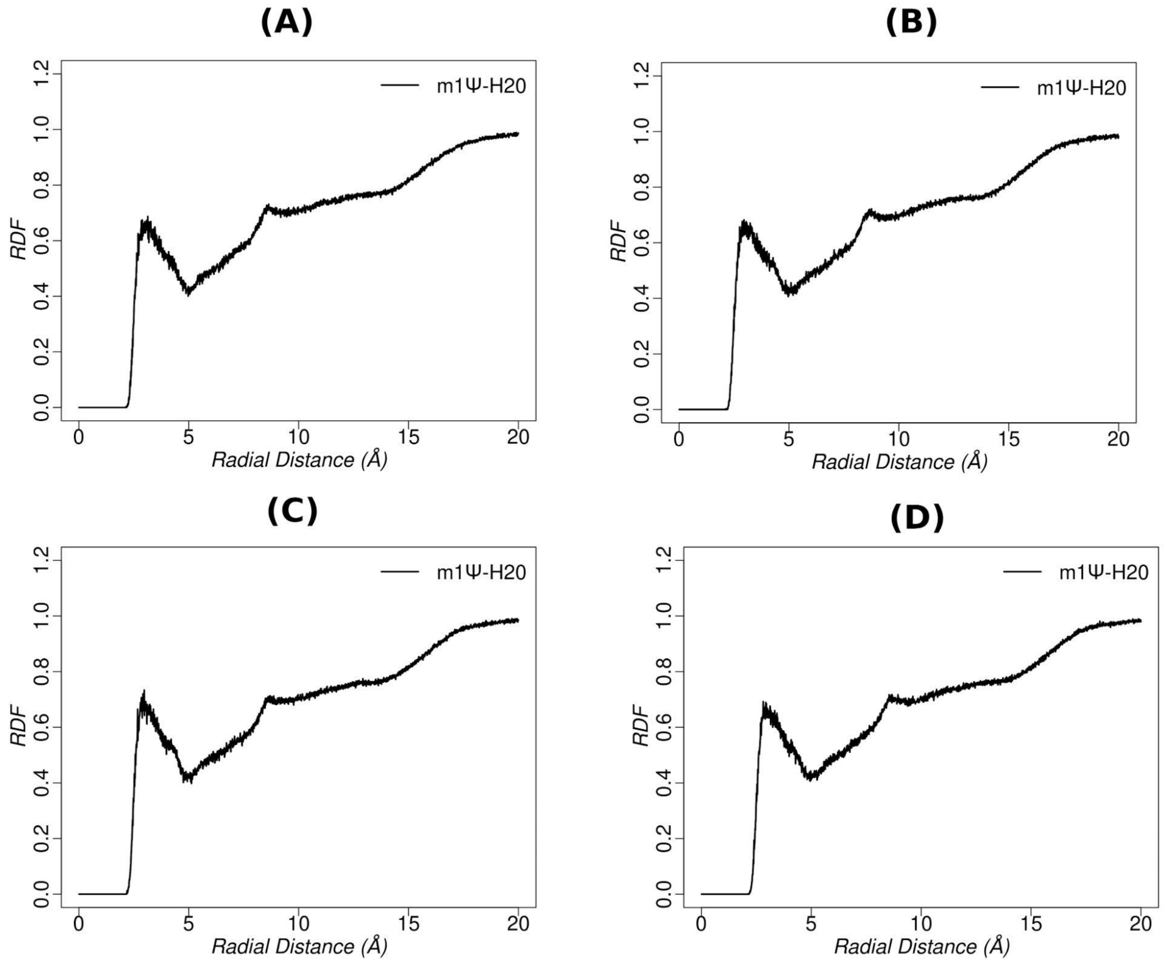
(A)
(B)
(C)
(D)
m1Ψ-H20
RDF
Radial Distance (Å)
0.0
0.2
0.4
0.6
0.8
1.0
1.2
0
5
10
15
20

**(2)**

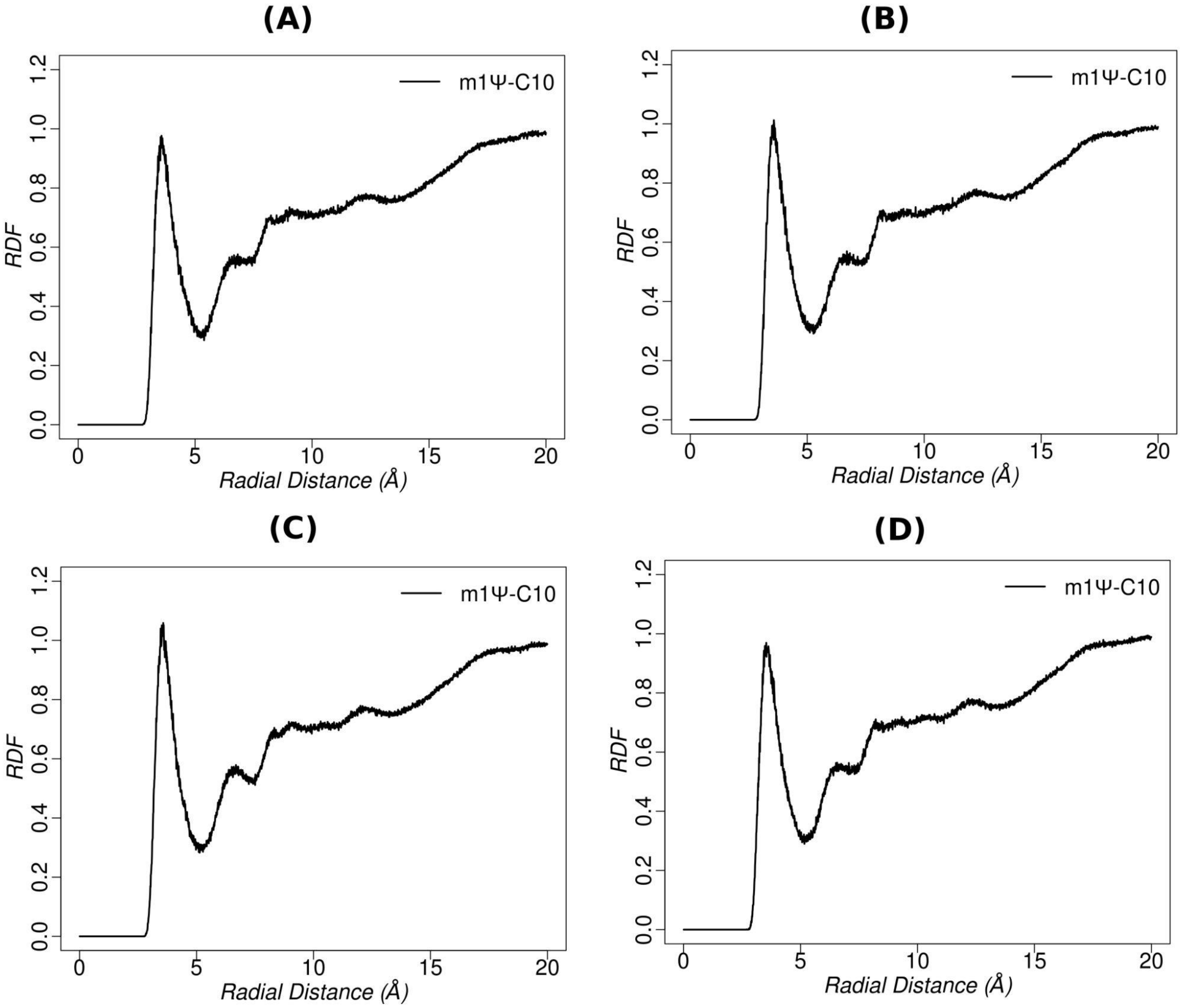

**Figure S16.** RDF of water oxygen atoms around (1) the H20 (methyl hydrogen) atom and (2) the C10 (methyl carbon) atom of $m^1\Psi$(5) for duplexes containing $m^1\Psi$-A pair: (A) Duplex-G$m^1\Psi$C ($m^1\Psi$-A), (B) Duplex-C$m^1\Psi$G ($m^1\Psi$-A) (C) Duplex-U$m^1\Psi$A ($m^1\Psi$-A) and (D) Duplex-A$m^1\Psi$U ($m^1\Psi$-A) (for the first set of simulation).

(1)

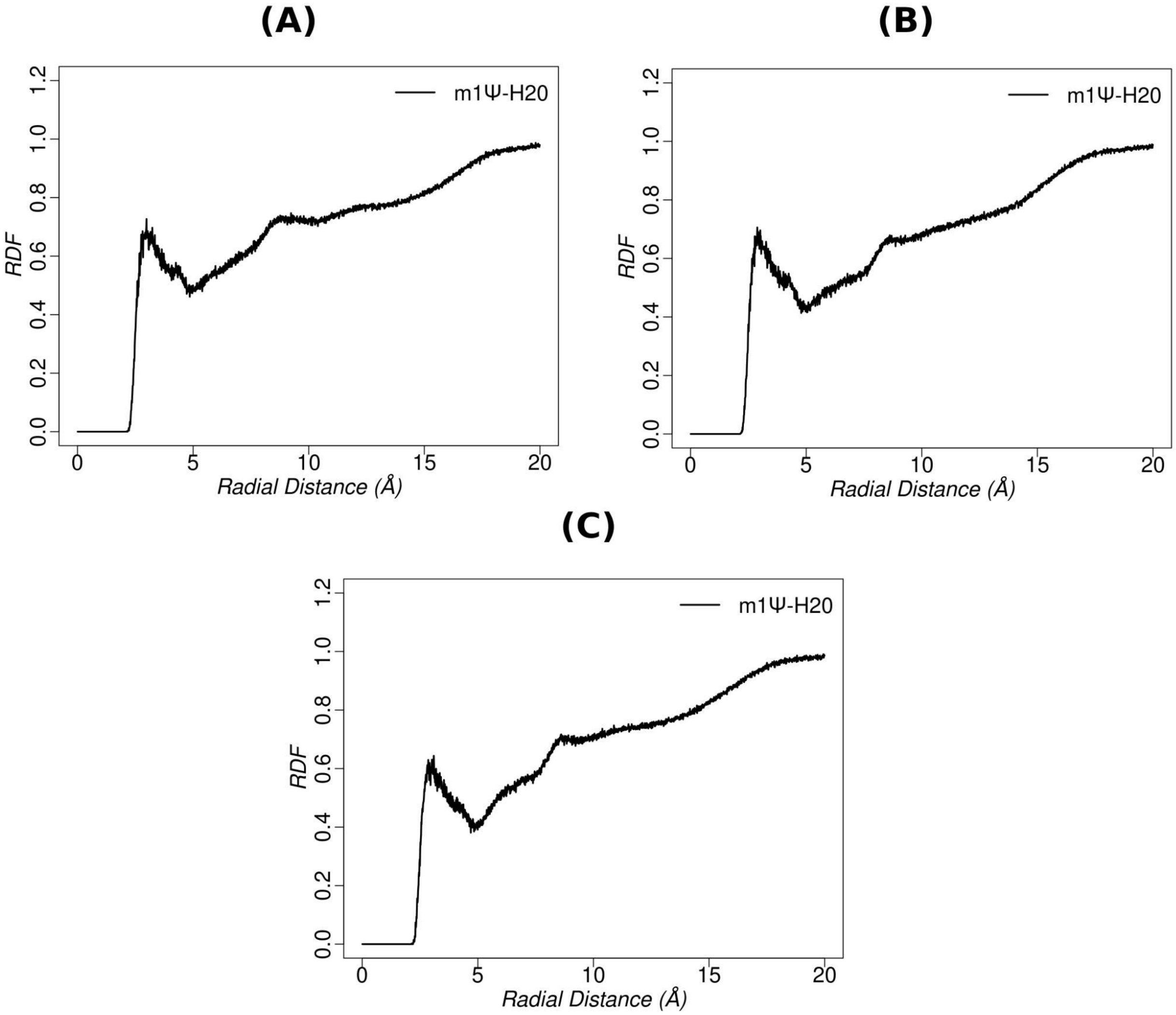

**(2)**

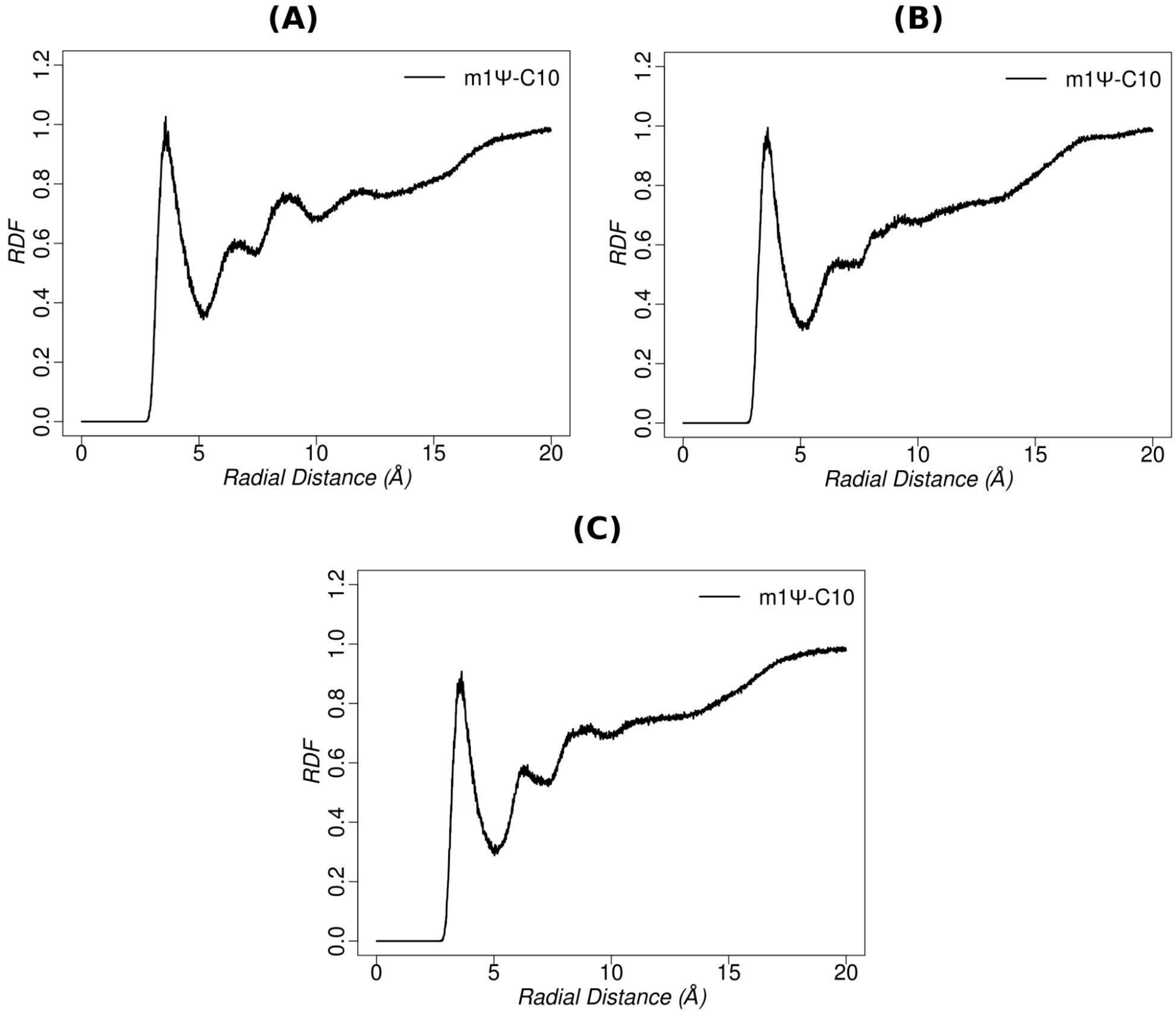

**Figure S17.** RDF of water oxygen atoms around (1) the H20 (methyl hydrogen) atom and (2) the C10 (methyl carbon) atom of m$^{1}$Ψ(5) for duplexes of (Gm$^{1}$ΨC context) containing (A) m$^{1}$Ψ-G, (B) m$^{1}$Ψ-U and (C) m$^{1}$Ψ-C mismatches respectively (for the first set of simulation).

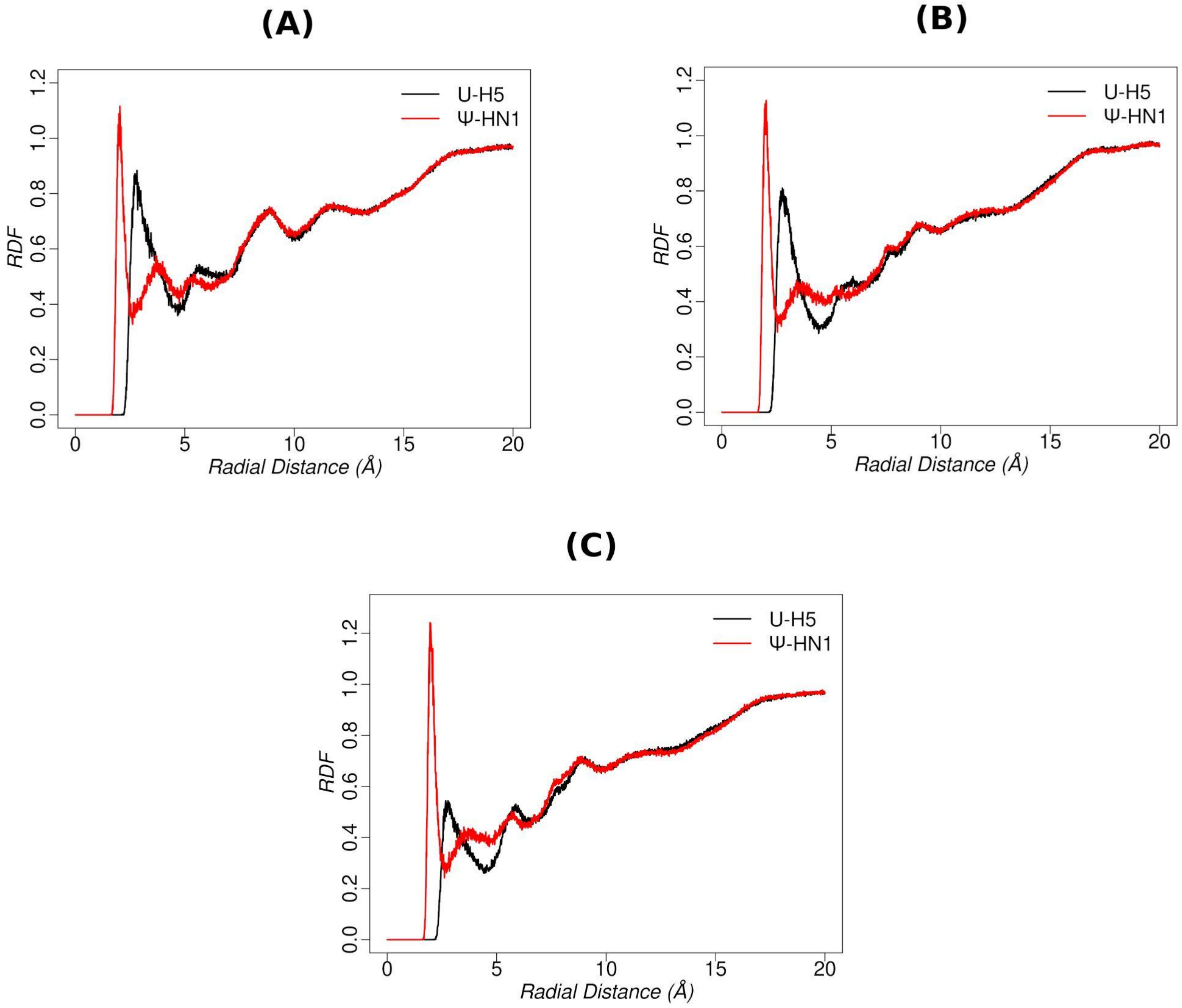

**Figure S18.** RDF of water oxygen atoms around the H5 atom of U(5) and that about the HN1 atom of Ψ(5) for (A) Duplex-GUC(U-G) and Duplex-GΨC(Ψ-G), (B) Duplex-GUC(U-U) and Duplex-GΨC(Ψ-U ) and (C) Duplex-GUC(U-C) and Duplex-GΨC(Ψ-C) respectively.

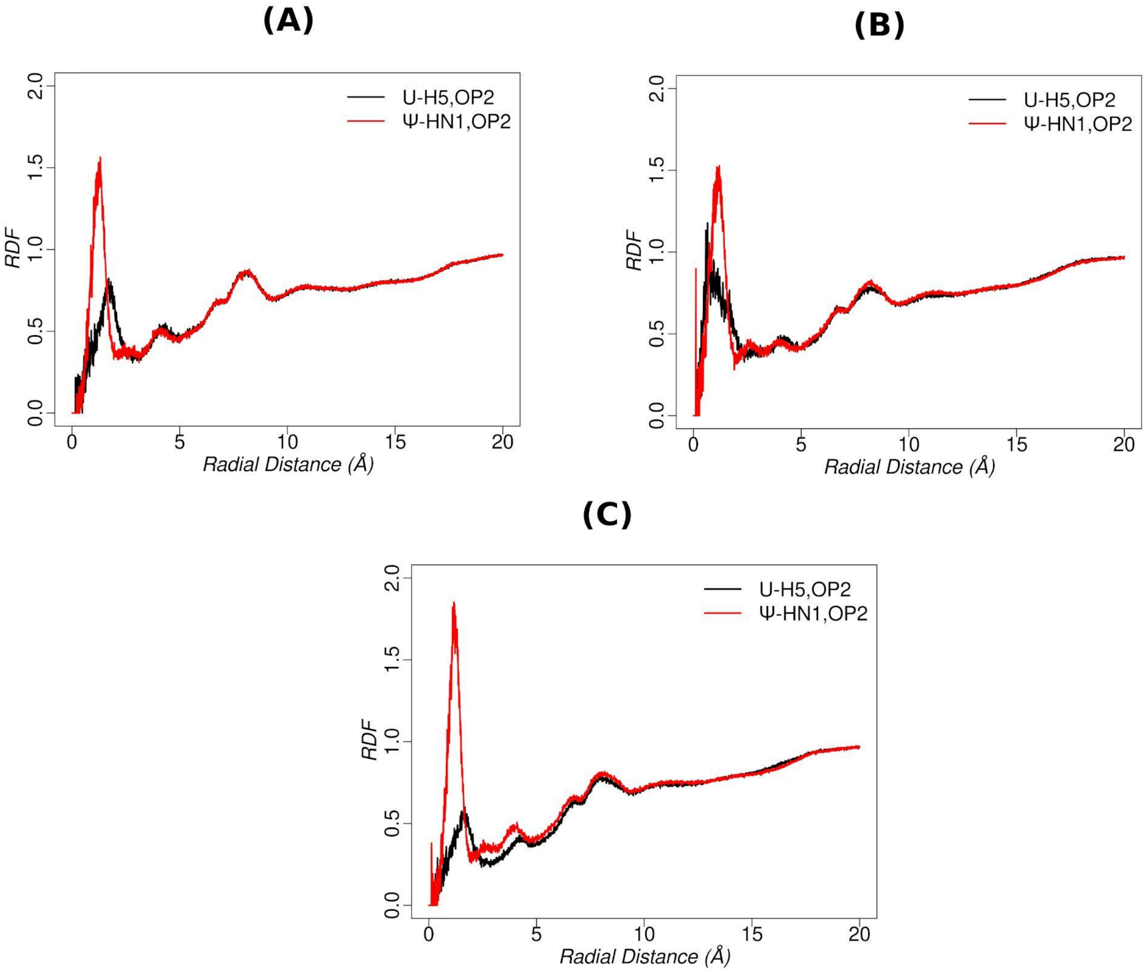

**Figure S19.** RDF of water oxygen atoms around geometric center of the H5 and OP2 atoms of U(5) and that about the geometric center of the HN1 and OP2 atoms of Ψ(5) for (A) Duplex-GUC(U-G) and Duplex-GΨC(Ψ-G), (B) Duplex-GUC(U-U) and Duplex-GΨC(Ψ-U) and (C) Duplex-GUC(U-C) and Duplex-GΨC(Ψ-C) respectively.

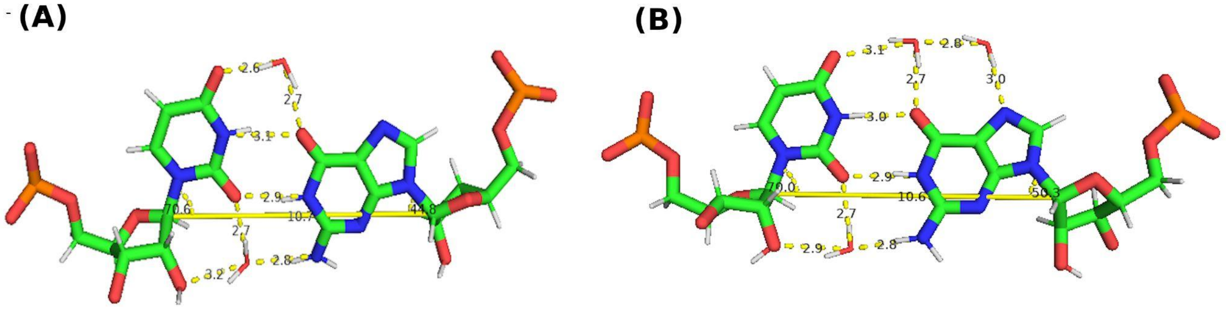

**Figure S20.** Snapshots of U-G wobble base pair (corresponding to Duplex-GUC (U-G)) with water mediated interactions between the bases (A) by two key water molecules (O4(5)-W-O6(14) and O2’-O2(5)-W-N2(14) water bridging interactions) and (B) by three key water molecules ((O4(5)-W-O6(14) and O2’-O2(5)-W-N2(14) water bridging interactions and an water molecule linking the O4(5)-W-O6(14) with N7(14)), other than the two direct hydrogen bonding interactions. Distances between hydrogen bond acceptor and donor atoms are shown with yellow dotted lines. The C1’-C1’ distances and λ ( C1’-C1’-N1 for U and C1’-C1’-N9 for G) angles are also shown.

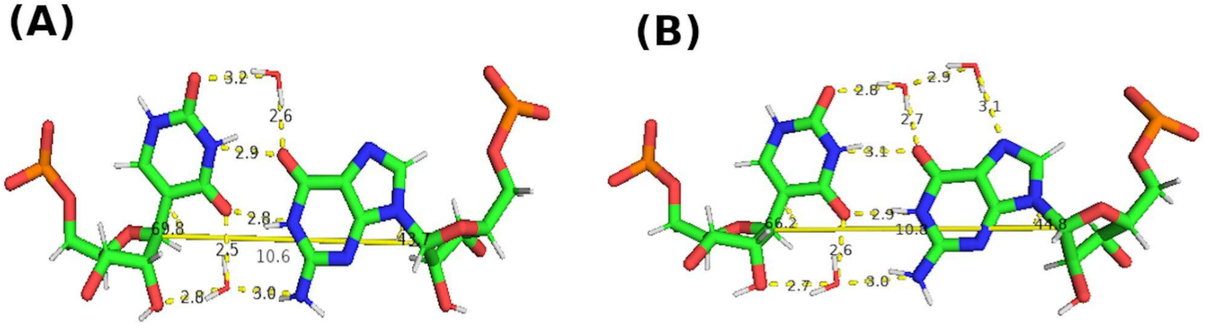

**Figure S21.** Snapshots of Ψ-G wobble base pair (corresponding to Duplex-GΨC (Ψ-G)) with water mediated interactions between the bases (A) by two key water molecules (O2(5)-W-O6(14) and O2’-O4(5)-W-N2(14) water bridging interactions) and (B) by three key water molecules ((O2(5)-W-O6(14) and O2’-O4(5)-W-N2(14) water bridging interactions and an water molecule linking the O2(5)-W-O6(14) with N7(14)), other than the two direct hydrogen bonding interactions. Distances between hydrogen bond acceptor and donor atoms are shown with yellow dotted lines. The C1’-C1’ distances and λ ( C1’-C1’-C5 for Ψ and C1’-C1’-N9 for G) angles are also shown.

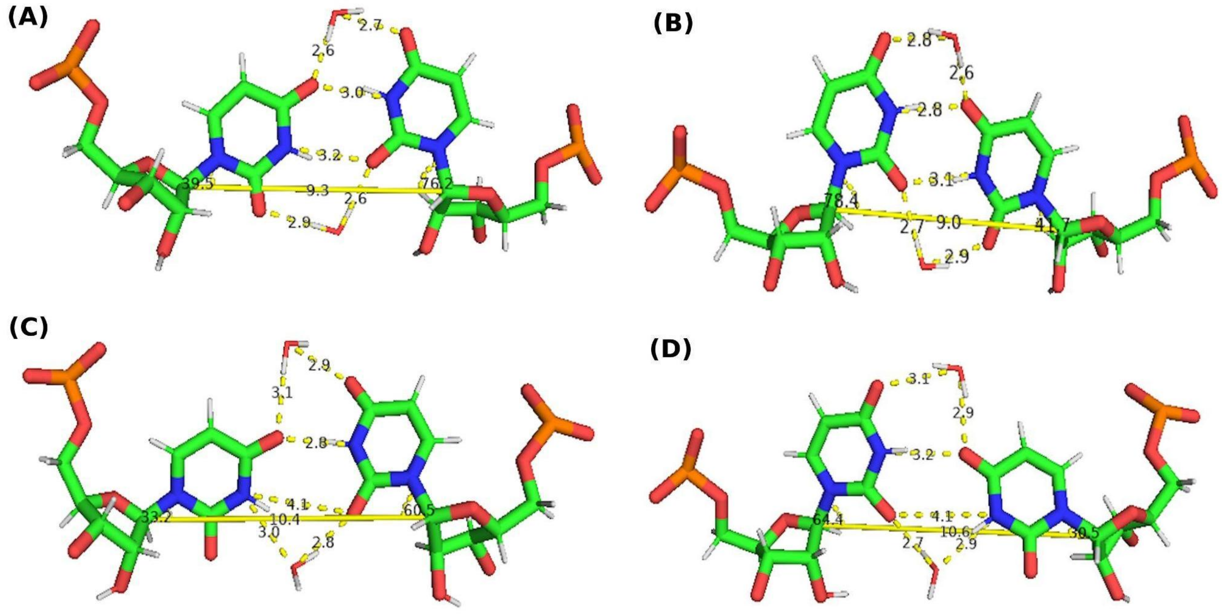

**Figure S22.** Snapshots of (A,B) Symmetric hydrogen bonded U-U wobble base pair with water mediated interactions (O4(5)-W-O4(14) and O2(14)-W-O2(5)) by two key water molecules, other than the two direct hydrogen bonding interactions, respectively for the two geometries observed in this study and (C,D) U-U 'stretched' wobble base pair with water mediated interactions (O4(5)-W-O4(14) and O2(14)-W-N3(5) (geometry 1) or O4(5)-W-O4(14) and O2(5)-W-N3(14) (geometry 2)) by two key water molecules other than one direct hydrogen bond, for the two geometries observed in this study (corresponding to Duplex-GUC (U-U)). Distances between hydrogen bond acceptor and donor atoms are shown with yellow dotted lines. The C1'-C1' distances and λ (C1'-C1'-N1 for U) angles are also shown.

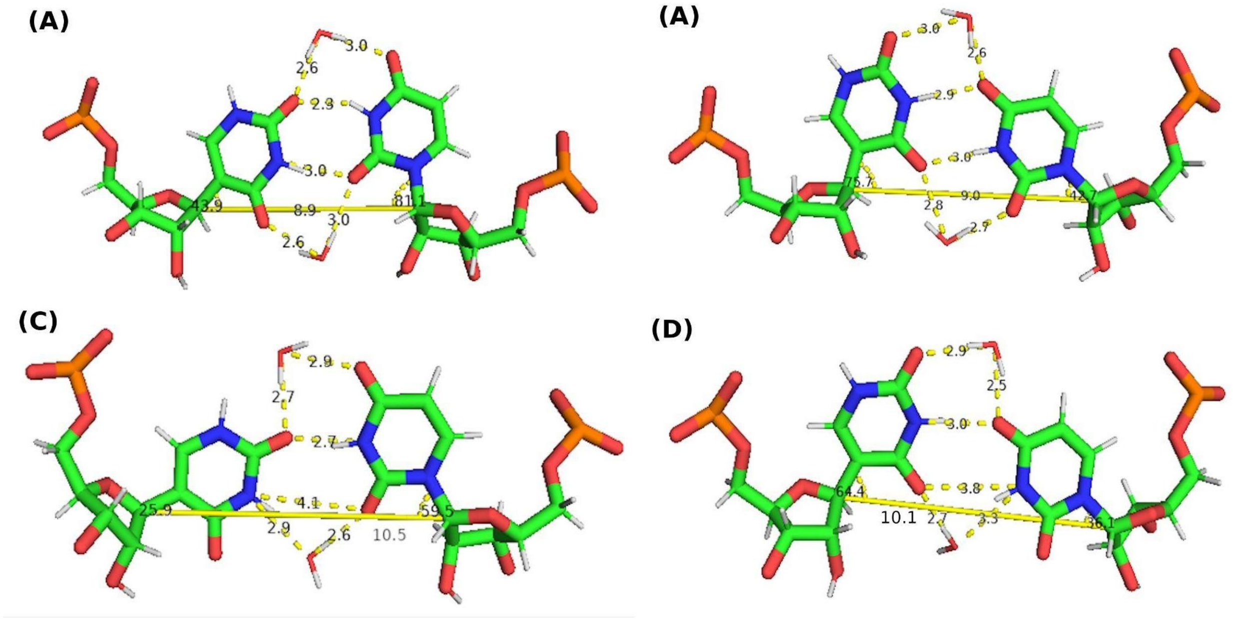

**Figure S23.** Snapshots of (A,B) Symmetric hydrogen bonded Ψ-U wobble base pair with water mediated interactions (O4(5)-W-O2(14) and O4(14)-W-O2(5)) by two key water molecules, other than the two direct hydrogen bonding interactions, respectively for the two geometries observed in this study and (C,D) Ψ-U 'stretched' wobble base pair with water mediated interactions (O4(14)-W-O2(5) and O2(14)-W-N3(5) (geometry 1) or O4(14)-W-O2(5) and O4(5)-W-N3(14) (geometry 2)) by two key water molecules other than one direct hydrogen bond, respectively for the two geometries observed in this study (corresponding to Duplex-GΨC (Ψ-U)). Distances between hydrogen bond acceptor and donor atoms are shown with yellow dotted lines. The C1'-C1' distances and λ (C1'-C1'-C5 for Ψ and C1'-C1'-N1 for U) angles are also shown.

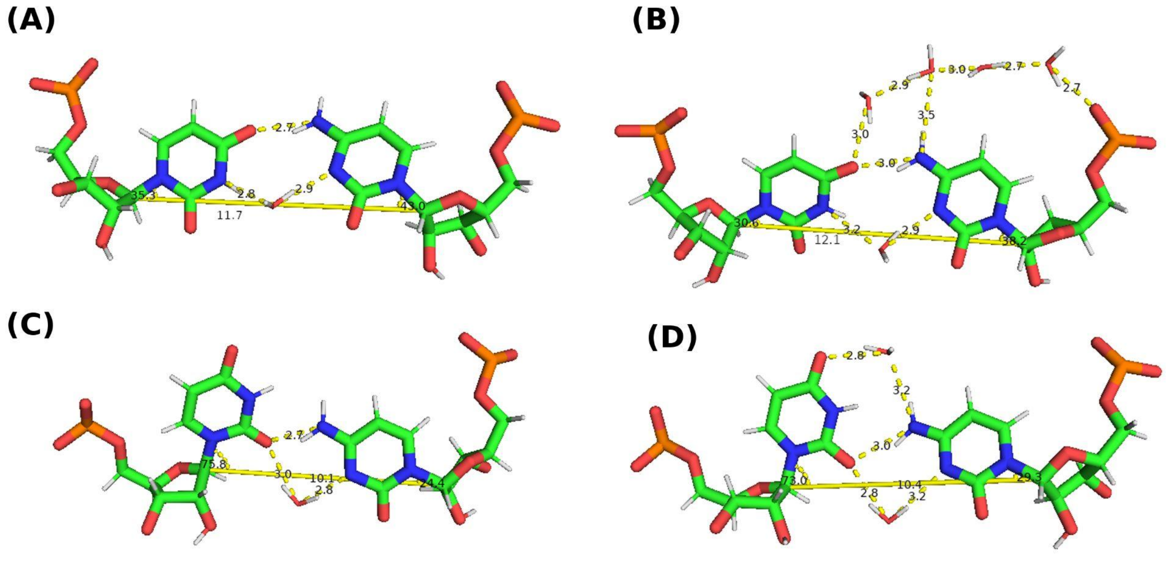

**Figure S24.** Snapshots of U-C pair (corresponding to Duplex-GUC (U-C)) with water mediated interactions by (A) one key water molecule (N3(5)-W-N3(14)) (B) by 5 key water molecules (N3(5)-W-N3(14) and four water molecules connecting O4(5), N4(14) and phosphate oxygen of C14) other than the one direct hydrogen bonding interaction for geometry 1; (C,D) by one key water molecule (O2(5)-W-N3(14)) or by two key water molecules (O2(5)-W-N3(14) and (O4(5)-W-N4(14)) other than one direct hydrogen bond for geometry 2. Distances between hydrogen bond acceptor and donor atoms are shown with yellow dotted lines. The C1’-C1’ distances and λ (C1’-C1’-N1 for U and C) angles are also shown.

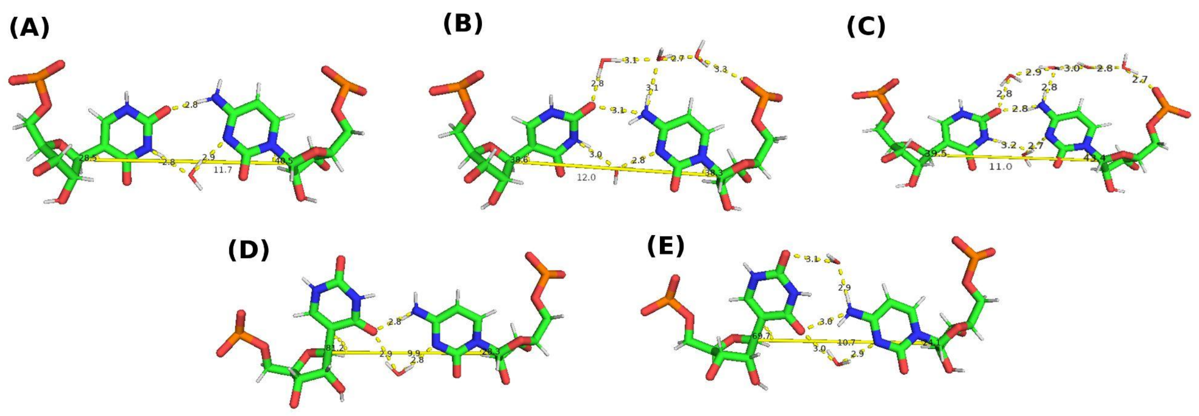

**Figure S25.** Snapshots of Ψ-C pair (corresponding to Duplex-GΨC (Ψ-C)) with water mediated interactions by (A) one key water molecule (N3(5)-W-N3(14)) (B) by 4 key water molecules (N3(5)-W-N3(14) and three water molecules connecting O2(5), N4(14) and phosphate oxygen of C14) and (C) by 5 key water molecules (N3(5)-W-N3(14) and 4 water molecules connecting O2(5), N4(14) and phosphate oxygen of C14), other than the one direct hydrogen bonding interaction for geometry 1, (D,E) by one key water molecule (O4(5)-W-N3(14)) or by two key water molecules (O4(5)-W-N3(14) and (O2(5)-W-N4(14)) other than one direct hydrogen bond for geometry 2. Distances between hydrogen bond acceptor and donor atoms are shown with yellow dotted lines. The C1'-C1' distances and λ ( C1'-C1'-C5 for Ψ and C1'-C1'-N1 for C) angles are also shown.

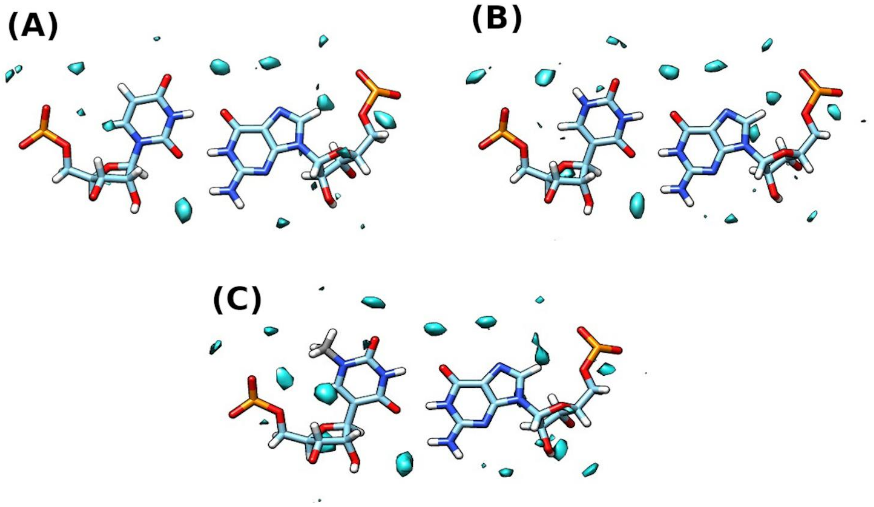

**Figure S26.** Water occupancy maps for (A) U-G; (B) Ψ-G (C) $m^1$Ψ-G pairs respectively (corresponding to the last 400 ns (40000 conformations) of the trajectory)

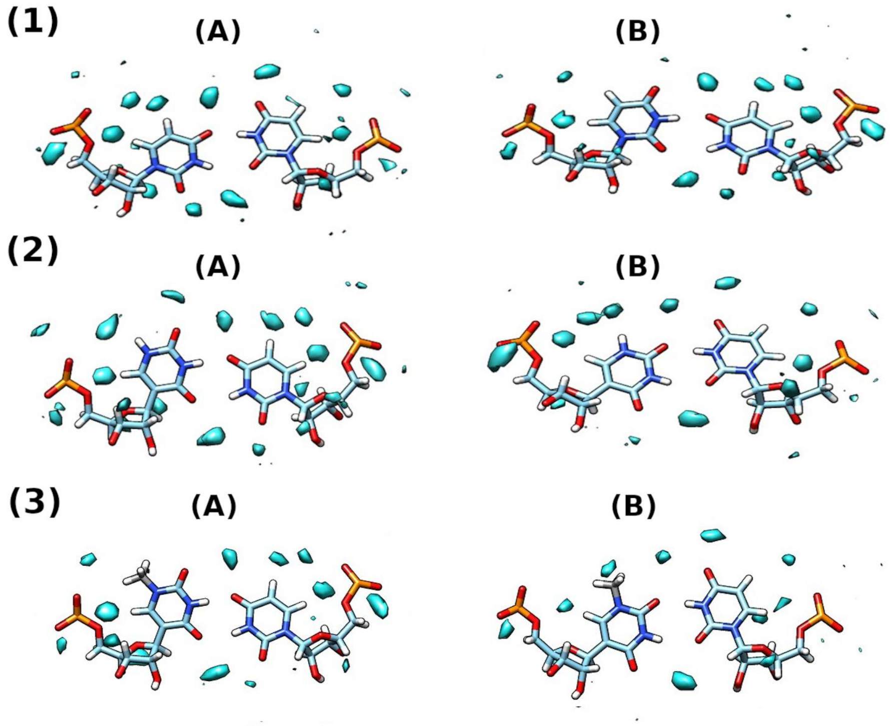

**Figure S27.** Water occupancy maps for (1) U-U; (B) Ψ-U (C) m[1]Ψ-U pairs respectively (A) and (B) corresponds to the the two geometries (for the most populated and the second most populated clusters respectively) observed for these pairs (corresponding to the 10000 conformations from each cluster)

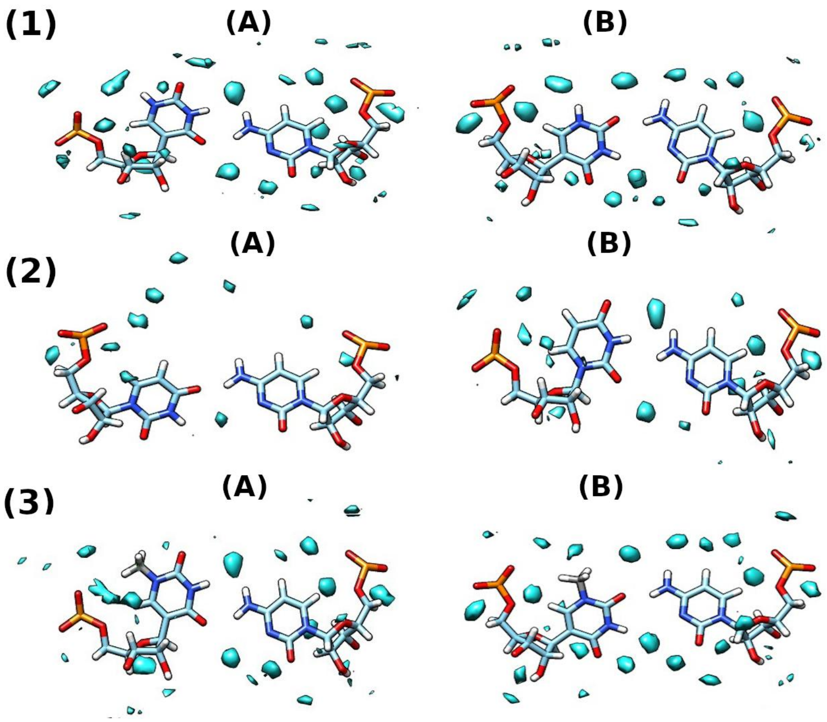

**Figure S28.** Water occupancy maps for (1) U-C; (B) Ψ-C (C) m[1]Ψ-C pairs respectively (A) and (B) corresponds to the the two geometries (for the most populated and the second most populated clusters respectively) observed for these pairs (corresponding to the 10000 conformations from each cluster)

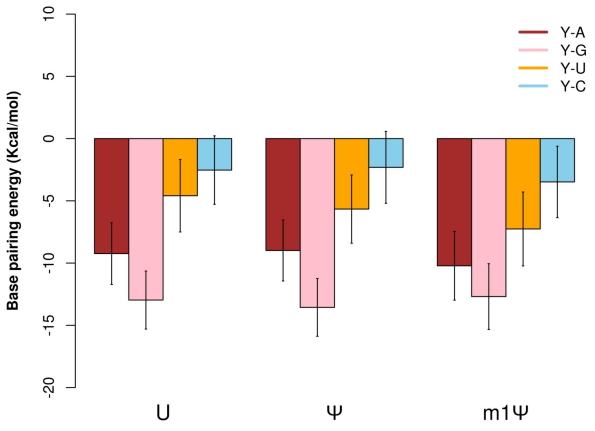

**Figure S29.** Comparison of the base pairing energies corresponding to the base pairs of Y (U/Ψ/m[1]Ψ) with A, G, U and C respectively obtained from MD simulations (the values corresponding to the first set of simulation for m[1]Ψ-modified duplexes are plotted).

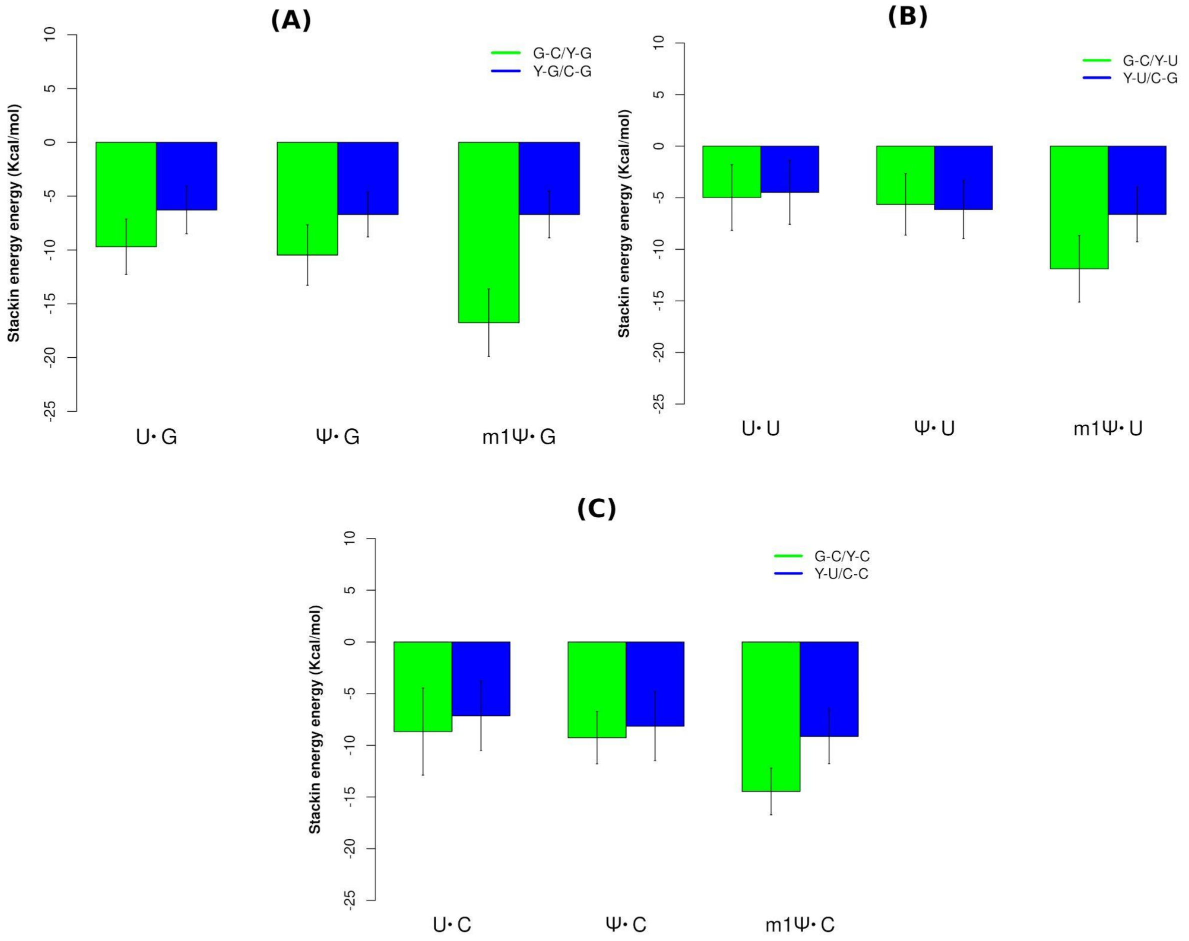

**Figure S30.** Comparison of the base pair stacking energies of the (A) base pair steps containing U-G, Ψ-G and $m^1\Psi$-G pairs respectively (B) base pair steps containing U-U, Ψ-U and $m^1\Psi$-U pairs respectively and (C) base pair steps containing U-C, Ψ-C and $m^1\Psi$-C pairs respectively obtained from MD simulations (the values corresponding to the first set of simulation for $m^1\Psi$-modified duplexes are plotted). Y in the figure legend indicates U/Ψ/$m^1\Psi$.
